# GRAViTy-V2: a grounded viral taxonomy application

**DOI:** 10.1101/2024.07.26.605250

**Authors:** Richard Mayne, Pakorn Aiewsakun, Dann Turner, Evelien M. Adriaenssens, Peter Simmonds

**Affiliations:** Peter Medawar Building for Pathogen Research, Nuffield Department of Medicine, University of Oxford, 3 South Parks Road, OX1 3SY, Oxfordshire, United Kingdom; Department of Microbiology, Faculty of Science, Mahidol University, Mahidol University, 73170, Nakhon Pathom, Thailand; School of Applied Sciences, University of the West of England, Frenchay Campus, BS16 1QY, Bristol, United Kingdom; Quadram Institute Bioscience, Rosalind Franklin Rd, NR4 7UQ, Norwich, United Kingdom

**Keywords:** Bioinformatics, Viral Taxonomy, Classification, Genomic Analysis

## Abstract

Taxonomic classification of viruses is essential for understanding their evolution and therefore their distribution, host interactions and pathogenic mechanisms. Classification methodologies usually rely on comparison of aligned sequence motifs in conserved genes, by genome organisation and gene complements, and at lower taxonomic ranks such as genus and species, through genome sequence identities. Building on our previous classification framework based on a novel whole-genome analysis method, we here describe Genome Relationships Applied to Viral Taxonomy Version 2 (GRAViTy-V2), which encompasses a greatly expanded range of features and numerous optimisations, packaged as an application that may be used as an alignment-free general-purpose virus classification tool. Using 28 datasets derived from the International Society on Taxonomy of Viruses 2022 taxonomy proposals, GRAViTy-V2 output was compared against human expert-curated classifications used for assignments in the 2023 round of ICTV taxonomy changes. GRAViTy-V2 produced taxonomies equivalent to manually-curated versions down to the family level and in almost all cases, to genus and species levels. However, discrepancies with our results primarily arose through various human and automated sequence annotation errors and erroneous annotations of coding sequences used in their original classification. Analysis times ranged from 1–506 min (median 3.59) on datasets with 17–1004 genomes and mean genome length of 3,000–1,000,000 bases, on a standard consumer-grade laptop. We discuss how the output from GRAViTY-V2 outputs allows for a full analysis of why taxonomic classifications were proposed, the value of the program for quality control of genetic comparisons, and how to optimise the speed of classification through proper use of GRAViTy-V2’s workflow management system.

## Introduction

Taxonomic classification of viruses is essential for understanding their evolution, distribution, host interactions and pathogenic mechanisms. Historically, attributes such as virion structure, pathogenicity in their hosts, replication mechanisms and epidemiology / transmission routes have been used to define virus taxa, typically classifying them into order, families, genera and species, analogous to the taxonomy of cellular life forms. However, the recent application of high throughput sequencing technologies, and the application to aquatic, terrestrial and gut microbiome samples has revealed an astonishing diversity of viruses infecting prokaryotes and a range of eukaryotes, including amoebae, algae, insects, fish and plants well beyond human, veterinary and crop plant hosts that have been typically investigated in previous decades [1, 2, 3, 4].

To ensure that virus taxonomy better captures the true diversity of viruses, the International Committee on Taxonomy of Viruses (ICTV), on consultation with the wider virology community, agreed to include and formally classify viruses known only by their nucleotide sequences [5]. More recently, the principles behind a purely evolutionarily based classification of viruses have been proposed based on genomic relatedness [6], albeit acknowledging the value of additional information, where available, on structural, biological and morphological characteristics, in defining the taxonomic grouping boundaries upon which these virus relationships are assigned. While these fundamental principles for classification are widely agreed [6], how sequences may be analysed and relationships determined remains highly variable between virus groups. The existence of at least six and likely many more independent origins of viruses [7], and the varying use of different sets of hallmark genes to reconstruct evolutionary relationships within each realm [8], does not provide a common framework for taxonomy equivalent, for example, to the use of ribosomal gene sequence comparisons to reconstruct the deeper evolution of cellular organisms. Consequently, taxonomic assignments in different virus groups may be based on sequence comparisons or, at deeper taxonomic levels, the detection of protein structural homologies, of quite different gene sets subjectively chosen by researchers. All of this requires expert curation, effective sequence alignment methodologies for often highly divergent virus sequences and expertise in protein structure prediction [9]. There consequently exists no simple, agnostic approach for investigating virus relationships, even at lower taxonomic levels such as order, family and genus.

A further challenge that frustrates efforts to classify viruses relates to tractability of computation. Since the advent of metagenomic sequencing, the rate of new virus discovery has increased exponentially [10]. Viral genomes are comparatively small, but calculating similarity between them based on the results of alignments — once a mainstay through tools such as PASC [11] — quickly becomes infeasible when hundreds of virus genomes are compared. Although more efficient algorithms continue to be developed, the current gold standard instead focuses on generating alignments between short, conserved genetic sequences (motifs), such as RNA-dependent RNA polymerases (RdRp) in RNA viruses [6]. Techniques for better detecting homology when sequences are highly diverse, such as abstracting DNA/RNA as translated amino acid sequences, are also routinely used, but this simple task can still become intractable at scale. Motif-based methods furthermore disregard homologies between other genomic regions that cannot be detected across all viral groups under the analysis, but which may still be detectable among some subgroups and are taxonomically relevant for those subgroups.

In previous work [12, 13], we described and evaluated ‘GRAViTy’ (Genome Relationships Applied to Virus Taxonomy), a framework for identifying and classifying viruses in which a single metric representing a comparison of genes, their relative locations, orders and orientations is computed to generate a taxonomy. Initial iterations of the framework were, however, computationally expensive, not packaged as a discrete software and insensitive in specific conditions, such as comparison of very long viral genomes and computing classifications for datasets with a large proportion of unclassified sequences.

In this article we present ‘GRAViTy-V2’, which implements a comprehensively-updated, expanded and optimised framework as a standalone, user-friendly application. This implementation allows one or more virus sequences (new or directly from the published accession numbers of complete genomes) to be compared with viruses classified in the latest release of the ICTV taxonomy. Through computation of a composite Jaccard score, overall genomic similarity to existing taxa can be quantified, and visualised in the form of heatmaps and dendrograms. The new version further introduces a range of ‘explainability’ features for describing how classifications were arrived at and hence support a wider range of taxonomy tasks.

The use of ‘grounded’ in this article’s title is derived from Computational Grounded Theory, which was used as inspiration for the design of our software and holds that more methodologically rigorous, interpretative approaches to content analysis are derived from a combination of computational pattern recognition and expert human knowledge [14].

We present here a comprehensive evaluation of GRAViTy-V2, in which we analysed sequence data from all accepted and ratified ICTV taxonomy proposals (TPs) from 2022 and compared results with expert-curated taxonomies. We demonstrate how our software may effectively complement the current gold standard of human curation and allow virologists additional insights into taxonomic classification based on metrics of genetic similarity and relative genomic location of the constituent genetic components in viral genomes. We conclude by discussing both the applications and limitations of the software, and contrast GRAViTy-V2 with several other widely-used and novel tools.

## Materials & Methods

### Software

GRAViTy-V2 is written in Python 3.10 and is compatible with bash-like environments. It is distributed as an open-source software package, with installation and operating instructions, via GitHub (https://github.com/Mayne941/gravity2) with a GPL 3.0 license. It may be unpacked and installed, along with all dependencies, on Debian-based operating systems using a packaged installer bash script. The application runs on a local Uvicorn server as a RESTful API, composed in FastAPI [15]. Users may interact with GRAViTy-V2 either through a browser-based graphical user interface (SI Document 1, S1), or by cURL’ing its endpoints from the command line or appropriate third-party programs. GRAViTy-V2 may also be interacted with via a command line interface (CLI), which may be preferable for expert users when run on shared resources. It is designed to be compatible with personal computers but may also be deployed on HPC or cloud infrastructure.

### Algorithm

The GRAViTy-V2 framework is a condensed version of the first, 2018 release in which the two pipelines (comparison of reference and unclassified genomes, respectively) have been merged (Fig. 1). Input data requirements are a list of reference sequences and, optionally, additional unclassified sequences: GRAViTy-V2 may be used to evaluate existing classifications, classify new sequences, or a combination of the two. ICTV-curated Virus Metadata Resource (VMR) lists complete genome sequences of taxonomically classified viruses and provides optimal data for building reference databases. GRAViTy-V2 utility functions accordingly are used to scrape the latest VMR, extract a subset of taxonomically-relevant viruses (either user-selected or automatically-generated) and generate a database of sequences against which unclassified sequences may be compared. Data input format requirements are either a VMR-like document containing accession IDs, multi-FASTA files, or a combination of both.

**Fig. 1.**
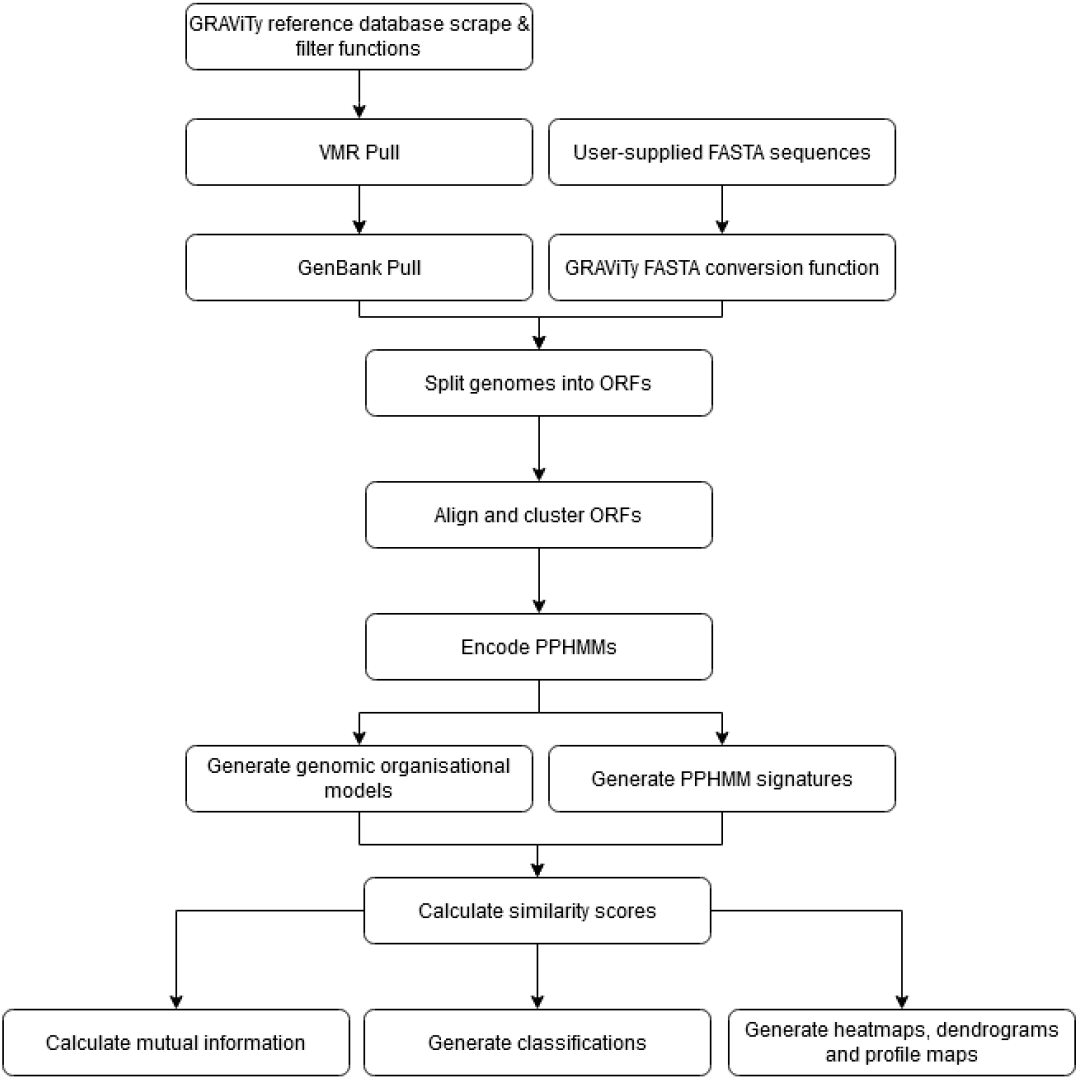
GRAViTy-V2 process flow.

Classification is performed by empirically detecting and extracting open reading frames (ORFs) of lengths exceeding a user-set threshold (default 100 amino acids), from coding complete viral genomes. Where the first version would use INDSC sequence annotations to identify coding-complete sequences where available, the current version always opts for empirical ORF detection, firstly to provide data more comparable to that derived from the typically unannotated genome sequences of newly discovered viruses, and secondly, to avoid problems arising from potential errors in sequence annotations.

ORFs are translated, aligned, clustered and converted to protein profile hidden Markov models (PPHMMs). PPHMMs are assembled into a database from which similarities between profiles (PPHMM signatures) and their genomic positions and orientations (Genomic Organisational Models, GOMs) are computed. These scores are then combined by calculating the geometric mean of generalised composite Jaccard (GCJ) (intersection over union) similarity scores: 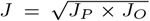, where *J*_*P*_ is the similarity score for the model’s PPHMM component and *J*_*O*_ the GOM [13]. All classifications are supported by statistical analysis of feature importance, using mutual information score implemented in Python’s Scikit-learn library [16]. Another algorithm modification in version 2 is the ‘multiple pass system’, whereby searches through very large data spaces are made more efficient by omitting unnecessary information in initial ‘passes’ through a dataset. For example, where a user may have an unclassified viral genome and has no clear idea of its taxonomy, they may run a ‘first pass’ which compares their unknown sequences against representative single genomes (either user-selected or automatically generated by a GRAViTy-V2 utility function) from each family within the realm they are searching, wherein reference sequences may either be selected manually or automatically through a GRAViTy-V2 utility function. This provides an approximate taxonomy and allows the user to conduct a more granular ‘second pass’ run, using a greater quantity of reference genomes from clades most closely related (by GCJ score) to the unclassified genome. Selection of second pass reference genomes is currently a manual process as a user’s rules for inclusion will vary, but endpoints are available through both graphical user interface (GUI) and CLI for filtering input VMRs to user’s selection of taxa. If applied to the realm *Riboviria*, a two pass strategy reduces the search space from approximately 5 ×10^3^ genomes on the first pass to 117 on the second pass and results in an approximately ten-fold reduction in compute time.

### Experimental Evaluation

To evaluate GRAViTy-V2, a total of 28 taxonomy proposals (TPs) were analysed, of which 21 introduced at least one new species, 6 assigned new genera, 4 new families and 2 taxon reorganisations (Table 1), where new genera/family operations involved creation of entirely new taxa comprised only of members of new sequences. All these proposals had been subject to manual classification by by expert members of relevant ICTV study groups. TPs were sourced from ICTV archives (https://ictv.global/files/proposals/approved), all of which contained both tabular and graphical summaries (phylogenetic trees and in some cases, pairwise distance matrices) of proposed taxonomies.

**Table 1.** Dataset descriptions and run times. In cases where multiple passes were used (*†*), combined times are shown. Grouping level: taxon at which VMR of reference genomes was filtered; N: number of genomes; S: new species operation; G: new genus operation; F: new family operation; RT: reorganise taxon operation.

| Taxon | ICTV Code | Grouping Level | N | Mean Len (bases) | Time (mins) | Operation |
| --- | --- | --- | --- | --- | --- | --- |
| *Jingchuvirales* | 2021.015M | *Jingchuvirales* | 58 | $9.75 \times 10^3$ | 1.93 | 2G, 10S |
| *Alteviridae* | 2022.001F | *Orthornavirae* | 62 | $3.31 \times 10^3$ | 1.95 | 1F |
| *Picornavirales*† | 2022.001S | *Orthornavirae* | 1004 | $5.72 \times 10^3$ | 5.61 | 1F |
| *Herpesvirales*† | 2022.002D | *Herpesvirales* | 112 | $1.69 \times 10^5$ | 506 | 9S |
| *Bacilladnaviridae* | 2022.002F | *Arfiviricetes* | 287 | $1.93 \times 10^3$ | 3.93 | RT |
| *Alpharhabdovirinae* | 2022.002M | *Rhabdoviridae* | 273 | $1.19 \times 10^4$ | 19.9 | 1G, 14S |
| *Pestivirus* | 2022.002S | *Pestivirus* | 21 | $1.24 \times 10^4$ | 0.72 | 8S |
| *Botourmiaviridae* | 2022.003F | *Botourmiaviridae* | 154 | $2.73 \times 10^3$ | 1.70 | 15S |
| *Arenaviridae* | 2022.003M | *Arenaviridae* | 63 | $5.15 \times 10^3$ | 1.83 | 1G, 1S |
| *Baculoviridae* | 2022.004D | *Baculoviridae* | 100 | $1.34 \times 10^5$ | 110 | 6S |
| *Imitervirales* | 2022.004F | *Imitervirales* | 22 | $1.51 \times 10^6$ | 150 | RT |
| *Parvoviridae* | 2022.005D | *Parvoviridae* | 172 | $5.2 \times 10^3$ | 3.26 | 2G, 49S |
| *Mammonoviridae* | 2022.005F | *Nucleocytoviricota* | 41 | $2.42 \times 10^5$ | 61.3 | 1S |
| *Cripavirus* | 2022.006S | *Dicistroviridae* | 17 | $9.4 \times 10^3$ | 0.40 | 1S |
| *Cytorhabdovirus* | 2022.007M | *Rhabdoviridae* | 277 | $1.19 \times 10^4$ | 20.9 | 10S |
| *Ephemerovirus* | 2022.008M | *Rhabdoviridae* | 269 | $1.18 \times 10^4$ | 19.3 | 2S |
| *Fraservirus*† | 2022.010M | *Riboviria* | 118 | $5.94 \times 10^3$ | 5.02 | 1F |
| *Hartmanivirus* | 2022.001M | *Arenaviridae* | 64 | $5.18 \times 10^3$ | 1.80 | 2S |
| *Lispiviridae* | 2022.012M | *Lispiviridae* | 30 | $1.20 \times 10^4$ | 1.01 | 7G, 11S |
| *Mammarenavirus* | 2022.013M | *Arenaviridae* | 74 | $5.67 \times 10^3$ | 2.07 | 1S |
| *Myonnaviridae* | 2022.014M | *Myonnaviridae* | 31 | $8.94 \times 10^3$ | 0.78 | 13S |
| *Nyamiviridae* | 2022.016M | *Mononegavirales* | 21 | $1.02 \times 10^4$ | 1.42 | 2S |
| *Orthobunyavirus* | 2022.017M | *Peribunyaviridae* | 143 | $4.11 \times 10^3$ | 4.65 | 1S |
| *Orthobunyavirus* | 2022.018M | *Peribunyaviridae* | 171 | $4.64 \times 10^3$ | 5.78 | 29S, RT |
| *Phasmaviridae* | 2022.019M | *Phasmaviridae* | 29 | $4.18 \times 10^3$ | 0.98 | 3S |
| *Phenuiviridae* | 2022.020M | *Phenuiviridae* | 142 | $4.17 \times 10^3$ | 5.72 | 2G, 10S |
| *Varicosavirus* | 2022.021M | *Rhabdoviridae* | 276 | $1.16 \times 10^4$ | 2.55 | 9S |
| *Vesiculovirus* | 2022.022M | *Rhabdoviridae* | 269 | $1.18 \times 10^4$ | 9.13 | 2S |

All sequence data were downloaded automatically from GenBank via GRAViTy-V2 and where multipartite genomes were included, these were assembled by default in largest-to-smallest order although this can be user-specified. Experiments were conducted in a single pass except for new family operations, in which cases a two-pass methodology was adopted: representative single genomes from each family within the relevant realm were picked (automatically by the utility function) as reference viruses during the first pass, which was used to identify the closest three families. For the second pass, every species from these closest three families were used as reference viruses. Default GRAViTy-V2 parameters were used in the first instance for every dataset, but several were re-run with refined parameters.

Classifications predicted by GRAViTy-V2 were manually compared with curated taxonomies through recording the number of bootstrap supported phylogeny violations between GRAViTy- and ICTV-generated classifications, as interpreted from tabular summaries and comparison with maximum likelihood (ML) phylogenetic trees at family and genus levels, for both existing taxa and for proposed new or reorganised taxa. All violations were investigated to determine their cause, for which genome length, shared profile ratios and profile positions were used, all derived from GRAViTy-V2 output.

### Hardware

All experiments were conducted on a consumer-grade laptop with an Intel i9-11980HK processor (3.30 GHz, 8 core + 8 threads) and 32 Gb DDR4 RAM (3200 Mhz), via Windows Subsystems Linux (WSL) 2 running Ubuntu 22.04. Run times were benchmarked in all experiments.

## Results

### Software modifications

Key software modifications were:

1. Packaging framework as a multi-operating system application behind an application programming interface (API), with GUI, CLI entrypoints, installer script and extensive error handling.
2. Refactored codebase to run entire workflow from a single pipeline that supports ‘fire and forget’ triggering of single and batch jobs.
3. Implementation of a multi-stage workflow, which supports multiple ‘passes’ through reference datasets at different levels of granularity, which reduces run time by removing unnecessary calculations.
4. Extensive optimizations, including rewriting the application in Python 3.10, updating dependencies to latest versions and redesign of compute-intensive functions to memory efficient, parallelised equivalents.
5. New algorithm for extracting, assembling and calculating protein profile locations for ORFs, in a scalable manner that supports accurate comparison of multipartite genomes.
6. New output statistics and figures for explaining why the software has made a classification, including maps of protein profile locations within genomes, distance of protein profiles from mean location, and shared profile ratios normalised to genome length.
7. Development of a more sensitive similarity matrix scoring scheme that considers ratio of shared protein profiles, which may be combined with existing similarity schemes.
8. Support for use of Mash scores [17] rather than BLASTp for initial distance estimation, which can reduce run times and enhance sensitivity in certain conditions.
9. New utility functions to support taxonomic investigations, including FASTA to GRAViTy-V2-compatible input converter, multipartite genome assembler and web scraper function to gather the latest VMR version from the ICTV website.
10. Creation of several ‘premade’ pipelines with parameter configurations optimised for specific scenarios (e.g. comparing highly divergent viruses), for which users only need specify three parameters.
11. The option to export contiguous blocks of sequence in regions where PPHMM matches can be detected. The identification and extraction of sequences (e.g. RdRP and helicase regions of RNA viruses) greatly assists parallel analyses of virus relationships though conventional alignment and phylogeny methods.

### Run time benchmarking

Run times for each TP were benchmarked against mean genome size and the number of genomes compared (Table 1), and varied from 1–506 min (median 3.59) across datasets of size 21–1004 genomes. Time scaling increased with both mean genome size and number of pairwise comparisons (Fig. 2). The longest run time outlier was for the *Herpesvirales* dataset, which was notable for it containing a large number of comparatively long genomes, for which two passes were run. Experiments completed 50–70% faster than equivalent runs of the previous GRAViTy version (data not shown) and system requirements are now dramatically lower, such that HPC infrastructure was not required for our use case. Total compute time across all experiments was approximately 16 hours and was scripted to run overnight.

**Fig. 2.**
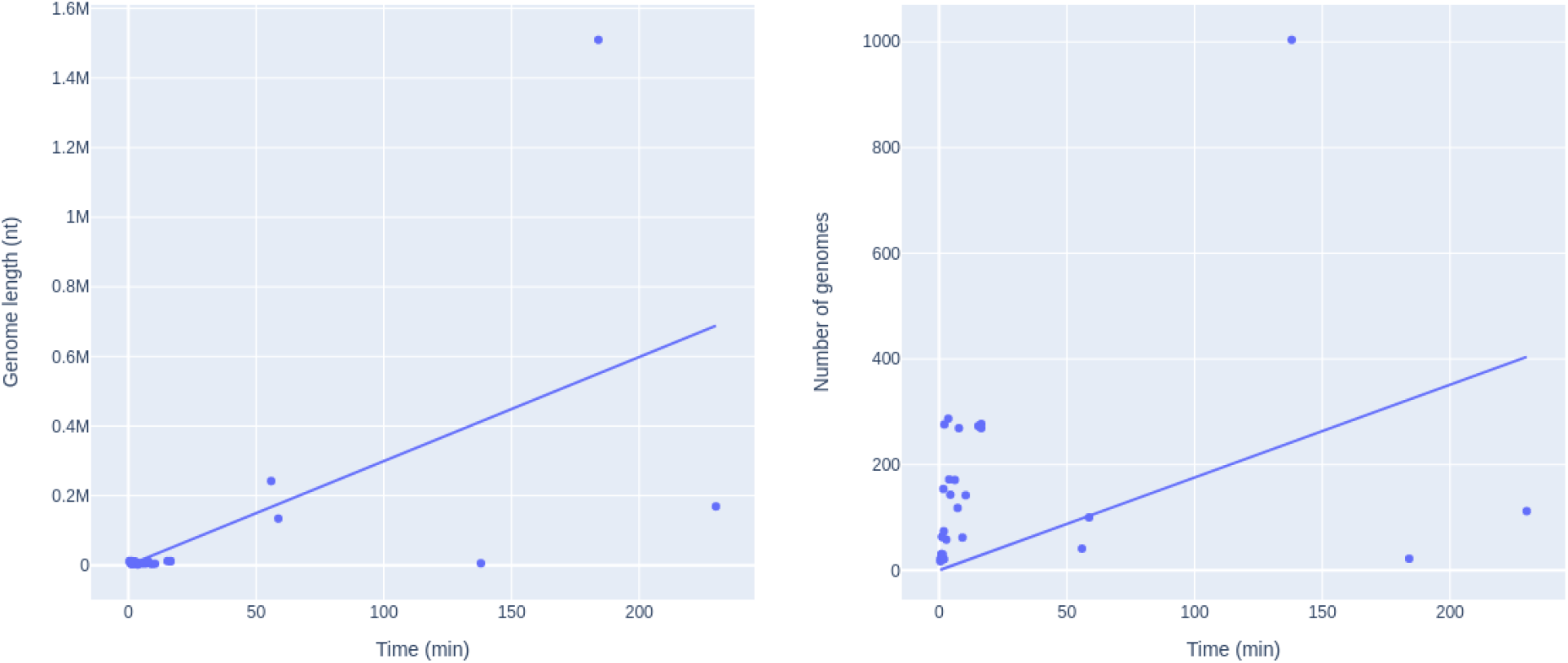
GRAViTy-V2 Run time benchmarks of against mean genome length and number of genomes, with linear regression lines showing positive trend (forced intercept at origin).

### GRAViTy-V2 graphical outputs

GRAViTy-V2 graphical outputs include a pairwise composite genomic similarity scores accompanied by a bootstrapped dendrogram ‘GRAViTy-V2 Heatmap’ (Fig. 3c); ‘Shared normalised PPHMM ratio matrix’, pairwise comparison of quantity of shared PPHMMs, normalised to the number of profiles assigned to each genome (Fig. 4a); ‘Barcode’ heatmap, genomic position of each PPHMM midpoint, normalised against genome length (heatmap colour) verses median position among all genomes (X-axis) (Fig. 4b). Full graphical output from all experiments are included (SI Document 2).

**Fig. 3.**
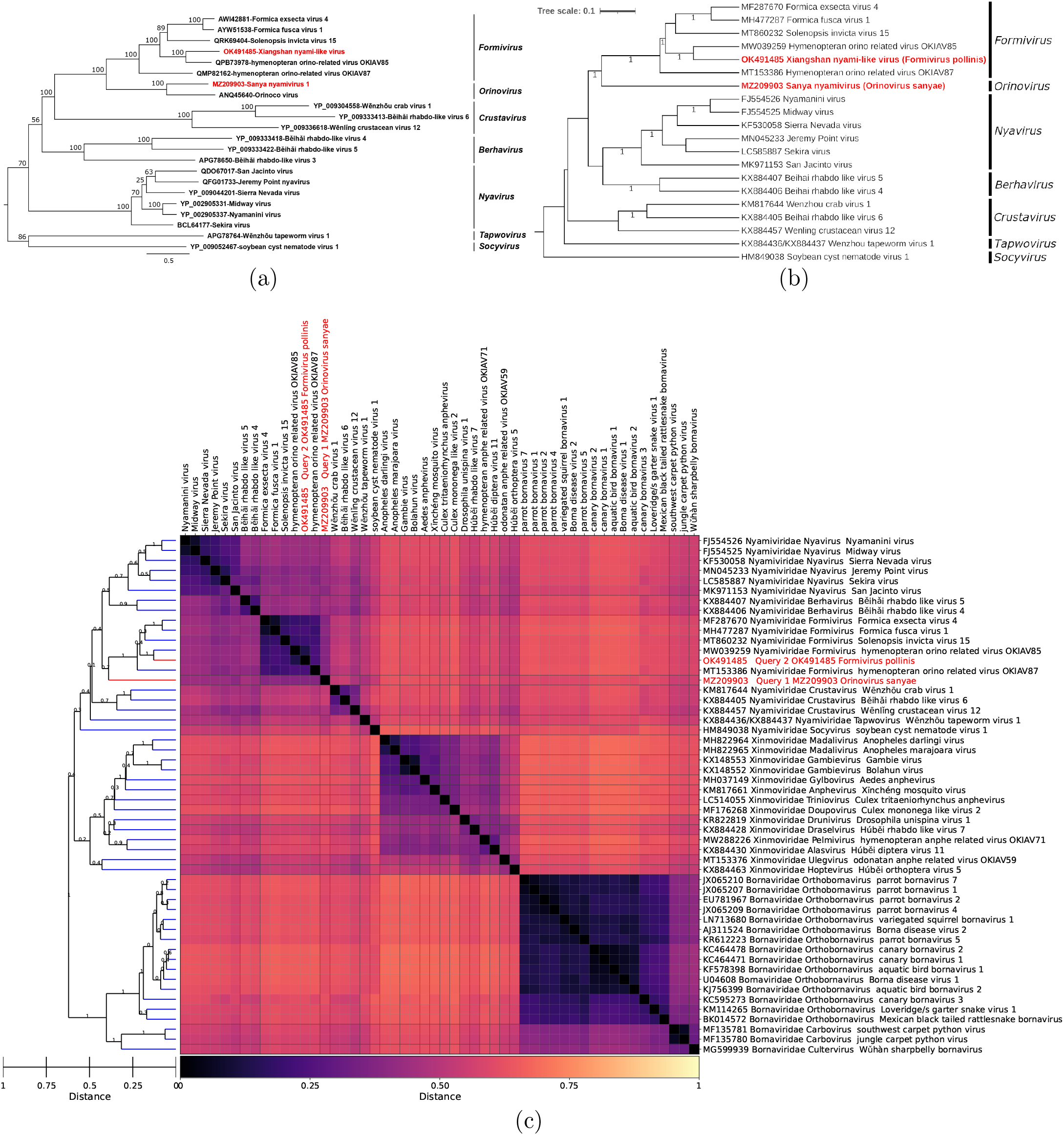
*Nyamiviridae* TP. (a) TP ML tree, from L protein AA alignment [18]. (b) GRAViTy-V2 tree (Orinocovirus and Beihai rhabdo-like virus 3 omitted as genomes non-coding complete). (c) GRAViTy-V2 heatmap, including neighbouring families *Xinmoviridae* and *Bornaviridae*. (Red text: sequences proposed as new taxa in TP; bootstrap values *<* 0.7 hidden).

**Fig. 4.**
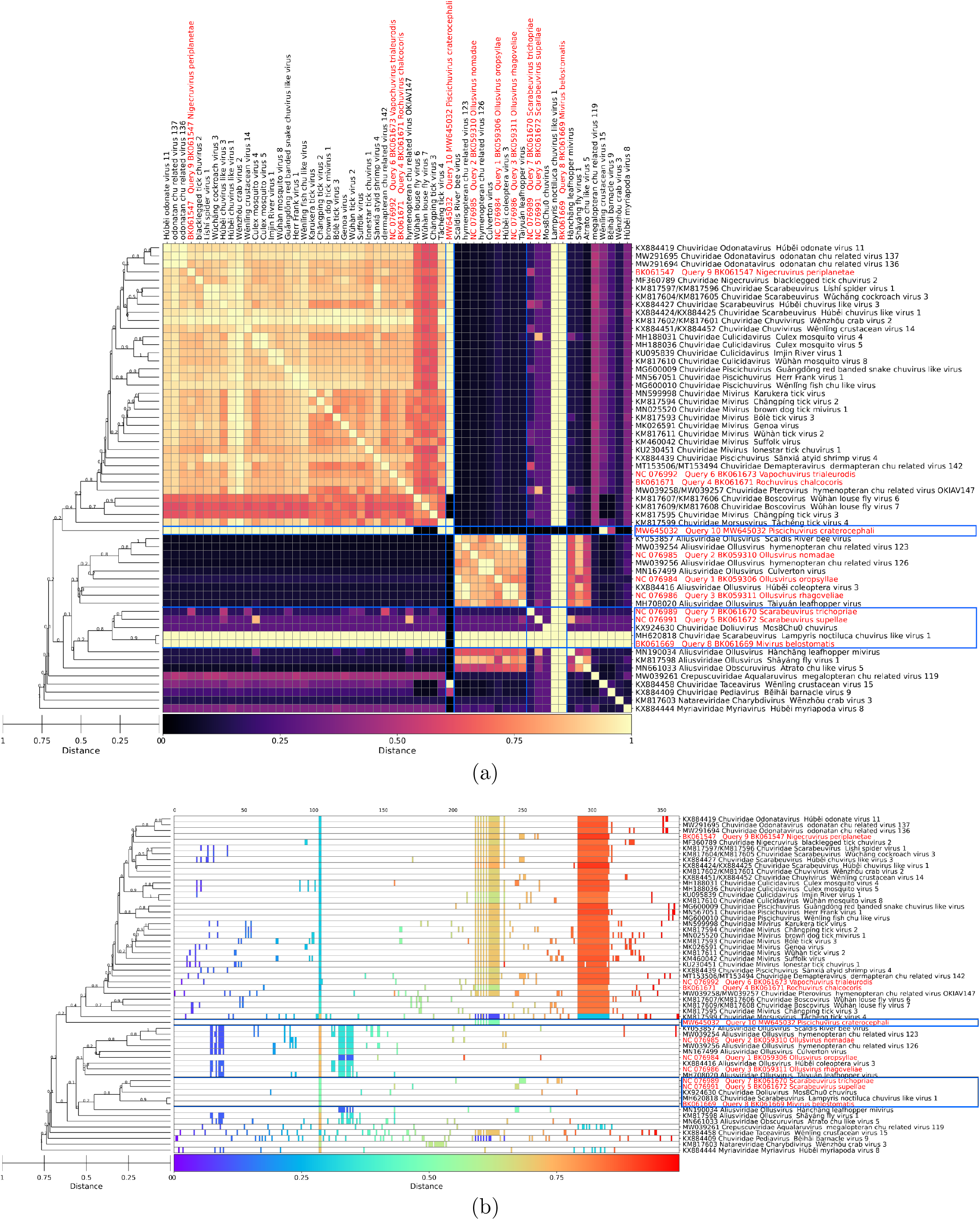
*Jingchuvirales* TP. (a) Shared normalised PPHMM ratio matrix. (b) Barcode heatmap. (Blue lines: incomplete sequences; red text: sequences proposed as new taxa in TP)

### Comparison between GRAViTy-V2 results and expert-curated taxonomies

Overall, GRAViTy-V2 output was highly consistent with expert-curated taxonomies, with dendrograms and heatmaps based on composite Jaccard scores reproducing sequence identities and patterns of phylogenetic genetic clustering determined by the ICTV study groups. Remarkably, this concordance was consistent across the diverse phylogenetic methods used by the study groups, which ranged from simple alignments of single conserved regions of short RNA viruses such as *Phasmaviridae* (Fig. 5) to complex, curated alignments of multiple marker genes from DNA viruses such as in the *Mamonoviridae* TP, which included genomes from the largest known viruses. Overall, three quarters of classifications (22/28) exhibited no family or genus-level violations between ICTV ratified taxonomic classifications and relationships determined by GRAViTy-V2 (Table 2).

**Table 2.** Results of GRAViTy-V2 TP analysis compared to curated classifications, with original TP taxonomic methods (tools listed where information provided in TP). N: Number of species; NF*{*/V*}*: Number of families / family-level violations compared to corresponding TP; NG*{*V*}*: Number of genera/ genus-level violations; CDS: (protein) coding sequence; ML: maximum likelihood.

| Taxon | N | NF | NFV | NG | NGV | TP Method | Violations |
| --- | --- | --- | --- | --- | --- | --- | --- |
| *Jingchuvirales* | 58 | 5 | 5 | 21 | 2 | ML tree (PhyML) from RdRp alignment (ClustalW/Muscle, with manual curation). | 6 incomplete genomes (BK061669, BK061670, BK061672, MW645032, MH620818, KX924630) causing FV/GVs. |
| *Altenaviridae* | 62 | 6 | 0 | 11 | 0 | ML trees from RdRp alignments; positions with low coverage eliminated. |  |
| *Picornavirales* | 1004 | 118 | 0 | 1004 | 0 | Maximum-parsimony trees from predicted RdRp and helicase domain alignments. |  |
| *Herpesvirales* | 112 | 3 | 0 | 22 | 8 | Neighbour-joining trees from alignments of concatenated predicted AA sequences of six conserved genes. | GVs 3× *Rhadinovirus*, 2× *Percavirus* genomes rearranged within *Gammaherpesvirinae*. GV 3× *Mardivirus* split main cluster, rearranged within *Alphaherpesvirinae*. |
| *Bacilladnaviridae* | 287 | 8 | 0 | 41 | 3 | ML trees (PhyML) constructed from alignments (Mafft) of representative AA sequences; unsupported branches collapsed (TreeGraph2). | GVs *Circoviridae* MH545516, KT73278{5–6} cluster with *Nenyaviridae*. |
| *Alpharhabdovirinae* | 273 | 1 | 0 | 45 | 0 | ML trees from complete L protein sequence alignments. |  |
| *Pestivirus* | 21 | 1 | 0 | 2 | 0 | ML tree from alignment of RdRp sub-domain (conserved NSB5 peptide). |  |
| *Botourmiaviridae* | 154 | 1 | 0 | 12 | 2 | ML trees (IQ-TREE) from multiple RdRp AA alignments (Mafft); low quality regions trimmed (trimAI). | GV *Magoulivirus* MK189195 clusters with *Betabotoulivirus*. *Rhizoulivirus* outgroups from *Betarhizoulivirus*. |
| *Arenaviridae* | 63 | 1 | 0 | 5 | 0 | ML trees from complete L and NP protein AA alignments. |  |
| *Baculoviridae* | 100 | 1 | 0 | 4 | 0 | ML tree (RAxML) from concatenated alignments of 38 core gene AA sequences. |  |
| *Imitervirales* | 22 | 1 | 0 | 15 | 0 | ML tree (IQ-TREE) from concatenated alignment of seven marker genes. | Sections of MT663534, KU877344 non-coding, but no violations. |
| *Parvoviridae* | 172 | 1 | 0 | 29 | 4 | Bayesian inference on three domains of the NS1 protein (BEAST, PhyML). | GVs *Sandeparvovirus* OK236393 and *Muscodensovirus* MT498824 outgroup. GV *Dendroparvovirus* MG74567{0,7} cluster with *Erythroparvovirus*. |
| *Mammonoviridae* | 41 | 7 | 0 | 39 | 0 | ML tree (IQ-TREE) from concatenated AA alignment of seven marker genes (Mafft). |  |
| *Cripavirus* | 17 | 1 | 0 | 3 | 0 | ML trees from nonstructural ORF alignments. |  |
| *Cytorhabdovirus* | 277 | 1 | 0 | 45 | 0 | ML trees from L protein sequence alignments (Muscle) |  |
| *Ephemerovirus* | 269 | 1 | 0 | 45 | 0 | ML tree from alignment of complete L protein sequence alignments. |  |
| *Fraservirus* | 118 | 3 | 0 | 20 | 0 | ML trees (IQ-TREE) from RdRp and bunyaviral-like glycoprotein alignments. |  |
| *Hartmanivirus* | 64 | 1 | 0 | 5 | 0 | Distance matrices for L and S segments. |  |
| *Lispiviridae* | 30 | 1 | 0 | 24 | 0 | ML trees (ModelTest-NG) from RdRp AA alignments (Mafft). |  |
| *Mammarenavirus* | 74 | 1 | 0 | 10 | 0 | Bayesian trees from nucleotide GPC, NP and L gene alignments. |  |
| *Myonnaviridae* | 31 | 1 | 0 | 8 | 0 | ML tree (IQ-TREE) from RdRp AA alignments. |  |
| *Nyamiviridae* | 21 | 3 | 0 | 22 | 0 | ML tree (IQ-TREE) from L protein AA alignments (Mafft). |  |
| *Orthobunyavirus* 017M | 143 | 1 | 0 | 7 | 0 | ML trees from alignments of S, M and L segments. |  |
| *Orthobunyavirus* 018M | 171 | 1 | 0 | 7 | 1 | ML tree from alignments of L segment (ClustalW). | GV *Orthobunyavirus* MK89661{5–7} outgroups, M segment (MK899616) doesn't code for any ORFs. |
| *Phasmaviridae* | 29 | 1 | 0 | 8 | 0 | ML tree from AA alignments of L segment (Mafft). |  |
| *Phenuviridae* | 142 | 1 | 1 | 21 | 6 | ML tree (IQ-TREE) from RdRp AA alignments (Mafft). | FV MW74189{4–6} all segments partial CDS with ~ 30% gaps. GV 6× double-segmented *Uukuvirus* cluster away from triple-segmented *Uukuvirus*. |
| *Varicosavirus* | 276 | 1 | 0 | 6 | 0 | ML tree from L protein AA sequences (Mafft). |  |
| *Vesiculovirus* | 269 | 1 | 0 | 31 | 0 | ML tree from L protein AA sequences (Muscle). |  |

**Fig. 5.**
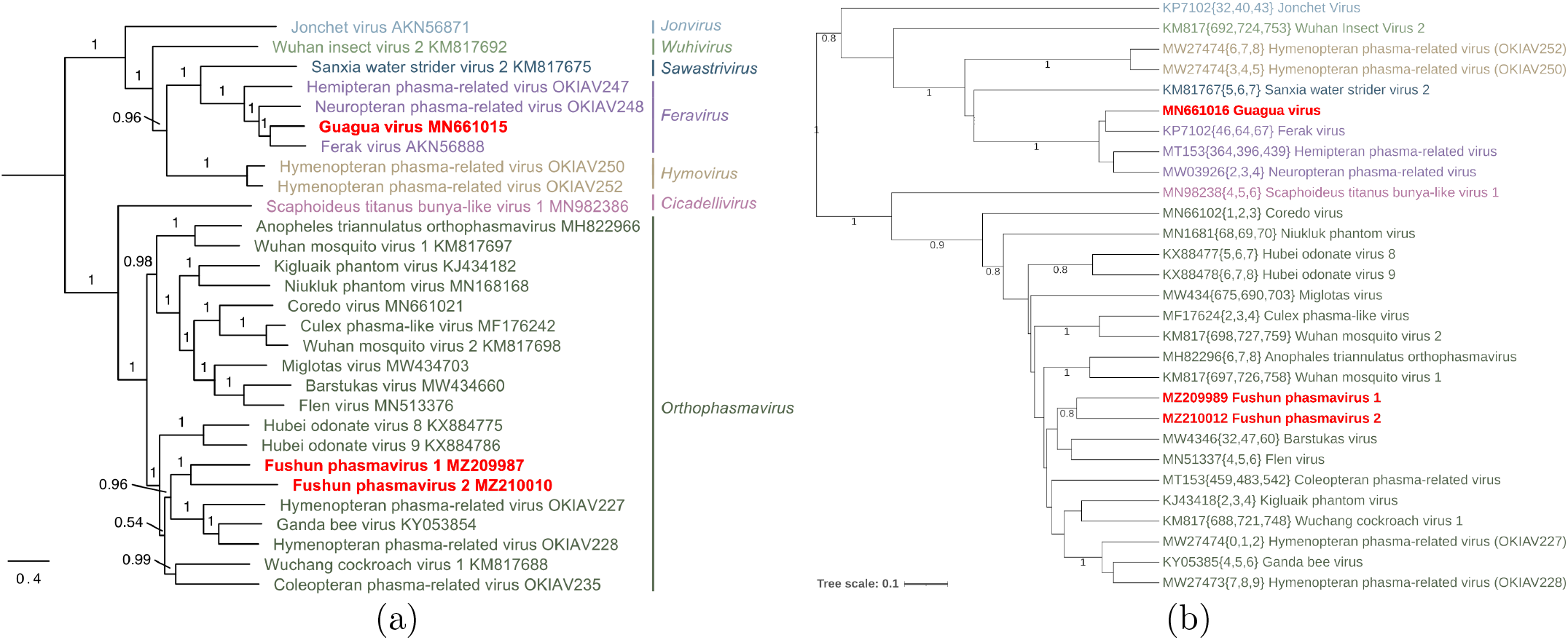
*Phasmaviridae* trees. (a) TP ML tree produced by alignment of L segment AA sequences [19]. (b) GRAViTy-V2 tree, with colours adjusted to match TP. (Red text: sequences proposed as new taxa in TP; bootstrap values *<* 0.7 hidden).

Of the comparisons yielding different virus relationships between methods, two were at family-level, in the orders *Phenuiviridae* and *Jingchuvirales* (Table 2). In the former, the outlier was a tripartite genome with three coding incomplete GenBank sequences that precluded a valid comparison with their current taxonomy, as GRAViTy-V2 requires coding complete genomes for estimation of sequence relationships. In the *Jingchuvirales* set, five of seven family-level (and one genus) violations were caused by the original inadvertent inclusion of a large number of incomplete genomes by the Study Group responsible for the original classification, which was based on an RdRp alignment (Fig. 6). Incomplete sequences were identifiable in both shared normalised PPHMM ratio and barcode heatmaps, as long bands of continuous colour and absent profiles, respectively (Fig. 4). Violations either directly involved the incomplete genome sequences themselves (five), or indirectly where complete genomes were pulled out of their correct family assignment through their similarity to the incomplete sequences. In consequent experiments, the *Jingchuvirales* dataset was improved by locating missing components of four incomplete sequences (BK061669, KX924630, MW645032, MH620818) and removing those for which no replacements could be found (BK0616*{*70,72*}*), which reduced the number of family violations to three zero and the number of genus violations to one (SI Document 1, S2).

**Fig. 6.**
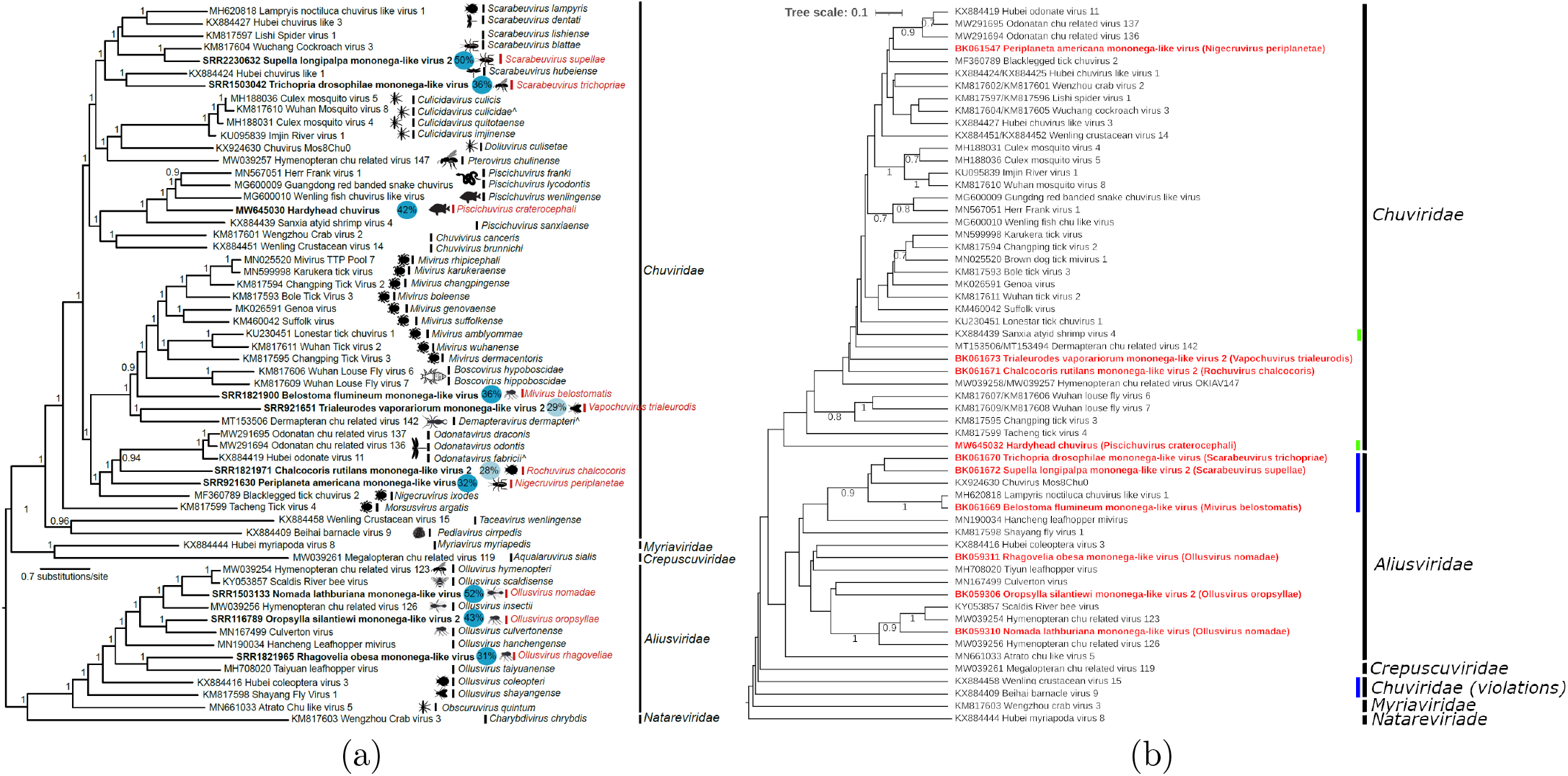
*Jingchuvirales* TP. (a) TP ML tree, from RdRp alignment [20]. (b) GRAViTy-V2 tree (red: sequences proposed as new taxa in T; blue: family violations; green: genus violations; bootstrap values *<* 0.7 hidden).

Datasets from seven taxonomy proposals resulted in genus-level violations (Table 2) and the majority of these exhibited sub-genus variation in trees provided in TPs based on phylogenetic analysis of aligned conserved genome regions. The rearrangements observed in the GRAViTy-V2 output did not however violate their original taxonomic classifications. In investigating the causes of these discrepancies, two genus violations were the result of the original inclusion of non-coding complete sequences for which replacements could not be found, leaving five datasets where were genuine discrepancies. These tended to occur in instances where clades contained genomes with either a variable quantity of segments (e.g. *Bacilladnaviridae*, SI Document 2, Figs. 9-10); a single, very large ORF (e.g. *Mymonaviridae*, SI Document 2, Figs. 41-42); or very remote homologies at the limit of what PPHMM profile comparisons can detect, as indicated by barcode graphs (*Herpesvirales*, SI Document 2, Fig. 7-8).

Several TPs proposed the creation of new taxa and GRAViTy-V2 results were concordant with these, notwithstanding the violations listed (e.g. *Alternaviridae*, SI Document 2, Fig. 3). Where this is not always obvious in visual inspection of GRAViTy-V2 heatmaps, especially at lower ranks, the software additionally estimates virus groupings including new taxon assignments, results for which are automatically reported in tabular output. This feature is calculated using hierarchical clustering using Thiels U Statistic derived from GRAViTy-V2’s scoring system and represents a key use case for GRAViTy-V2 over alignment-based methods, as its identification of new taxa functions independently of the inclusion of reference sequences with definite homologies.

## Discussion

### GRAViTy-V2 performance and proper use

GRAViTy-V2 has more than thirty optional parameters, the majority of which control the command line tools the framework uses (Mash, BLASTp, Mafft, HMMER, HHsuite). While every attempt has been made to simplify the user experience, there is no single set of default parameters that will suit the requirement of every user. However, only a few parameters significantly influence classifications, specifically the Mash similarity threshold, minimum protein length, use of BLASTp or Mash and HMMER bitscore threshold. Hence, these parameters should be selected carefully and with reference to knowledge of the genomes being analysed. Furthermore we have created several easy-to-use premade pipelines with parameter sets optimised for specific scenarios (comparison of similar, divergent or extremely long viral genomes) for which users only need specify their input VMR, input sequences (or location to automatically download their sequences to from GenBank) and output folder.

Experiment duration was proportional to the length and number of genomes analysed, and the optional user-selected parameters. It is advised to not classify datasets of *>* 1000 genomes with GRAViTy-V2 in a single pass and instead use a multiple pass methodology should users wish to keep run times within the range of minutes, rather than hours. Some parameters may significantly increase run time and but are not likely to improve classifications in the majority of use cases (e.g. alignment merging, PPHMM sorting). Users should use default parameters in the first instance and refine them iteratively as required. A user guide for optimising GRAViTy-V2 parameters is included in the GitHub repository.

### Mismatches between GRAViTy-V2 results and ICTV ratified taxonomy proposals

Input sequence quality was found to be the cause of all cases of family violations and approximately one fifth of genus violations. Inclusion of non-coding complete sequences, such as in the *Jingchuvirales* dataset (Fig. 6), or segmented genomes assembled inconsistently (SI Document 1, S3), were found to dramatically impact classifications. In initial experiments, an additional four experiments exhibited genus-level violations as result of including incorrect (mislabelled accession IDs or incomplete genome) sequences, all of which were corrected manually and re-run. Problematic sequences were easy to identify in GRAViTy-V2 graphs (Fig. 6; SI Document 1, S4), whereas this is traditionally a challenging manual task when working with large quantities of novel genomes, especially when they are metagenomically-derived. These observations highlight how taxonomists should be mindful of sequence quality in public databases.

There were several scenarios in which GRAViTy-V2 did not perform optimally, as measured by genus-level violations. Foremost were instances where viruses had single, extremely long ORFs, such as in the *Mymonaviridae*. The minimal genomic unit by which features are compared in GRAViTy-V2 is the ORF and the initial phase of analysis involves computing a metric of pairwise sequence relatedness (Mash distance or BLASTp bitscore), after which ORFs are clustered and aligned. In these instances, long ORFs may have limited similarity between even closely-related genomes and furthermore, the effectiveness of the GRAViTy-V2 GOM is reduced when a single genomic component is present. The issue was however mitigated in this instance through changing run parameters, specifically switching Mash to BLASTp to reduce the sensitivity of initial PPHMM clustering, reducing BLAST coverage thresholds and changing similarity scoring scheme to one that didn’t use the GOM (R-scheme). For the same reason, GRAViTy-V2 is unlikely to classify two genomes within the same taxon if they contain a different number of segments, which was the cause of violations in the *Phenuiviridae* dataset.

### Use cases for GRAViTy-V2 in comparison with other tools

When choosing viral taxonomy software, users should be aware that there is currently no package that will suit the needs of every user. Certain tools rely on building pairwise distances between genomic components, including PASC [11] and newer derivatives such as DEmARC [21]. These algorithms are effective for differentiating between similar genomes at species and offten genus level, but not in defining threshold similarity thresholds for family or higher rank assignments. VISTA [22] is an increasingly-popular tool for lower rank assignments that greatly builds on the pairwise similarity paradigm and is highly scalable, but is, however, reliant on a high reference data input requirement and prone to bias should any taxa be over-represented.

A range of newer approaches based on intergenomic distance metrics, which utilise variations on percent identity as base metrics but make additional novel use of network and clustering algorithms, have emerged in recent years, especially for classification of viruses infecting bacteria and archaea (e.g. vConTACT [23], VIRIDIC [24]). These approaches are comparatively rapid and effective for screening larger viruses and may be used in concert to support findings, but are not complete taxonomic tools and are usually not phylogeny-aware. Virus relationships as determined by GRAViTy-V2, conversely, are based on holistic comparisons of all coding regions within virus genomes. Its output is not based on rules for taxonomic assignments of specific virus groups, such as the reliance on specific marker or hallmark genes such as the RdRP of RNA viruses, nor on assignments based or the adoption of a variety of threshold similarity values to define genera and species. Such methodologies, based as they are on the properties of individual virus families, genera or species definitions, remain in the province of expert curation and cannot be reproduced by a general metric of genome relatedness as calculated by GRAViTy-V2, although our software’s ability to propose new taxa based on analysis of multiple extracted genomic features would be an informative adjunct to this process.

We therefore envisage the following uses in the analysis of virus metagenomic sequence data:

a. Running GRAViTy-V2 requires only the test sequence(s) or its accession number. Comparisons are made with the dataset of classified viruses accessed from ICTV databases (i.e. VMR) by the program itself. It therefore provides a useful screening tool for initial evaluation of virus sequence data and provisional assignments to orders, families or lower taxonomic ranks. Its output is far more informative than methods such as Kraken for the analysis of metagenomic sequence data that simply report profile or *k*-mer matches.
b. Unlike methods based on alignment construction, an important attribute of GRAViTy-V2-based analysis is that it can also identify and report sequences with no identified homology to existing classified viruses, and which therefore represent entirely new virus groups and component taxa.
c. GRAViTy-V2 provides a general tool for re-examining existing classifications, and identification of incomplete or incorrectly assembled sequences, as exemplified in many of the reported analyses in the study.
d. The reporting of positions of homology through PPHMM matching (Fig. 4b) allows the identification of conserved motifs in stretches of contiguous sequence. These can be selected, exported and used for alignment-based approaches of homologous gene regions. Particularly for very distantly related viruses, the cumbersome and often manual identification, extraction and alignment of conserved genome regions can be avoided through using GRAViTy-V2 output.

## Conclusions

Evaluations of official taxonomy proposals were completed within the scale of minutes to hours and were found to generate classifications similar to those created by expert groups. GRAViTy-V2 aided identification of several human errors, accidental inclusion of non-coding complete sequences and remote homologies (or lack thereof) between genomic components not identified in conventional, conserved region alignments. We propose that GRAViTy-V2’s approximate classifications are suitable for use in support of viral taxonomy workflows including, but not limited to, classification of newly-described viruses.

## Data availability

Data supporting this article (TPs and VMR) were accessed from the ICTV website https://ictv.global/. GRAViTy-V2 software is available via GitHub (https://github.com/Mayne941/gravity2) with a GPL 3.0 license.

## Supplementary data

Supplementary data are available attached.

