## Supplementary material for "GRAViTy-V2: a grounded viral taxonomy application": SI Document 2

Mayne, R., Aiwsakun, P., Turner, D., et al. (2024)

Supplementary information, Document 2

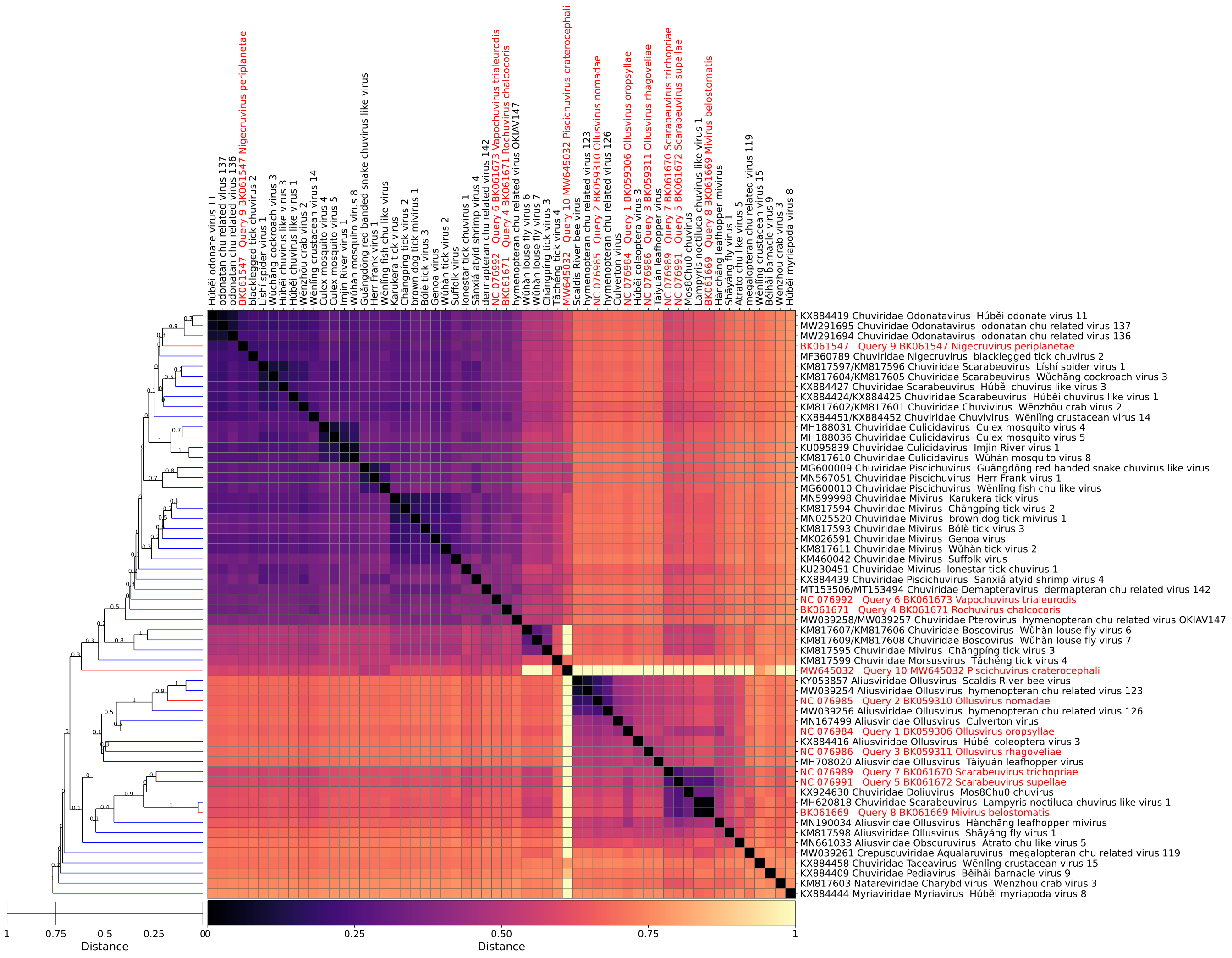


Figure 1: *Jingchuvirales*, GRAViTy-V2 heatmap

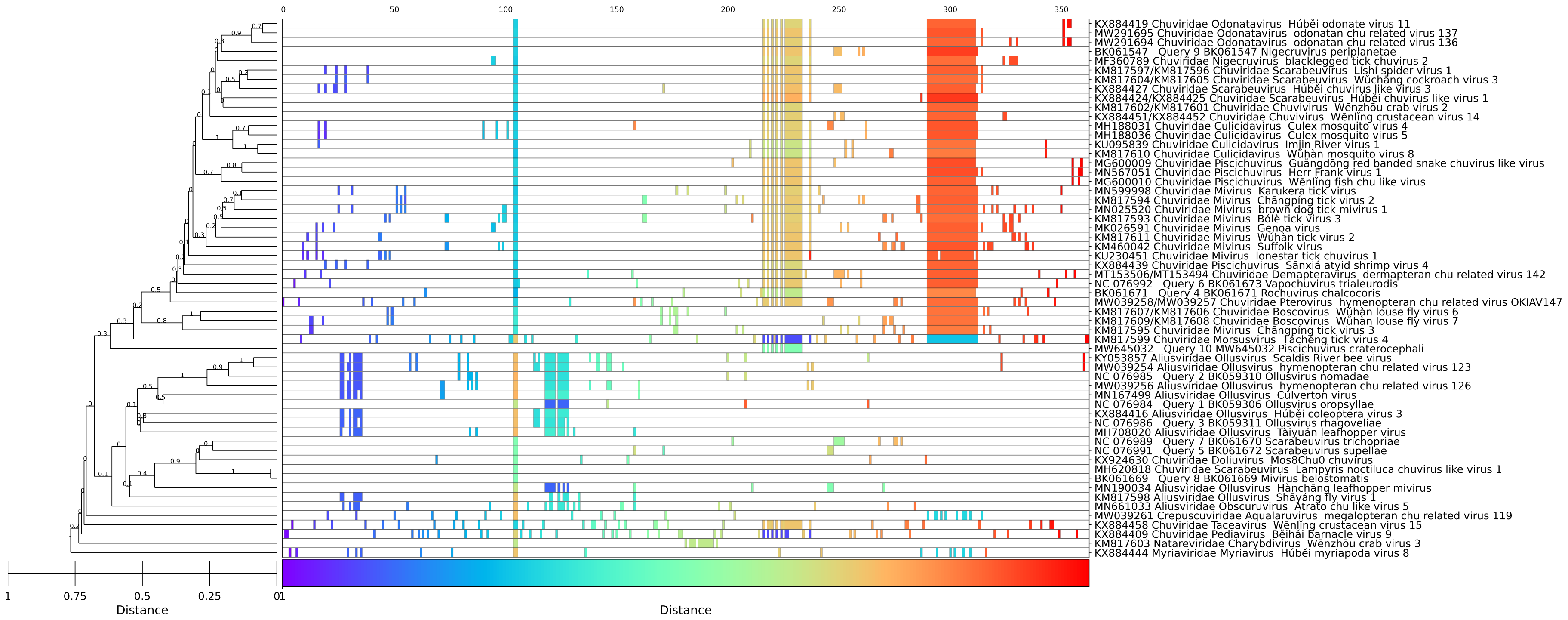


Figure 2: *Jingchuvirales*, GRAViTy-V2 barcode

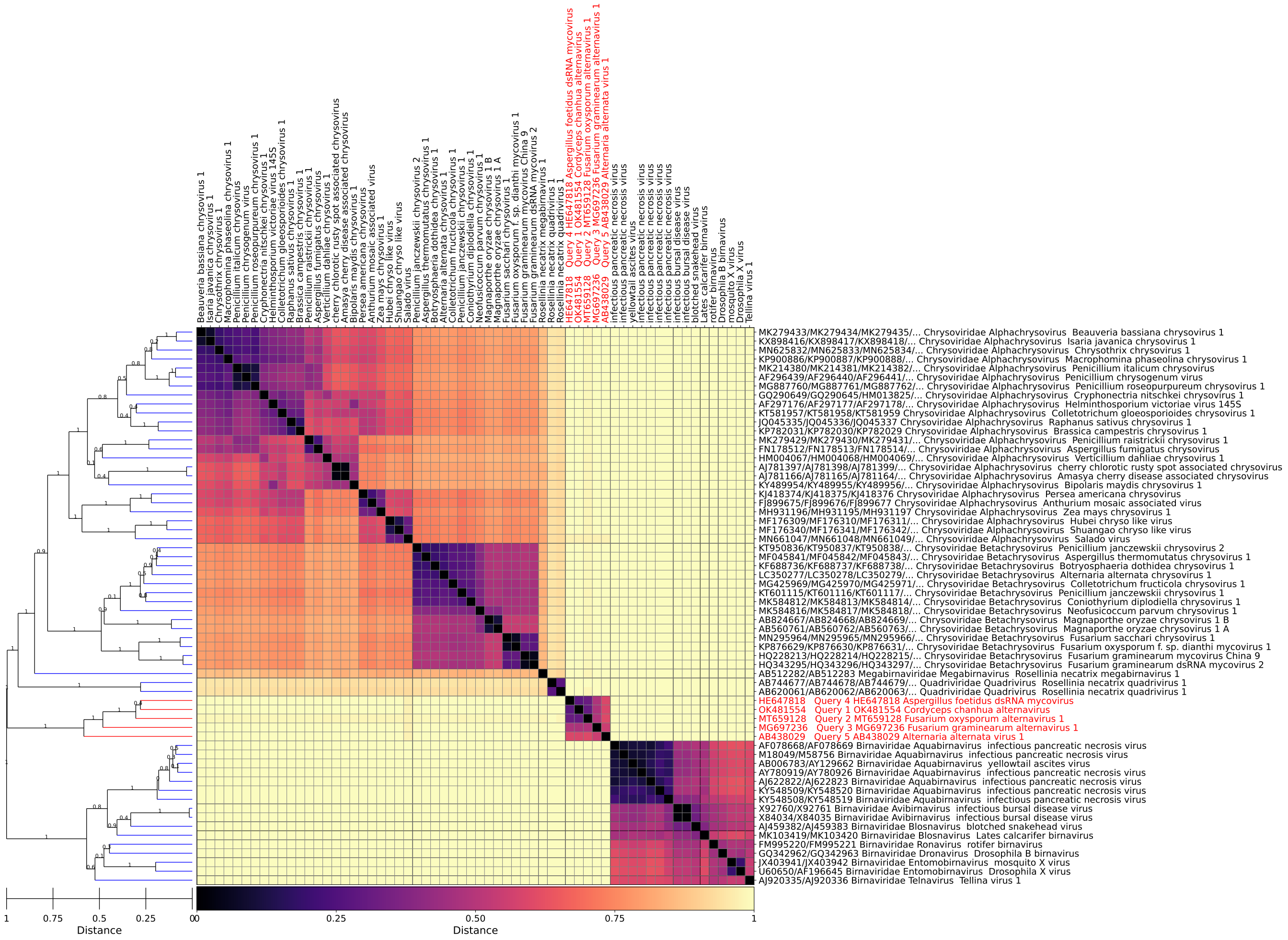

Figure 3: *Alternaviridae*, GRAViTy-V2 heatmap

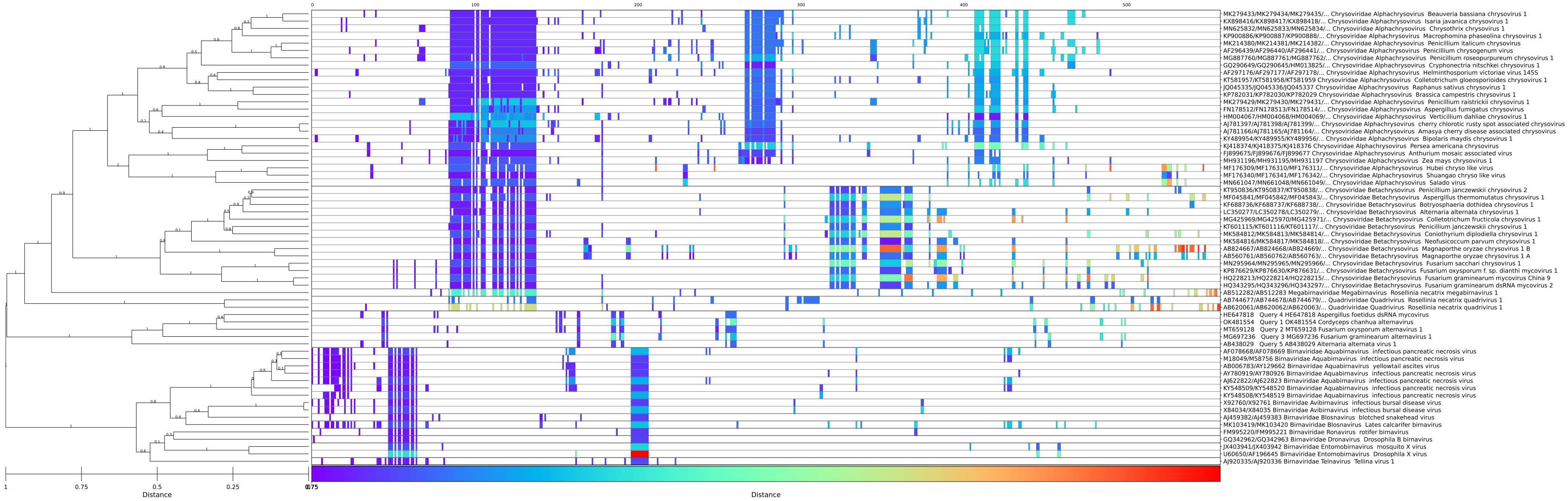


Figure 4: *Alternaviridae*, GRAViTy-V2 barcode

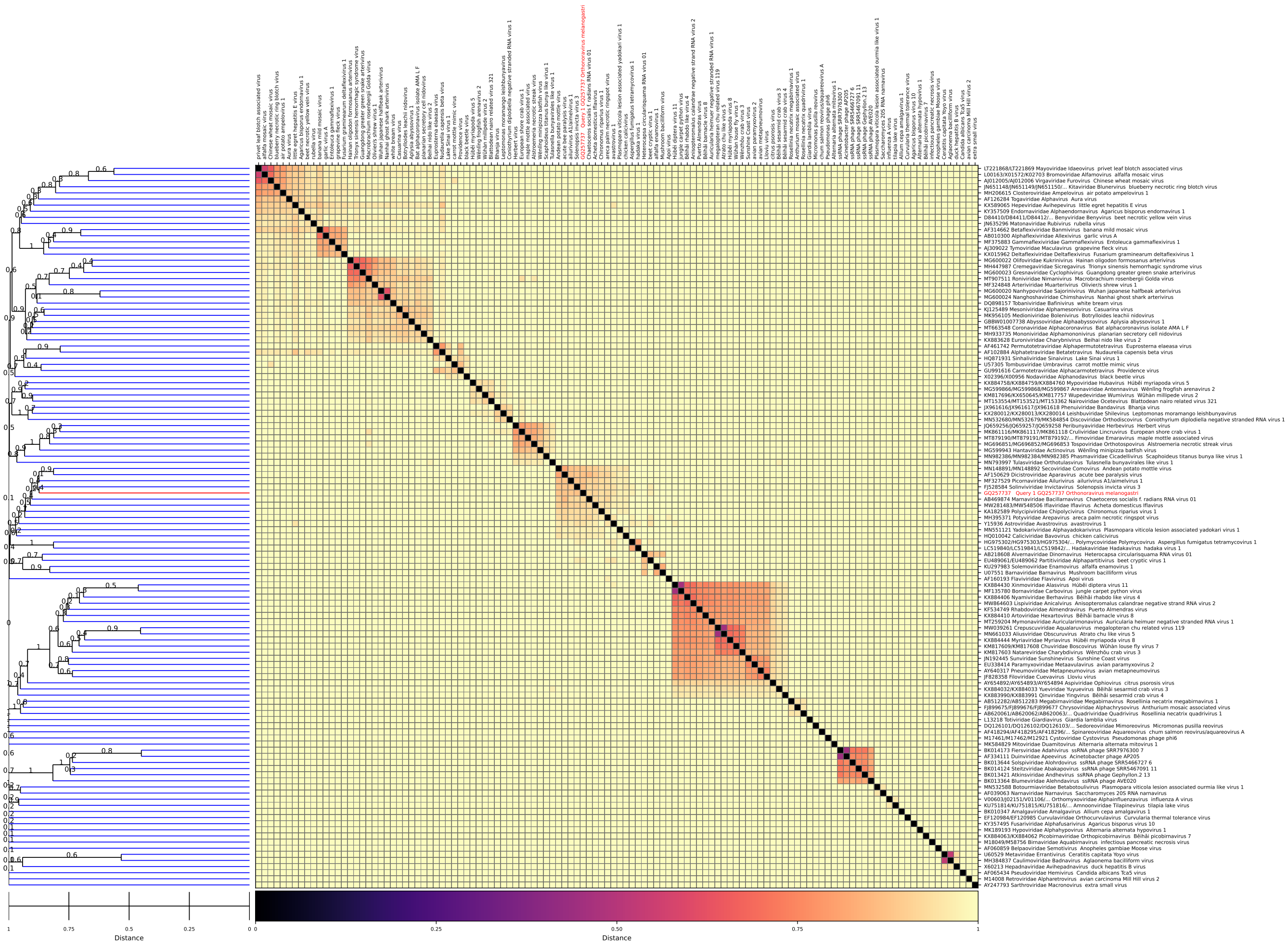

Figure 5: *Picornavirales*, GRAViTy-V2 heatmap

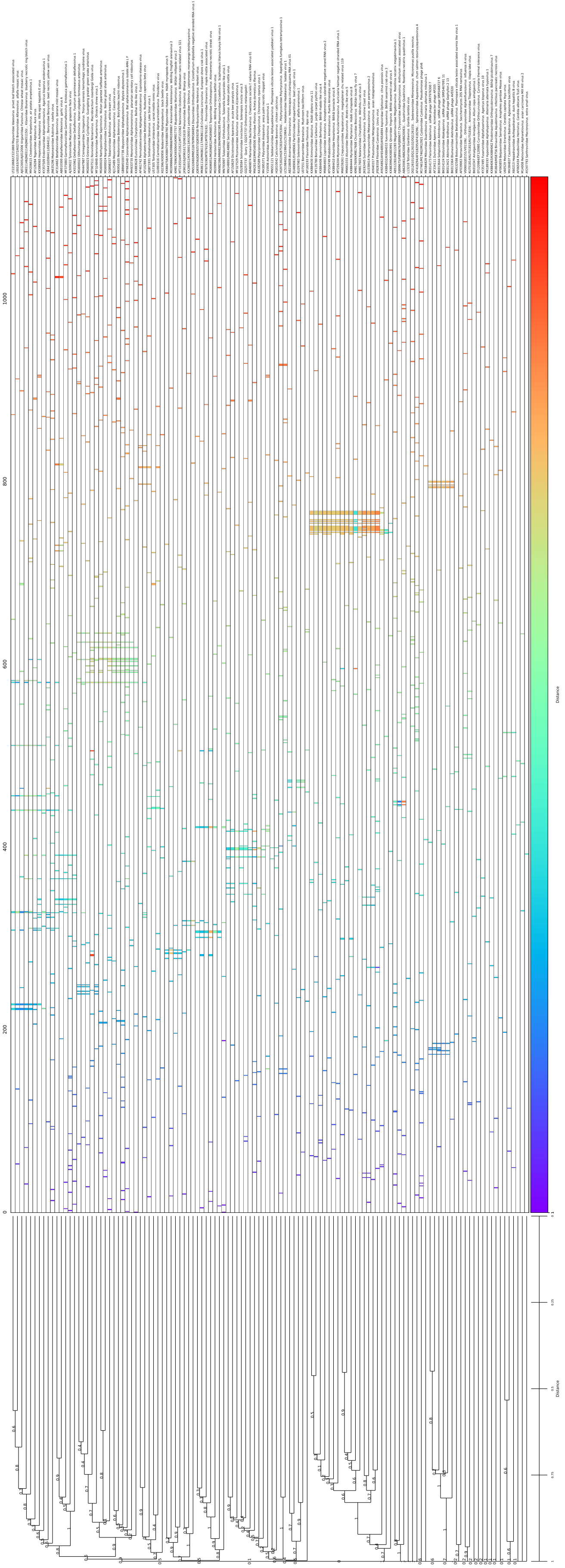

Figure 6: *Picornavirales*, GRAViTy-V2 barcode

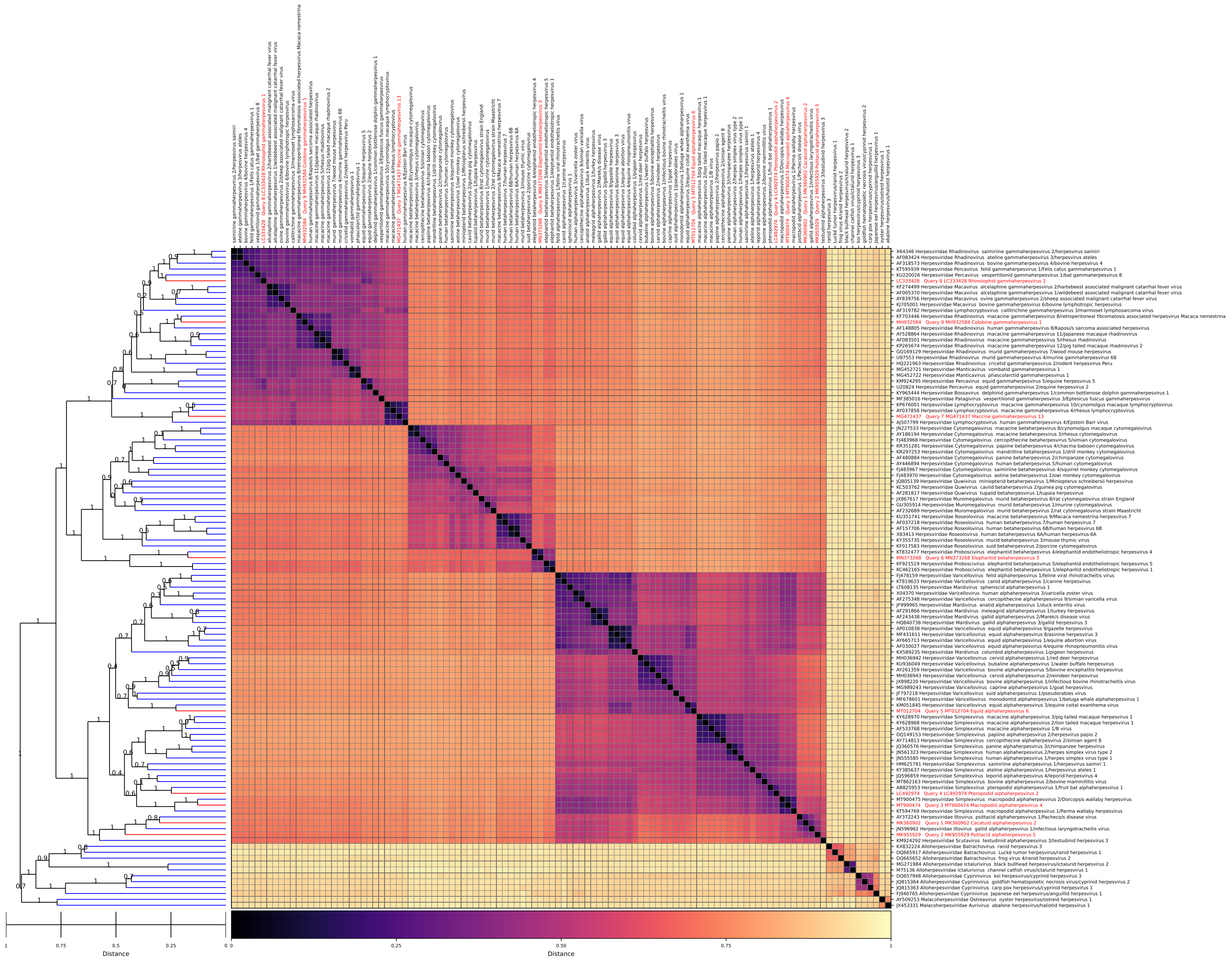

Figure 7: *Herpesvirales*, GRAViTy-V2 heatmap

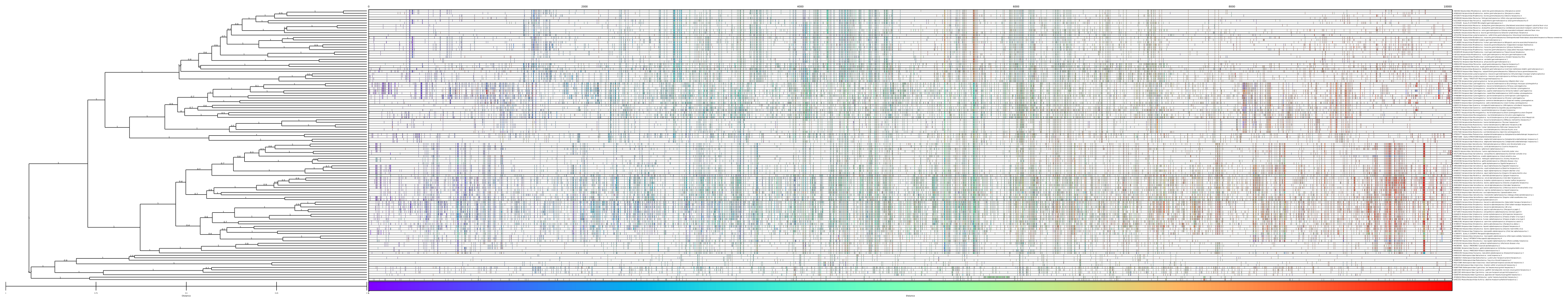

Figure 8: *Herpesvirales*, GRAVITy-V2 barcode

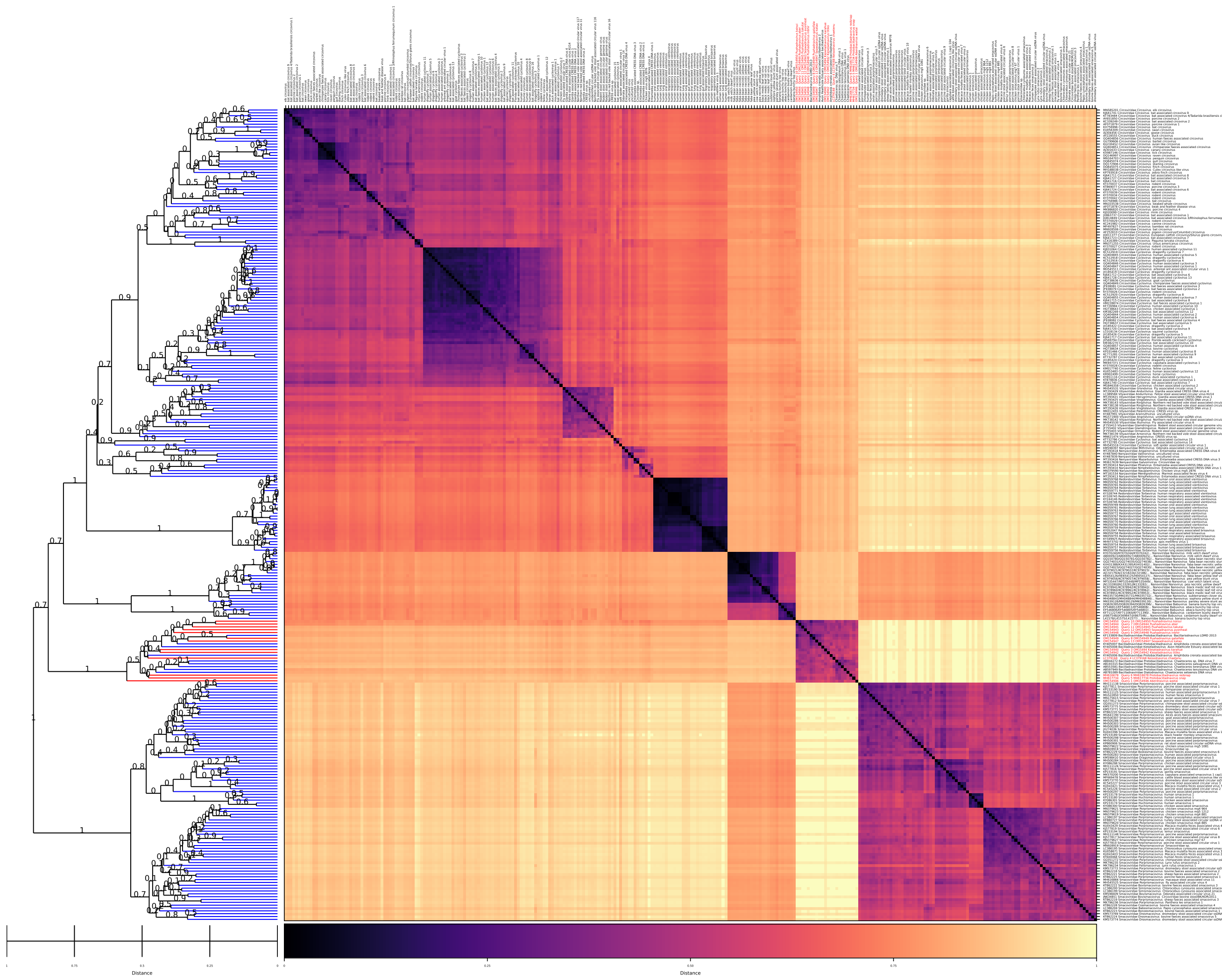


Figure 9: *Bacilladnaviridae*, GRAViTy-V2 heatmap

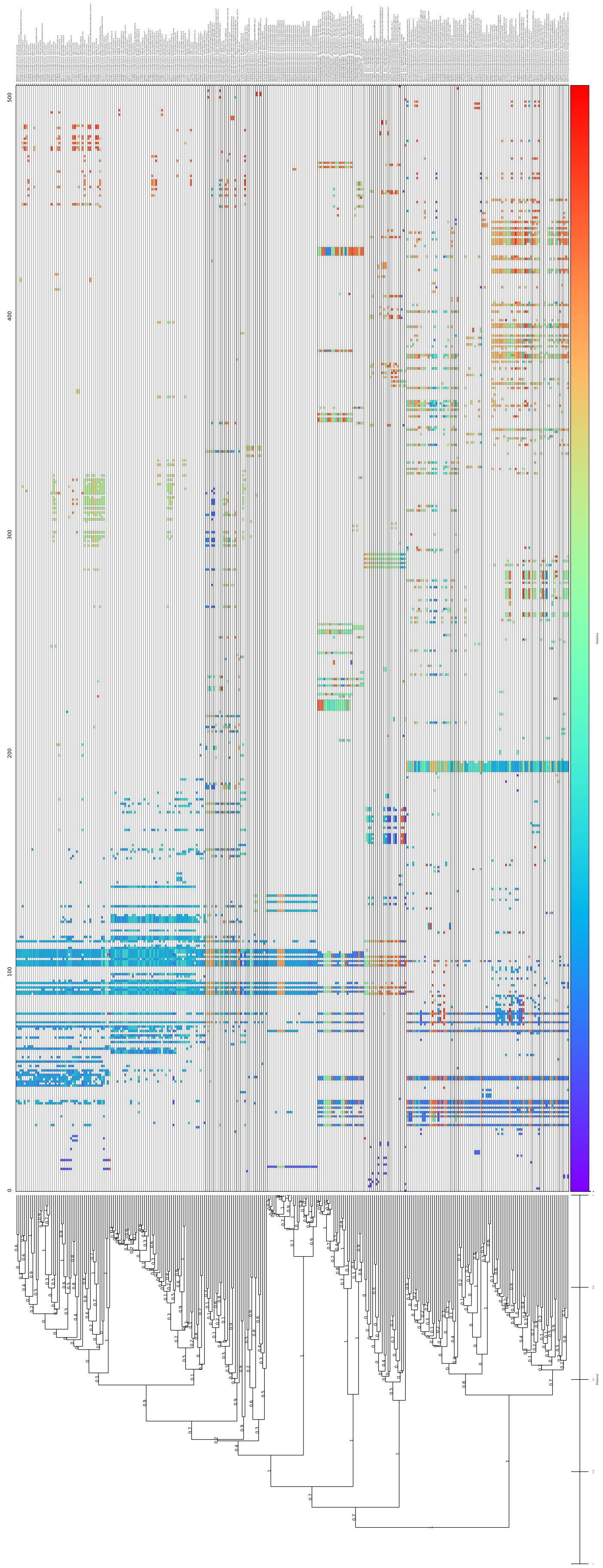

Figure 10: *Bacilladnaviridae*, GRAViTy-V2 barcode

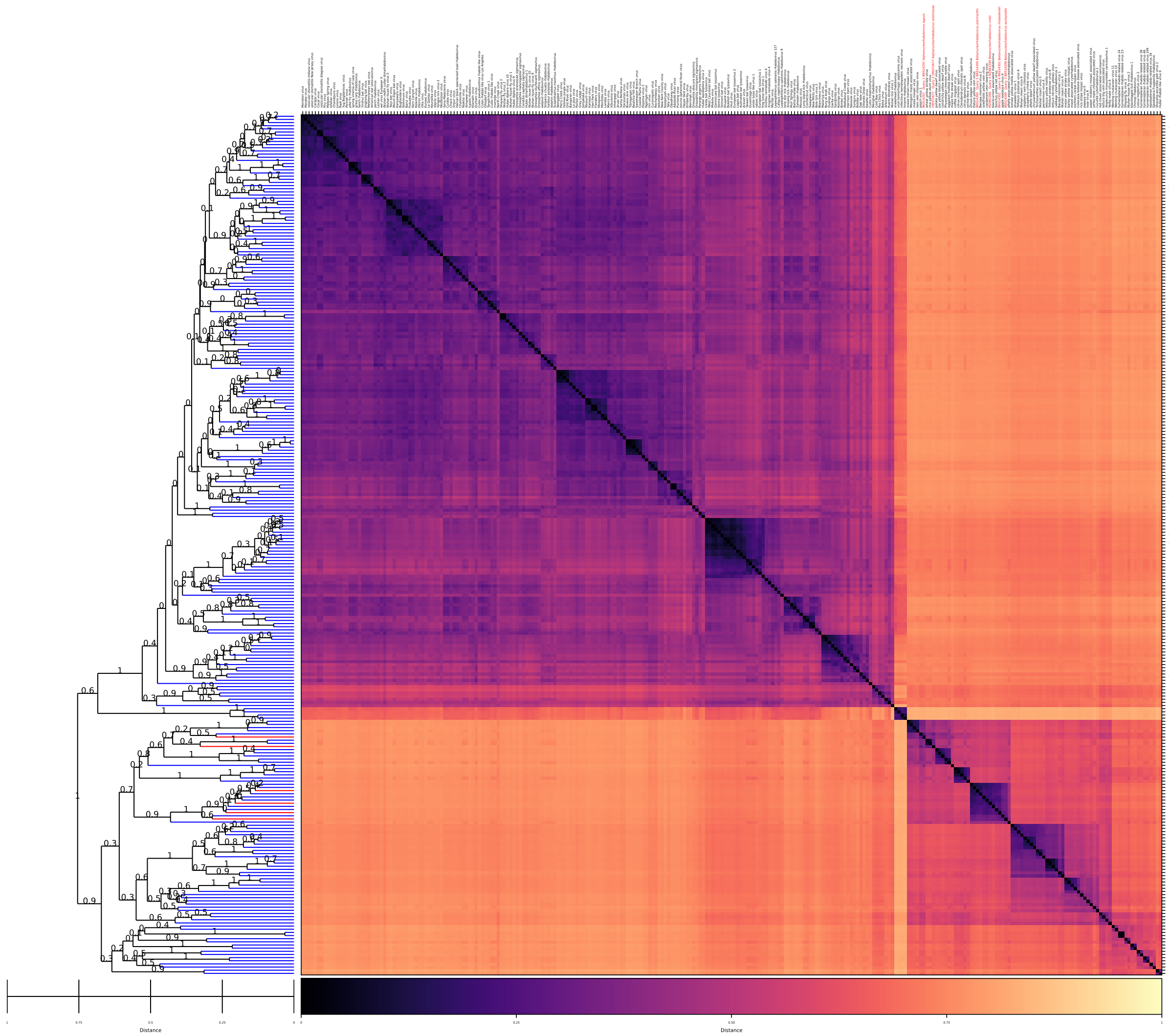

Figure 11: *Alpharhabdovirinae*, GRAViTy-V2 heatmap

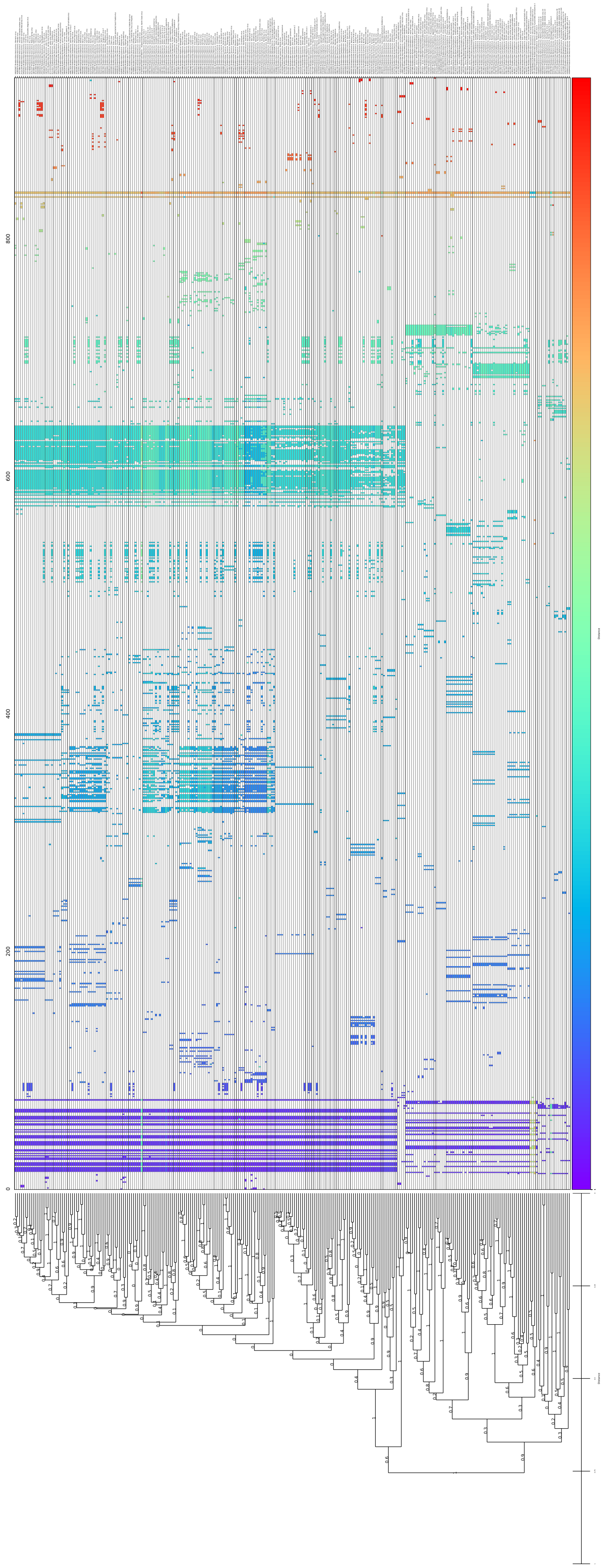

Figure 12: *Alpharhabdovirinae*, GRAViTy-V2 barcode

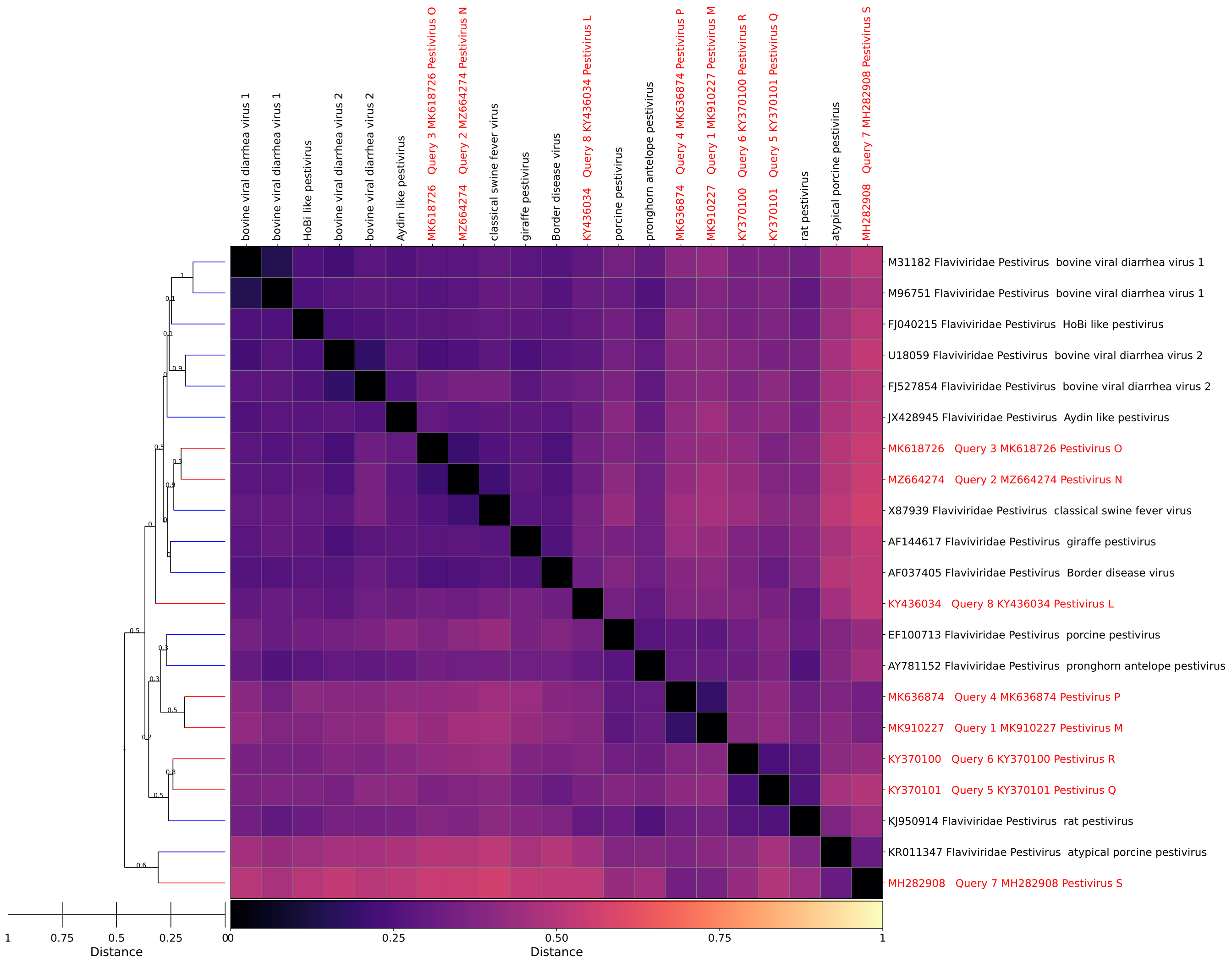


Figure 13: *Pestivirus*, GRAViTy-V2 heatmap

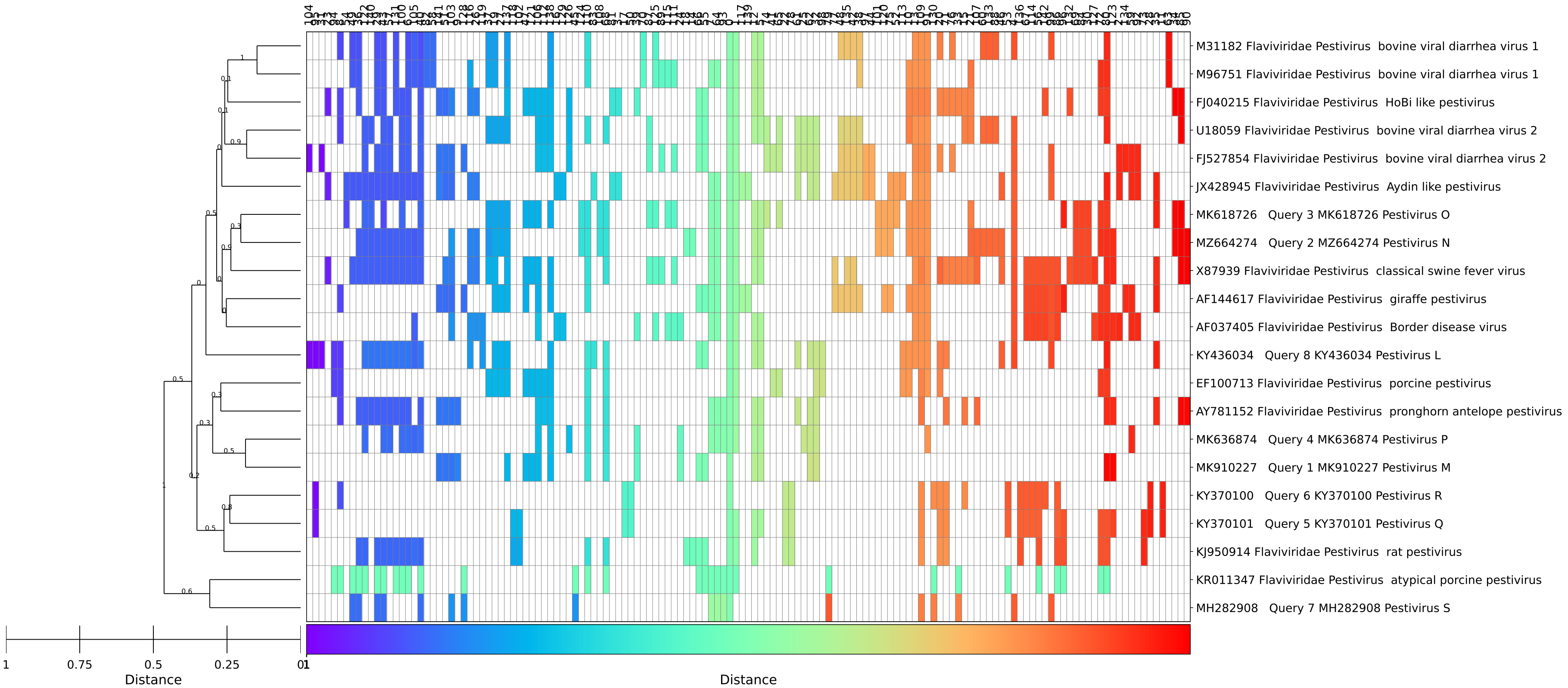


Figure 14: *Pestivirus*, GRAViTy-V2 barcode

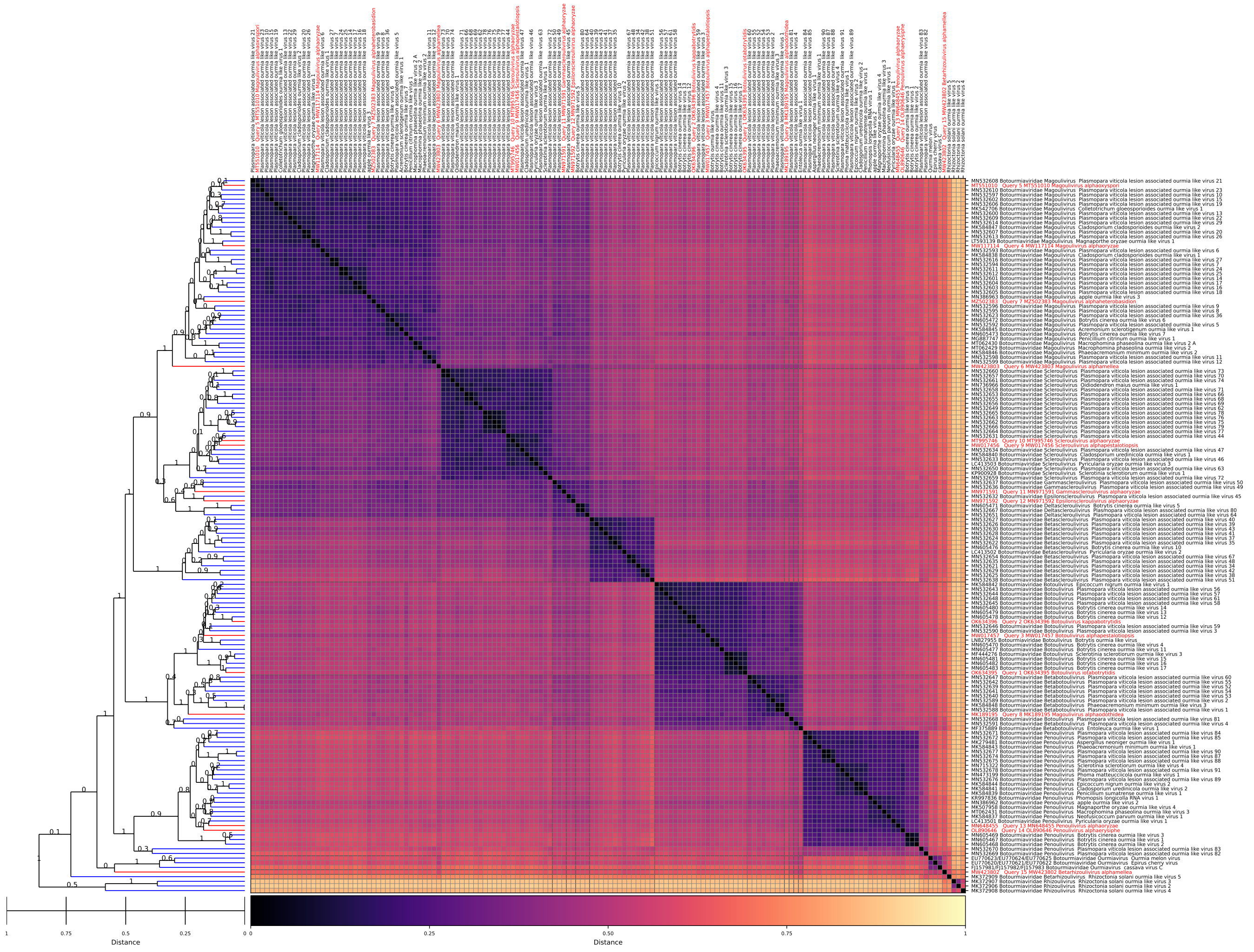


Figure 15: *Botourmiaviridae*, GRAViTy-V2 heatmap

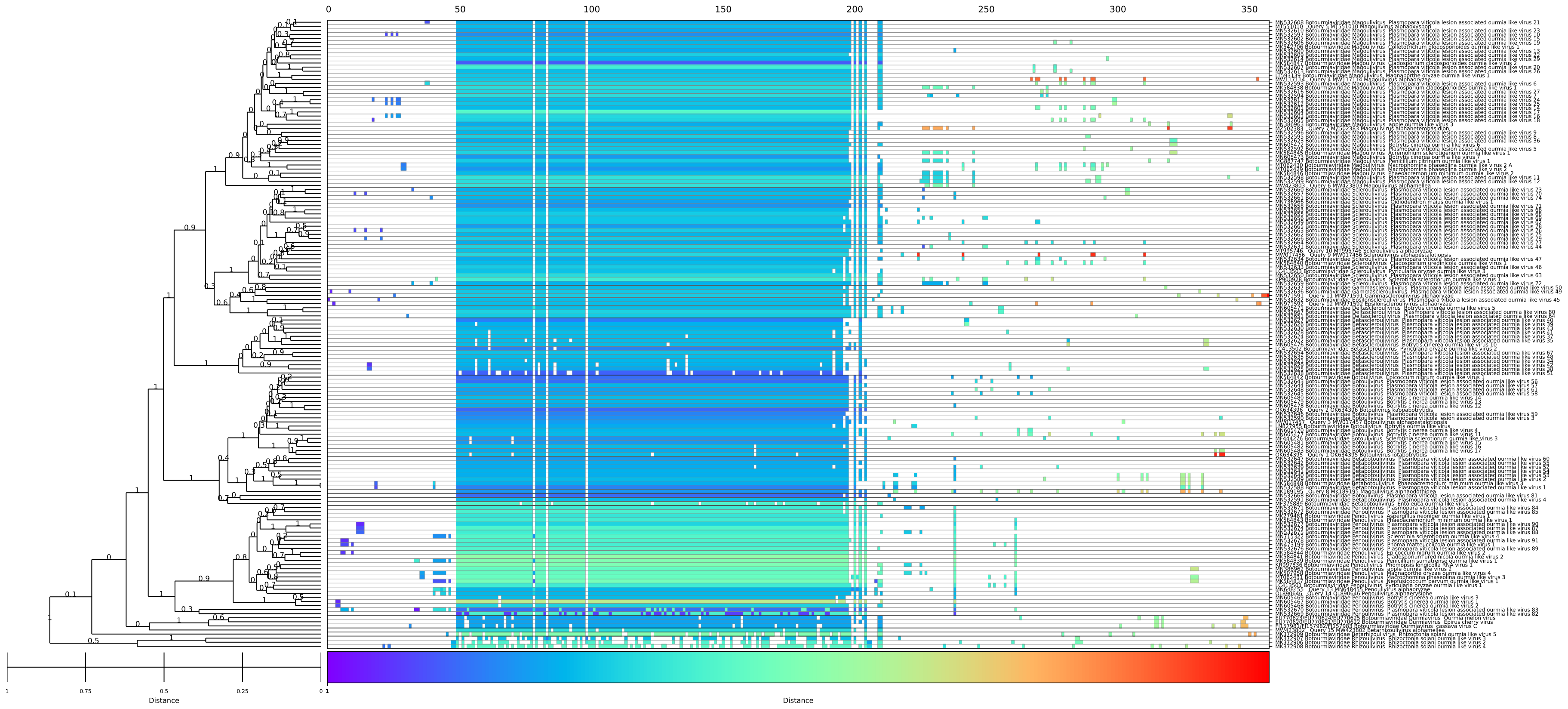

Figure 16: *Botourmiaviridae*, GRAViTy-V2 barcode

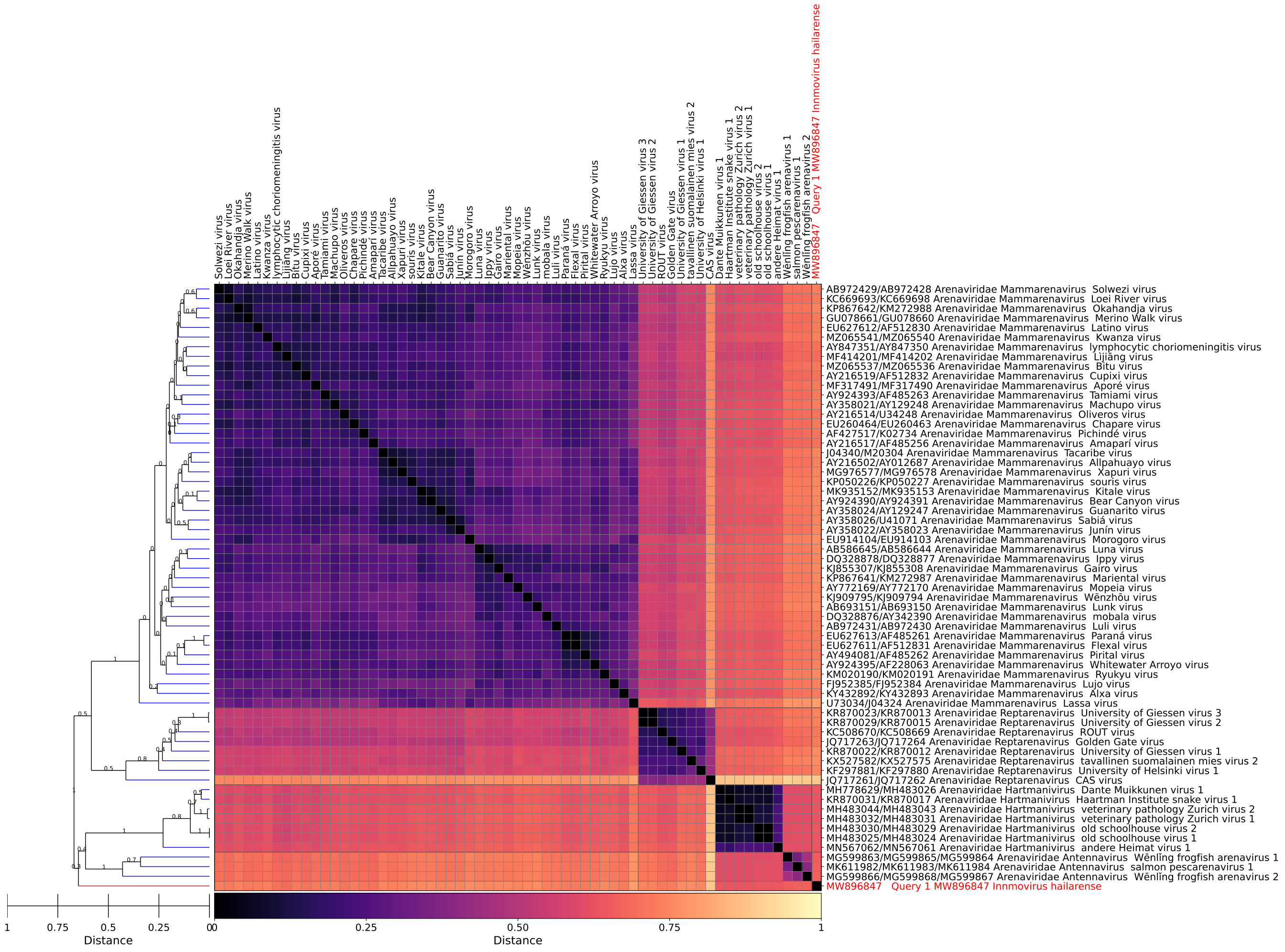

Figure 17: *Arenaviridae*, GRAViTy-V2 heatmap

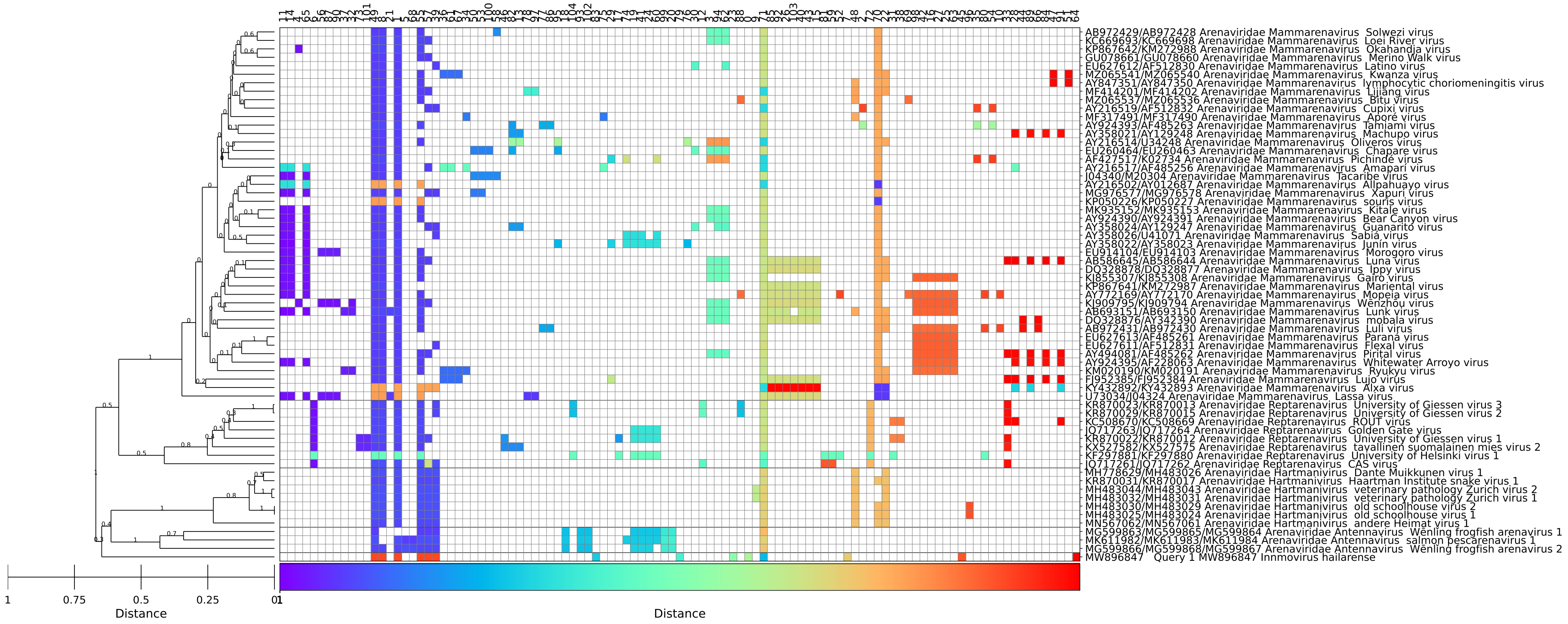


Figure 18: *Arenaviridae*, GRAVITy-V2 barcode

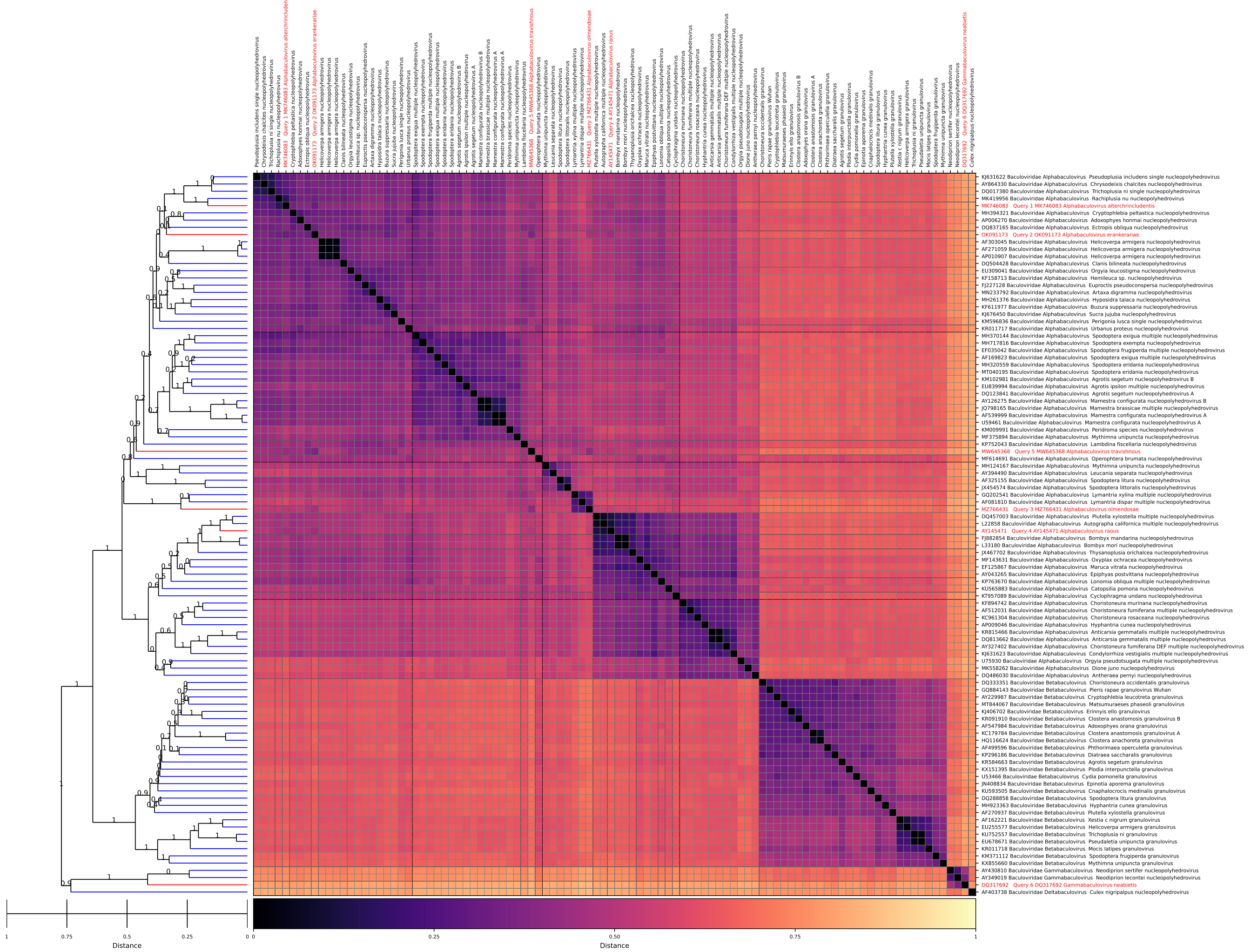


Figure 19: *Baculoviridae*, GRAViTy-V2 heatmap

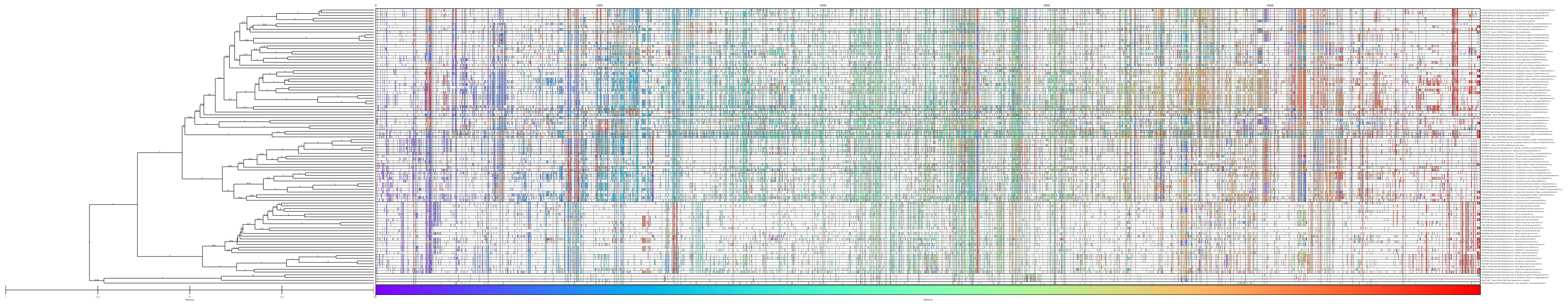

Figure 20: *Baculoviridae*, GRAViTy-V2 barcode

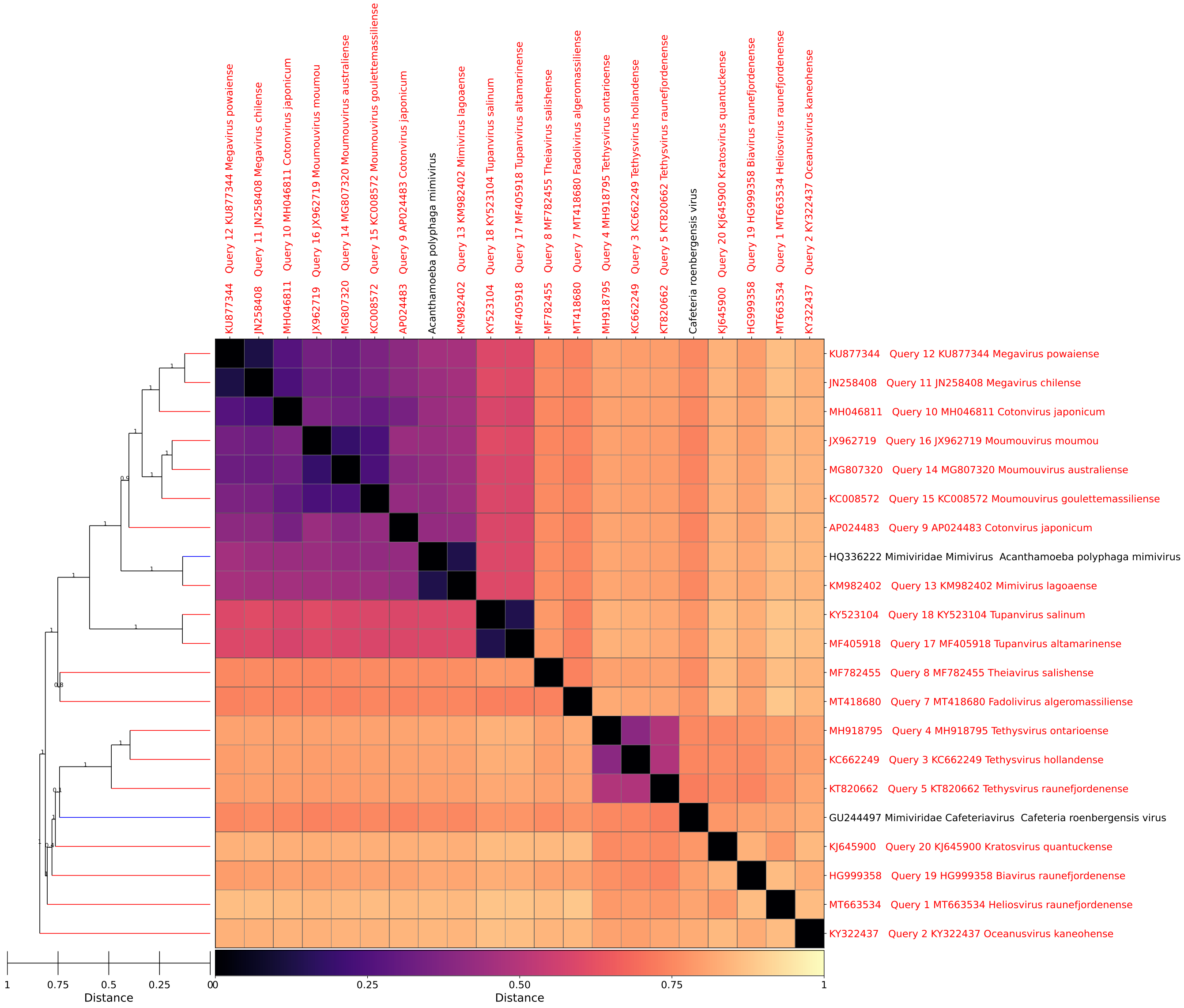


Figure 21: *Imitervirales*, GRAViTy-V2 heatmap

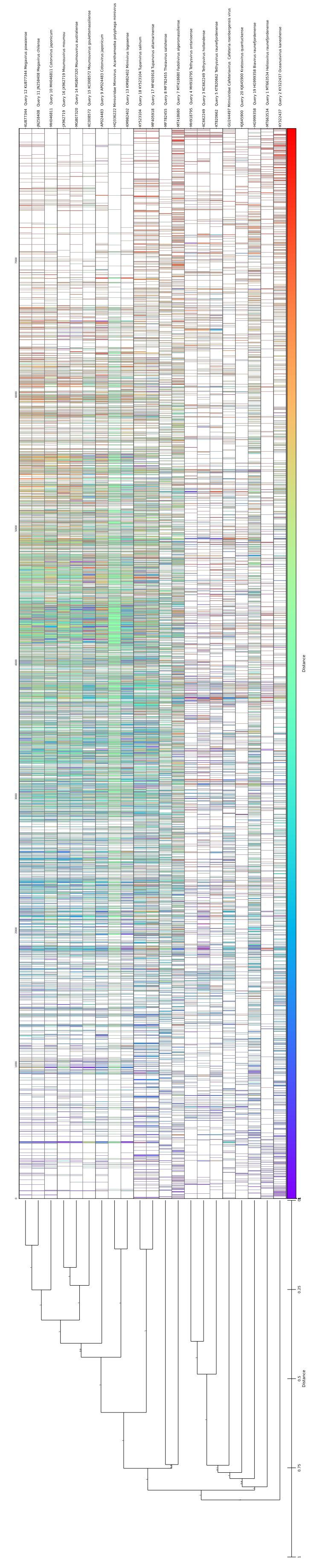

Figure 22: *Imitervirales*, GRAVITy-V2 barcode

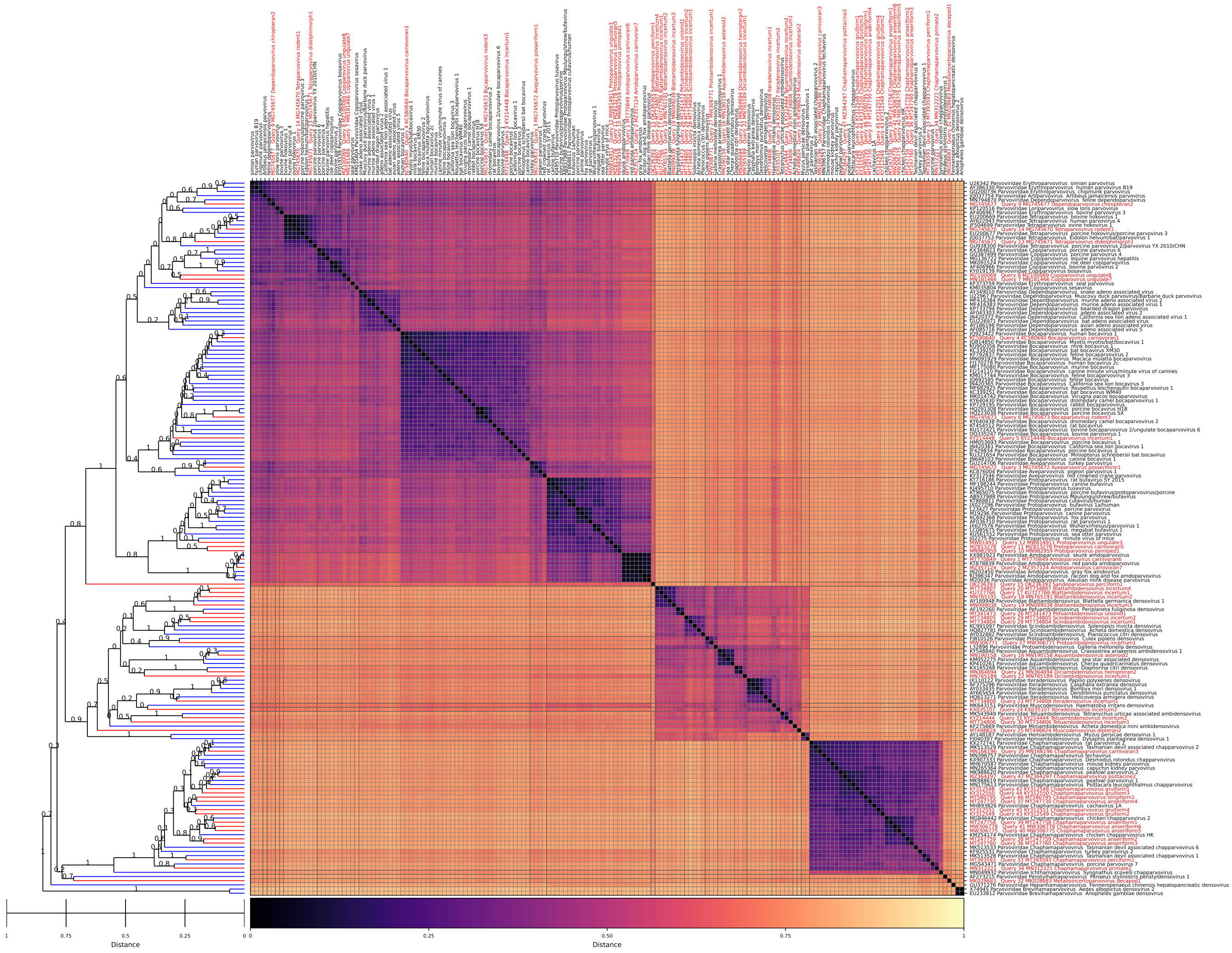


Figure 23: *Parvoviridae*, GRAViTy-V2 heatmap

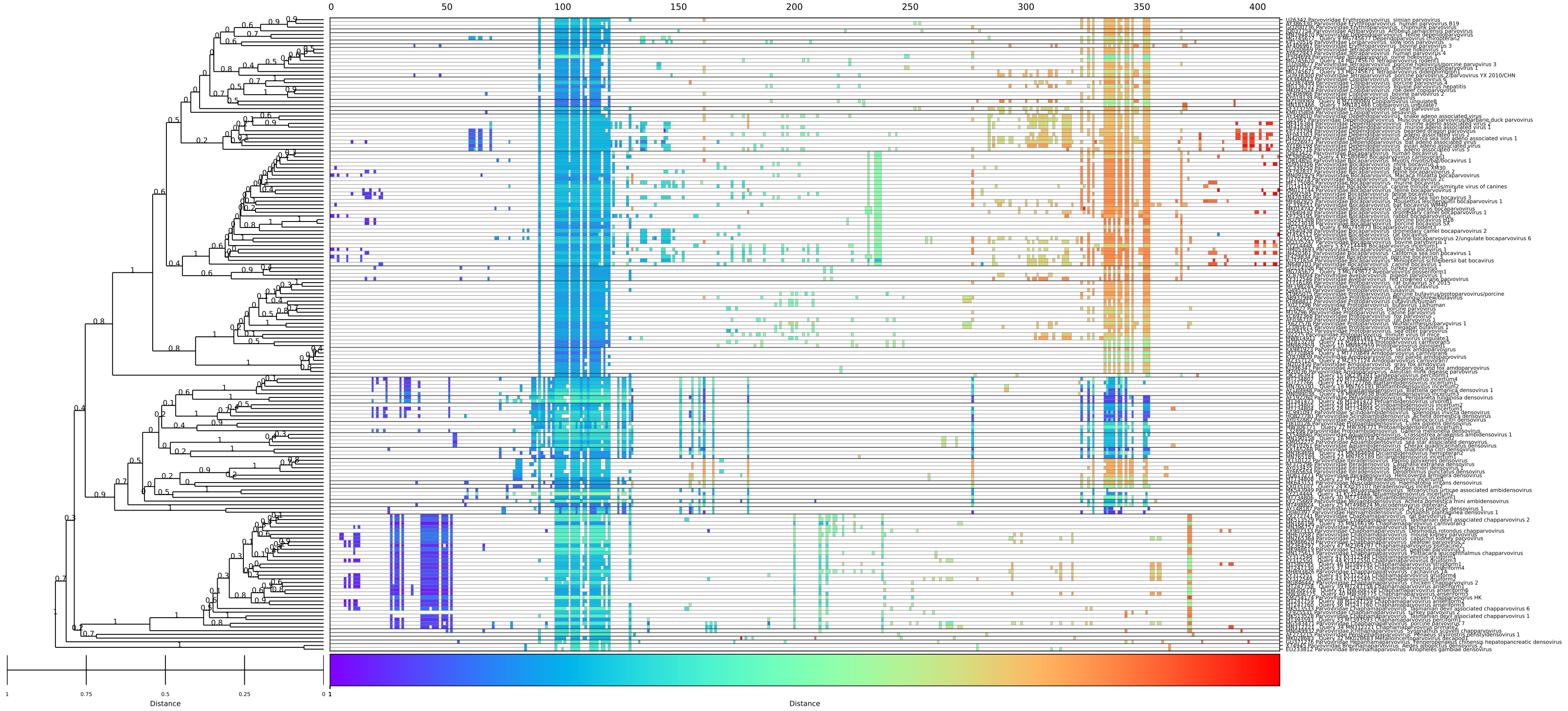


Figure 24: *Parvoviridae*, GRAViTy-V2 barcode

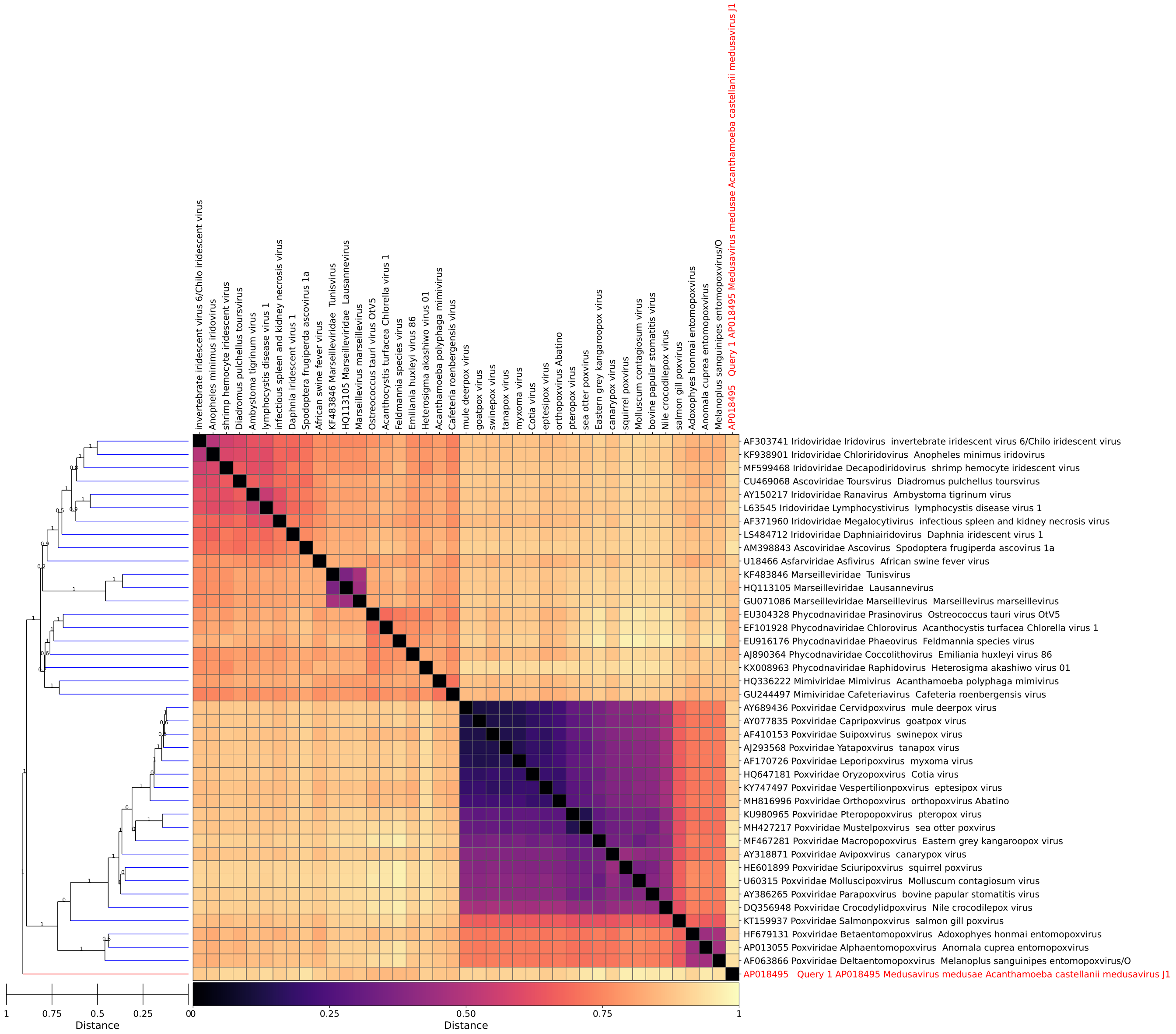


Figure 25: *Mamonoviridae*, GRAViTy-V2 heatmap

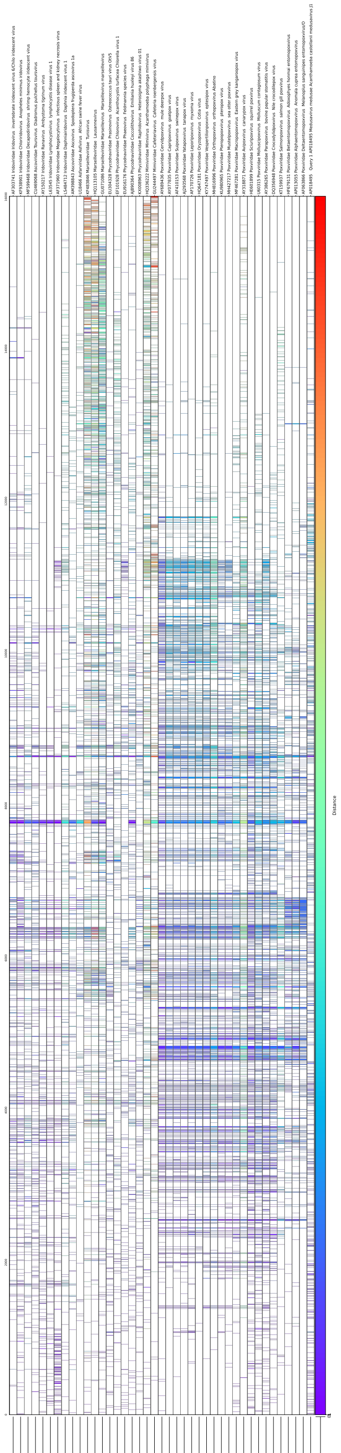

Figure 26: *Mamonoviridae*, GRAViTy-V2 barcode

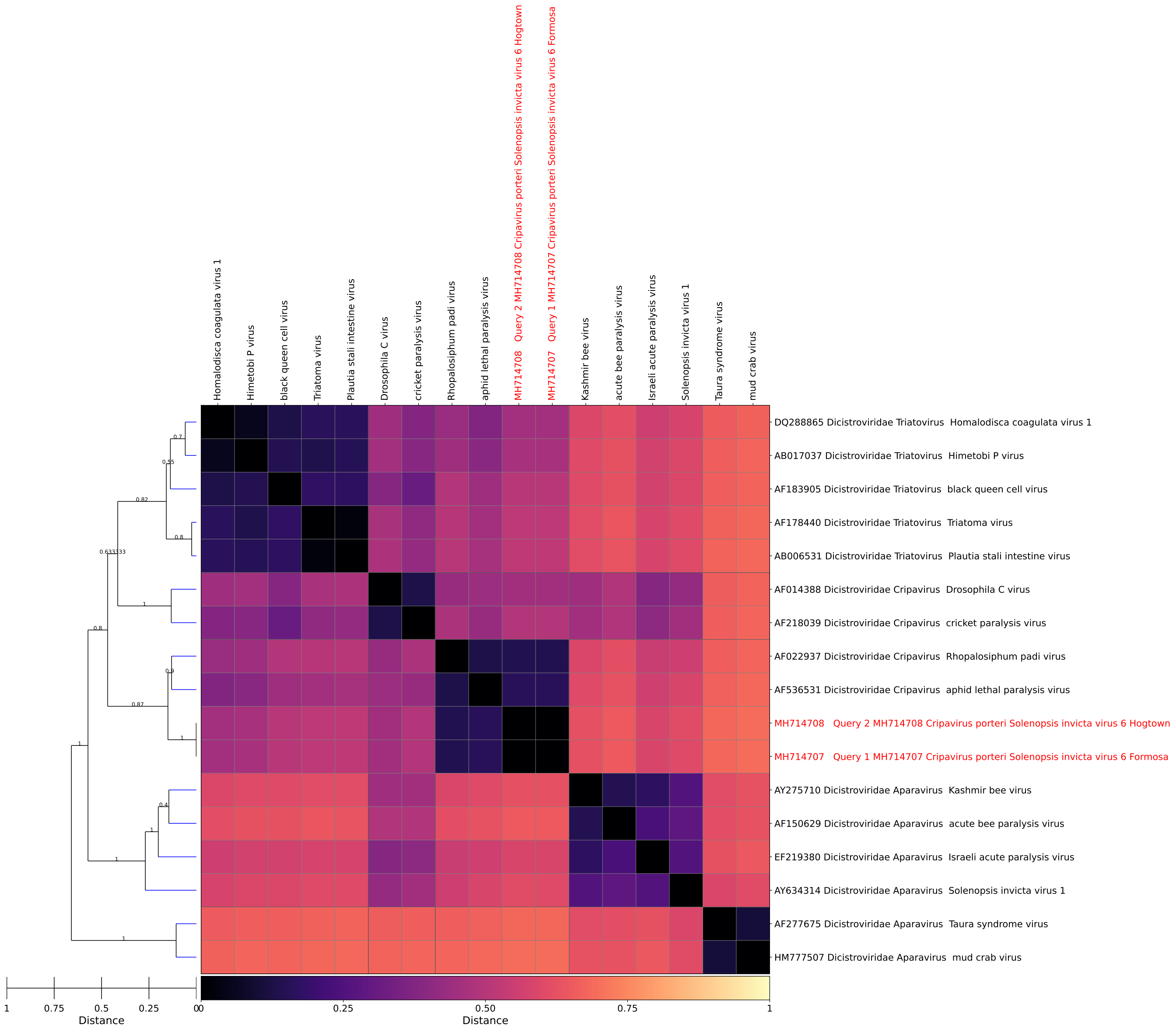


Figure 27: *Crispavirus*, GRAViTy-V2 heatmap

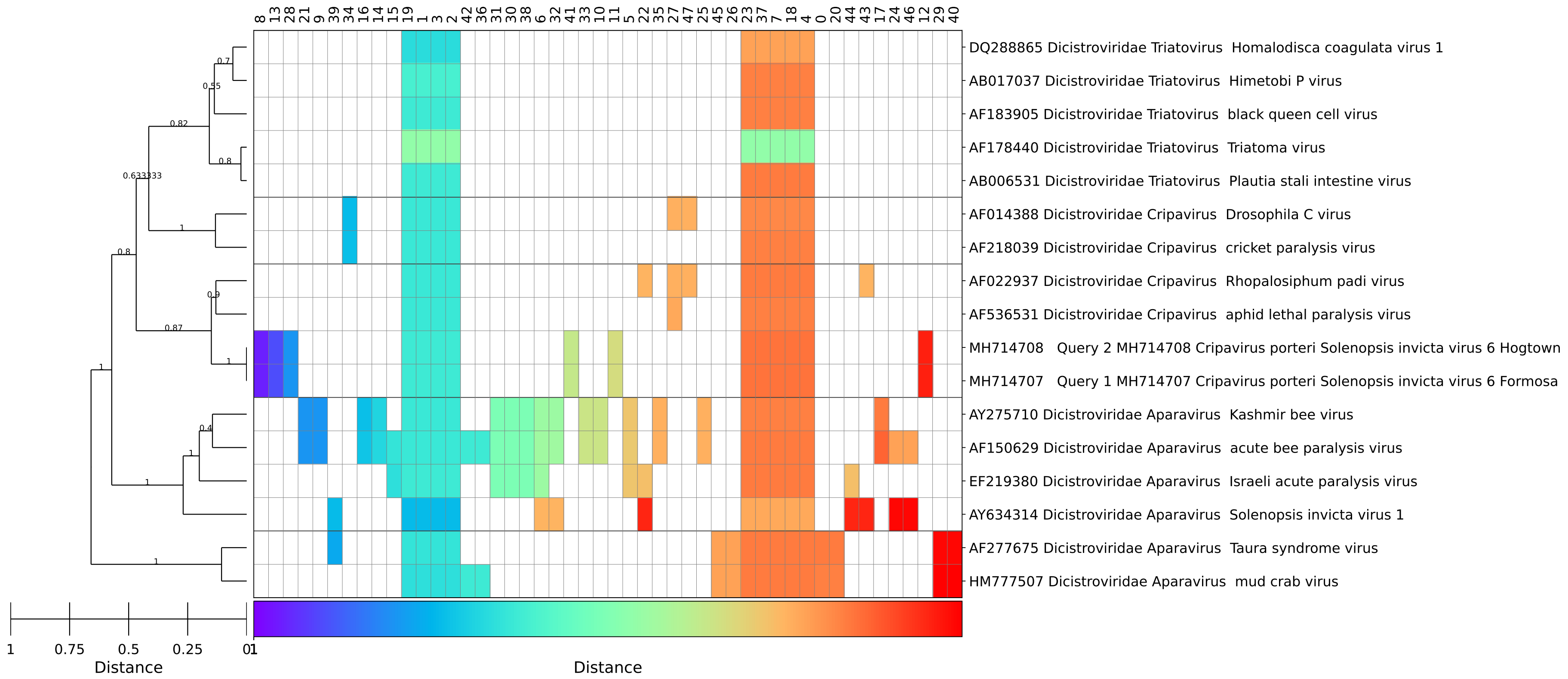


Figure 28: *Crispavirus*, GRAViTy-V2 barcode

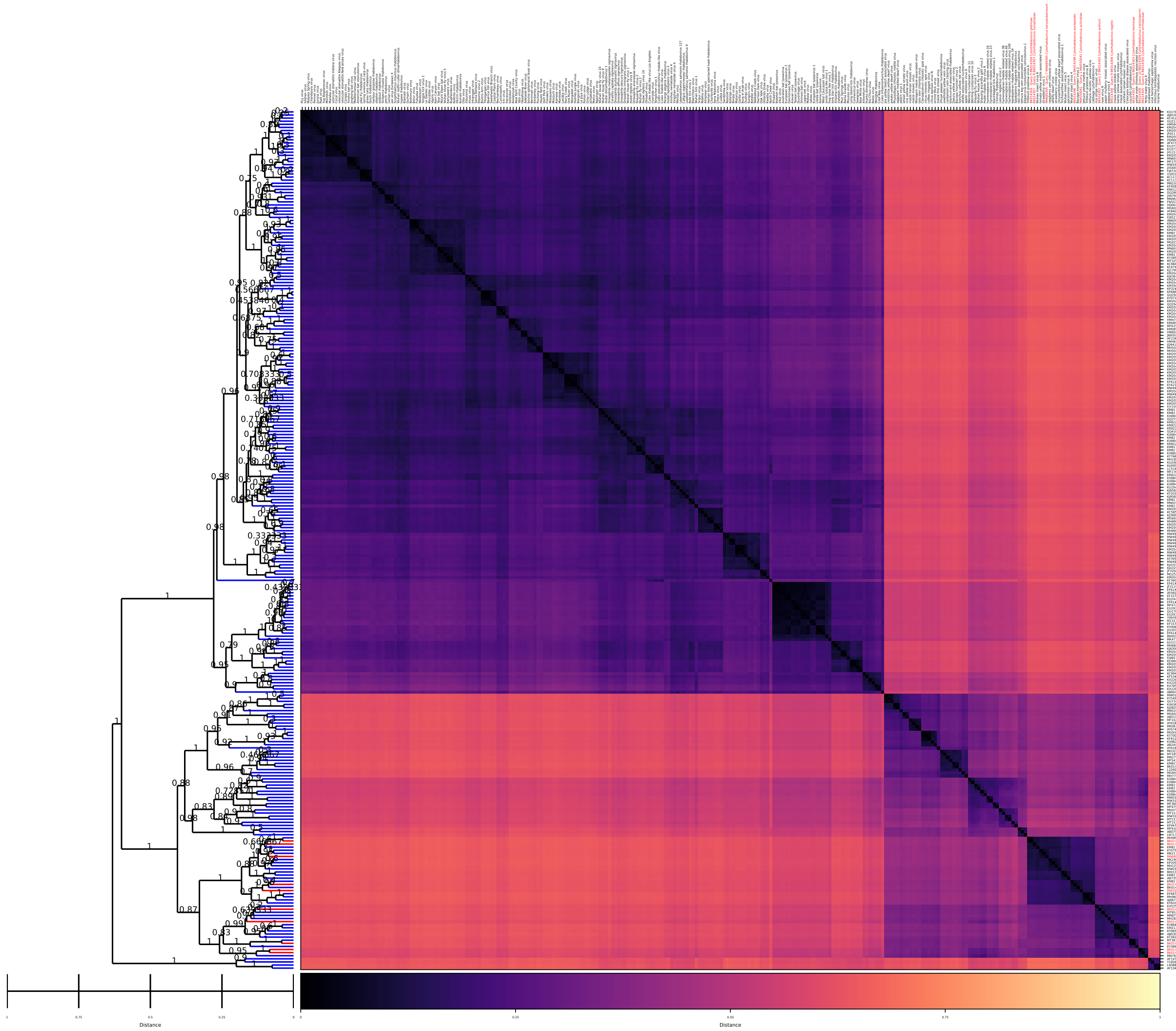


Figure 29: *Cytorhabdovirus*, GRAViTy-V2 heatmap

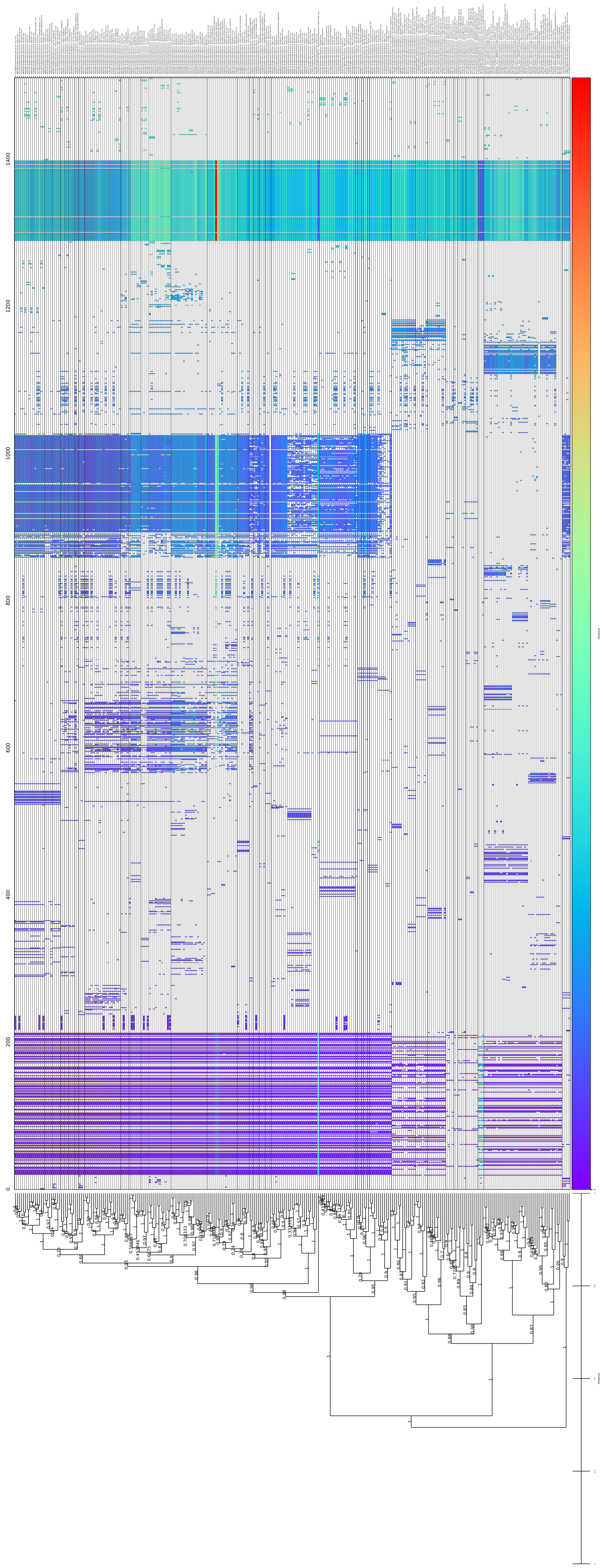

Figure 30: *Cytorhabdovirus*, GRAViTy-V2 barcode

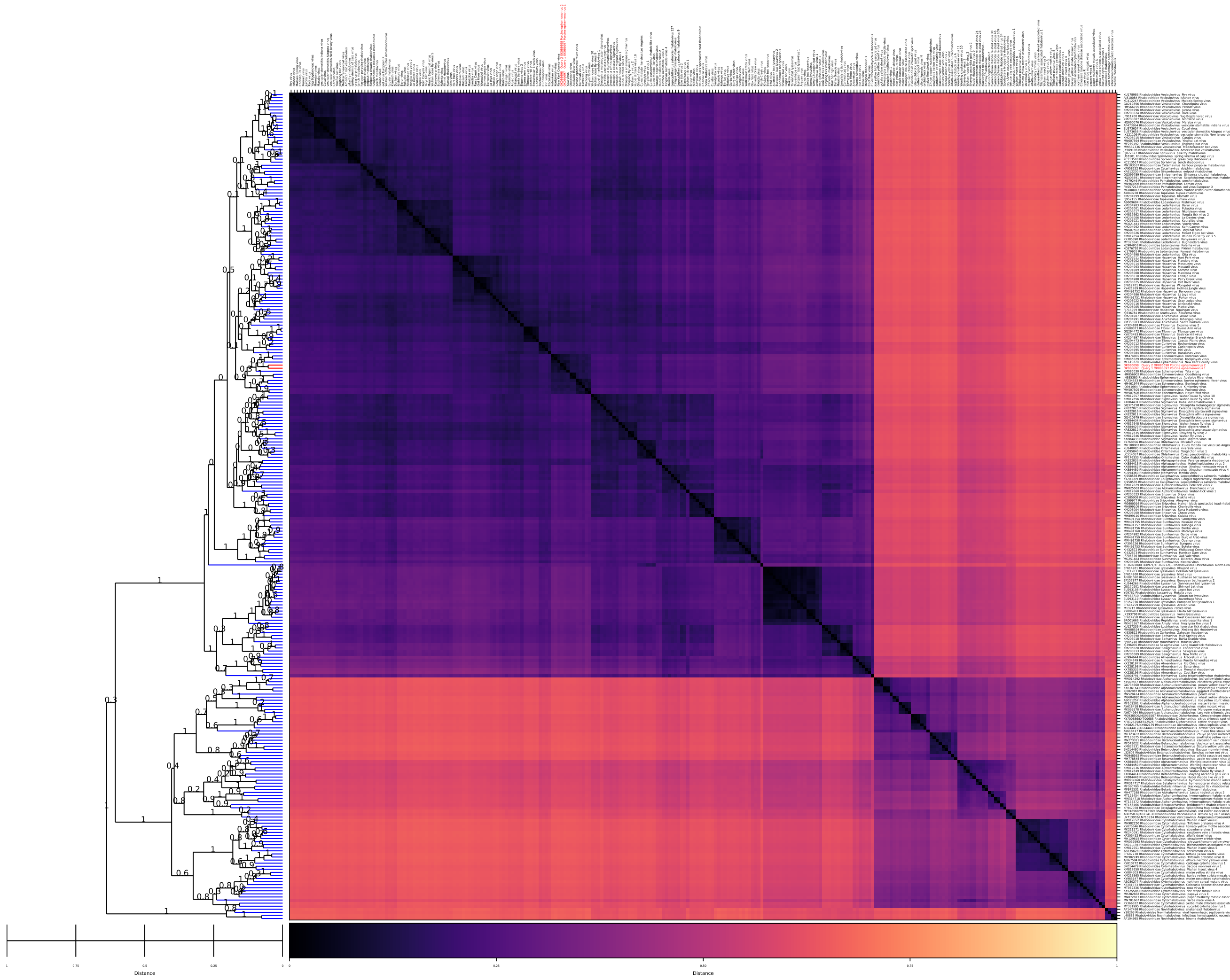

Figure 31: *Ephemerovirus*, GRAViTy-V2 heatmap

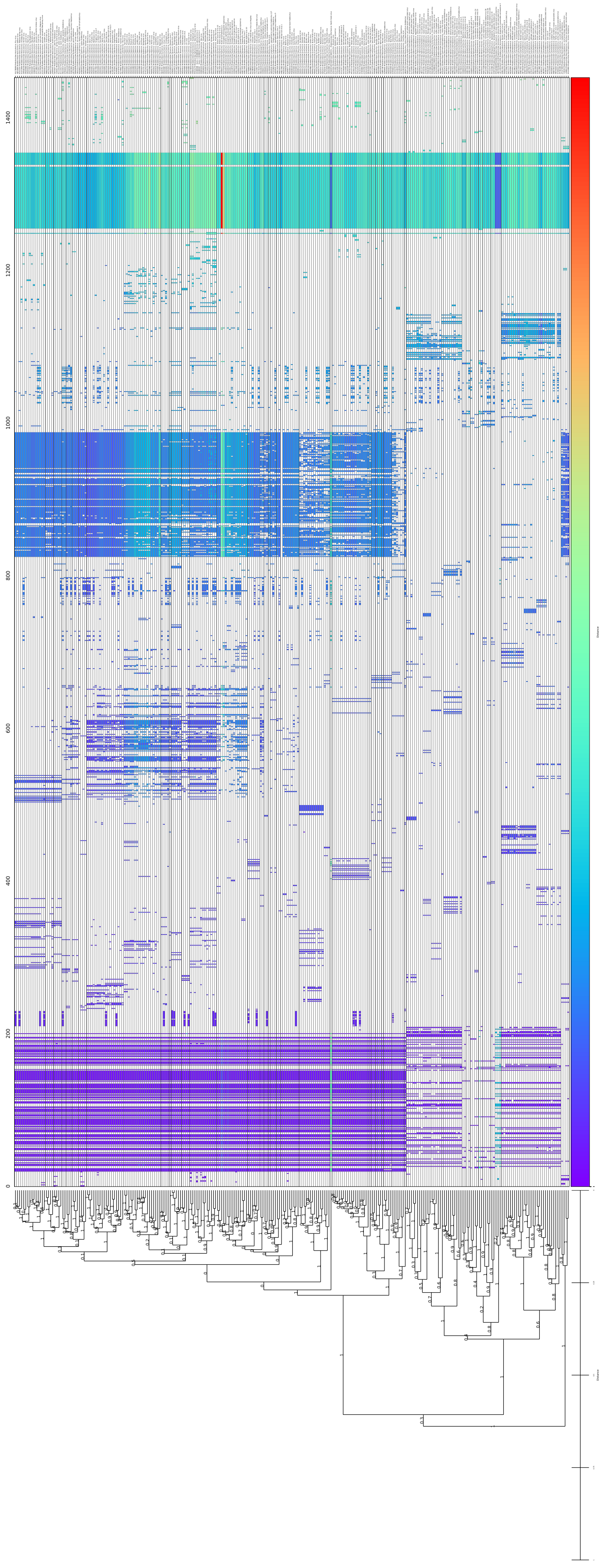

Figure 32: *Ephemerovirus*, GRAViTy-V2 barcode

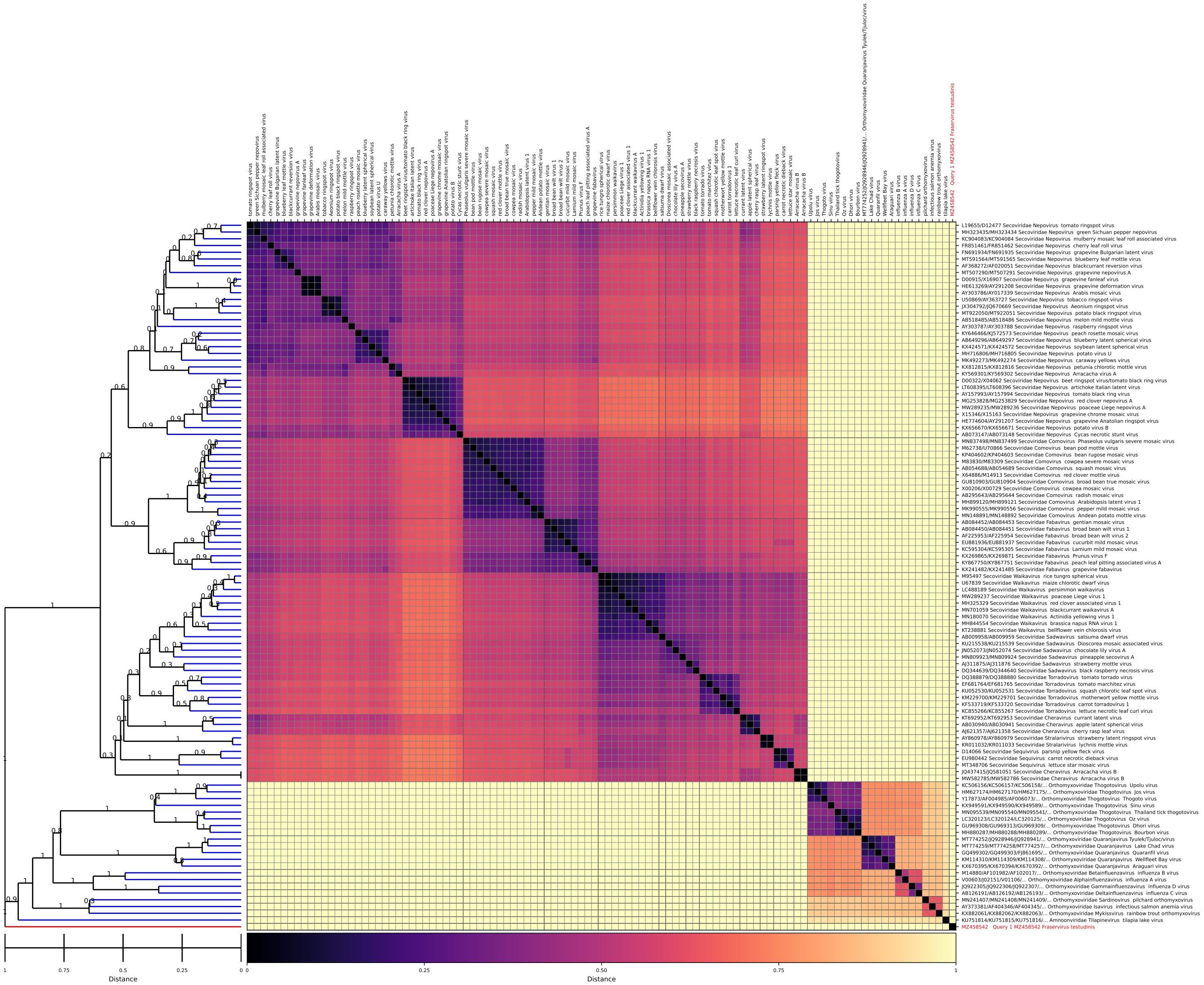


Figure 33: *Fraservirus*, GRAViTy-V2 heatmap

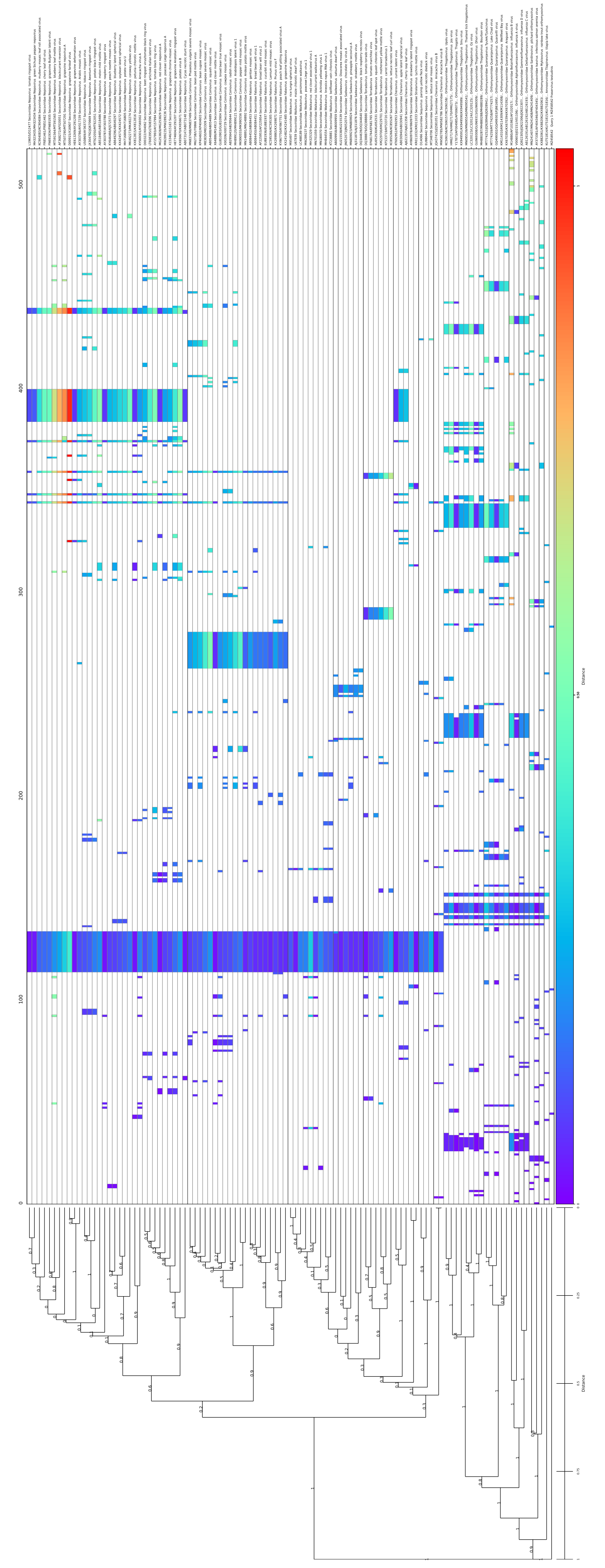

Figure 34: *Fraservirus*, GRAViTy-V2 barcode

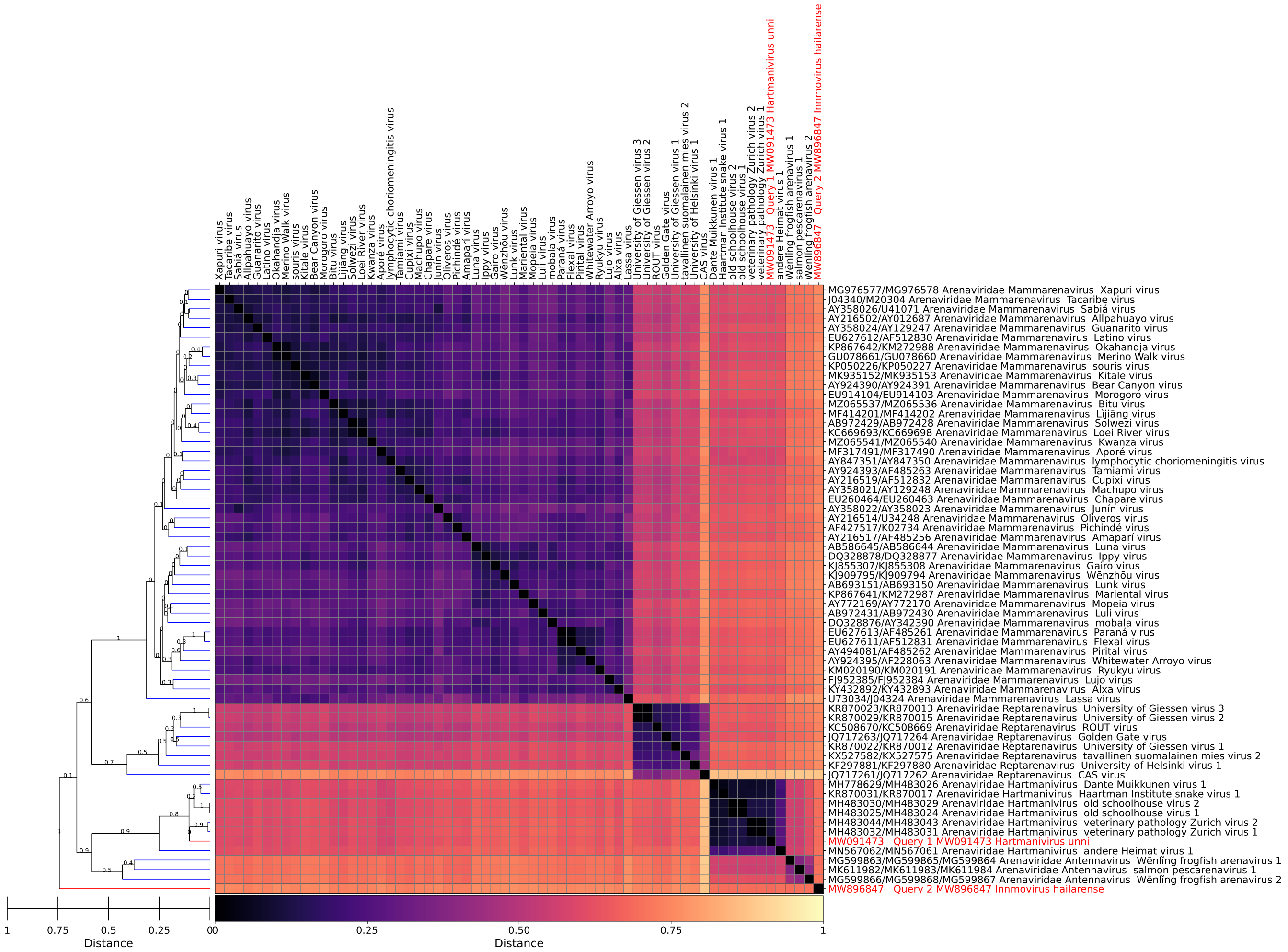

Figure 35: *Hartmanivirus*, GRAViTy-V2 heatmap

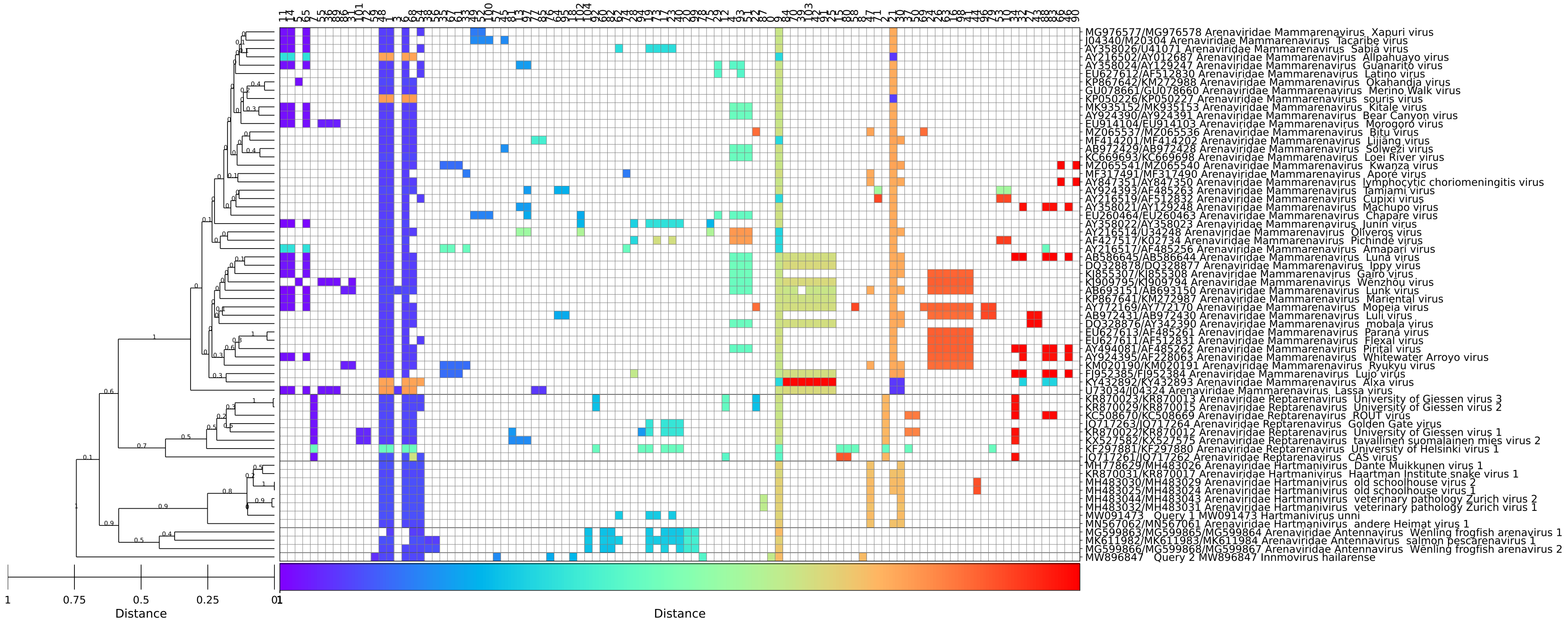

Figure 36: *Hartmanivirus*, GRAViTy-V2 barcode

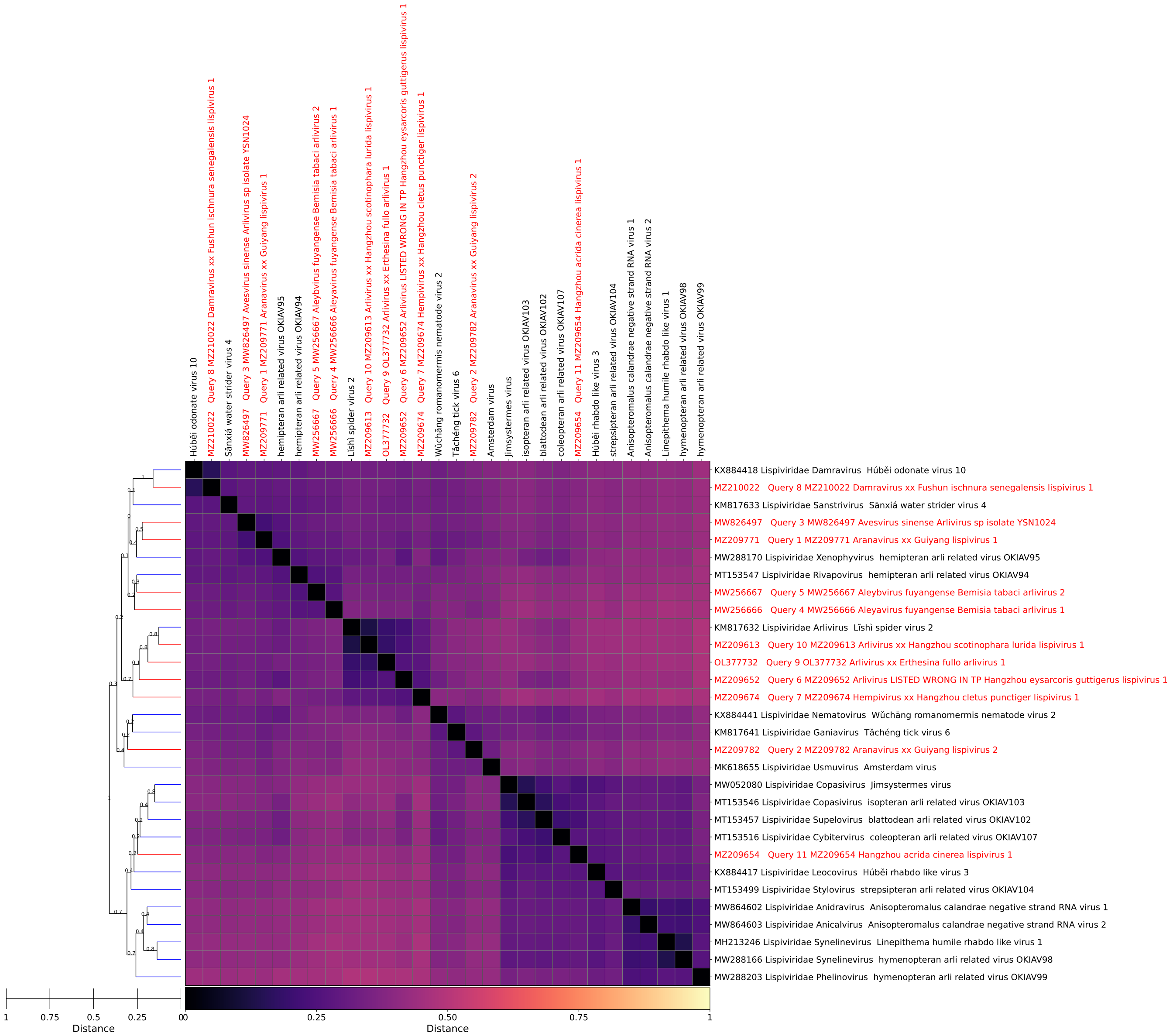

Figure 37: *Lispiviridae*, GRAViTy-V2 heatmap

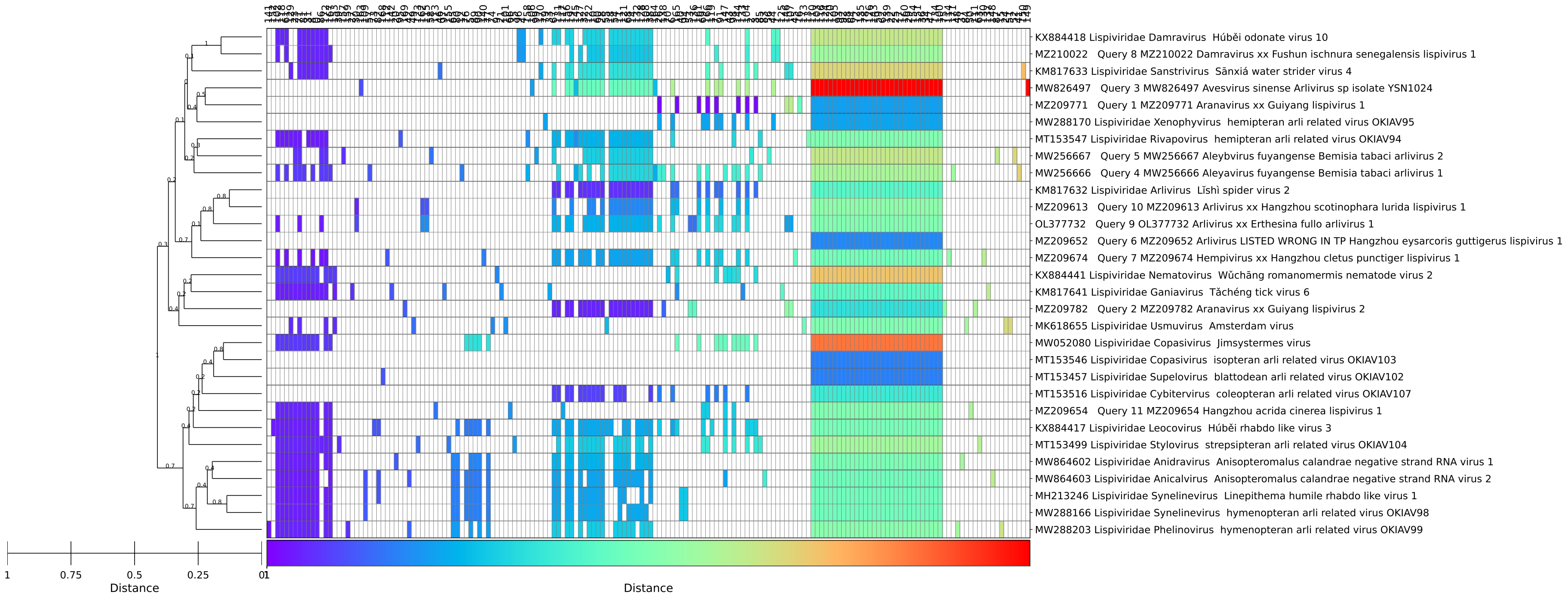


Figure 38: *Lispiviridae*, GRAViTy-V2 barcode

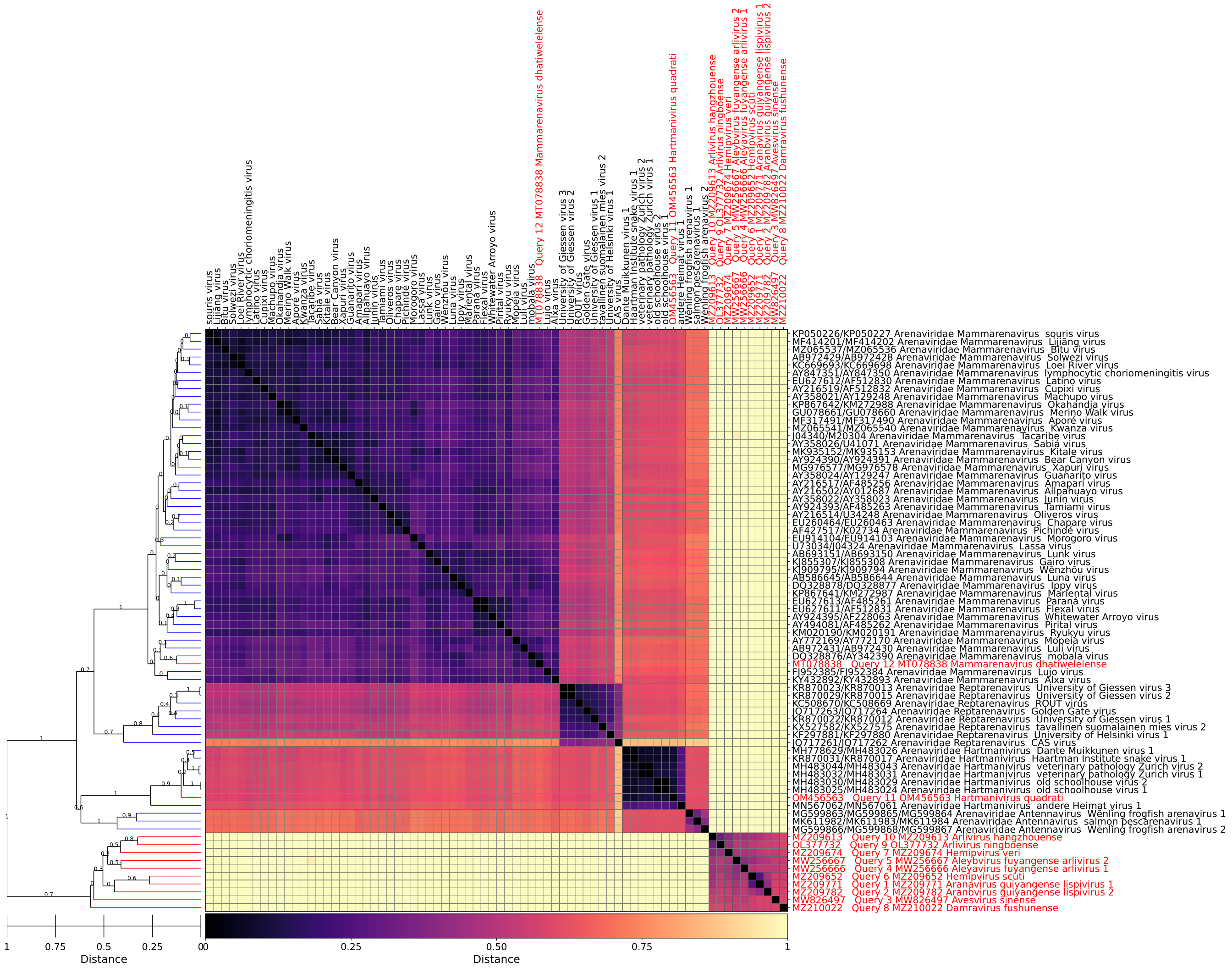


Figure 39: *Mammarenavirus*, GRAViTy-V2 heatmap

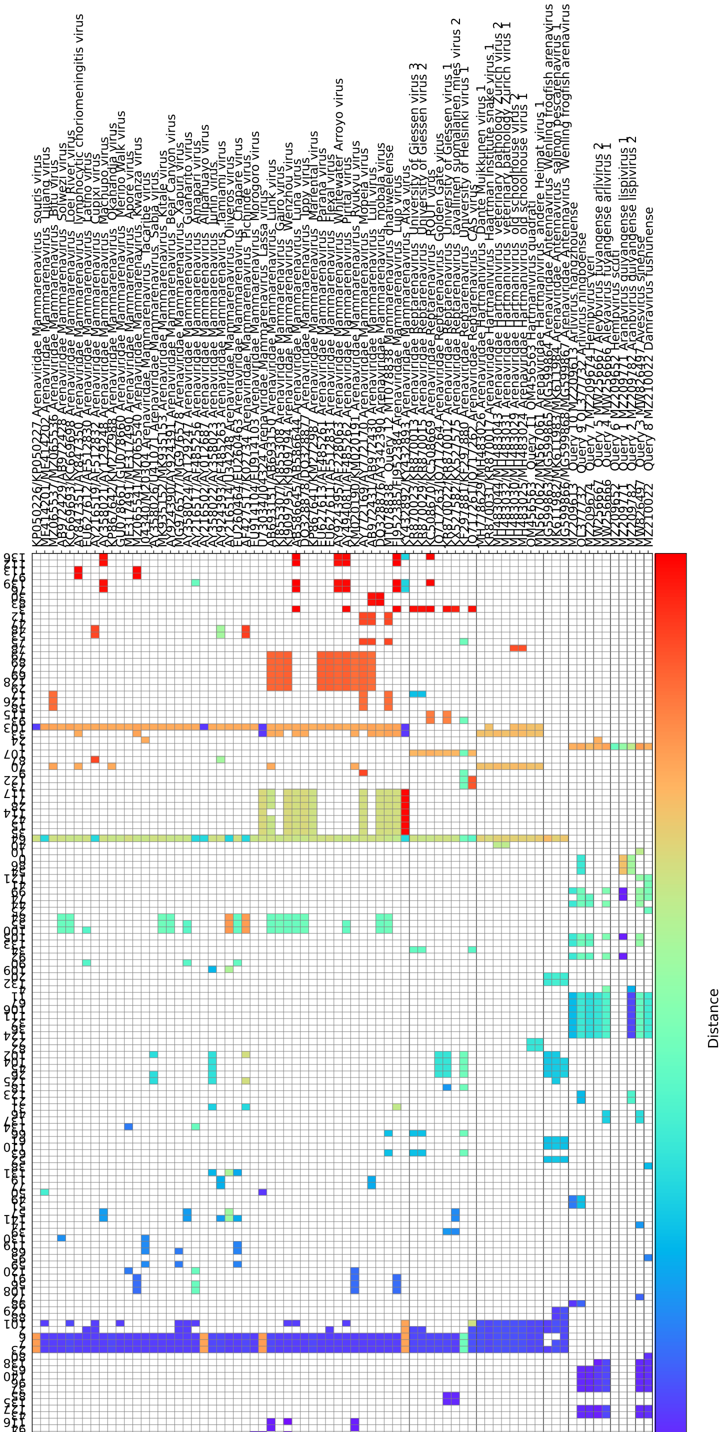

Figure 40: *Mammarenavirus*, GRAViTy-V2 barcode

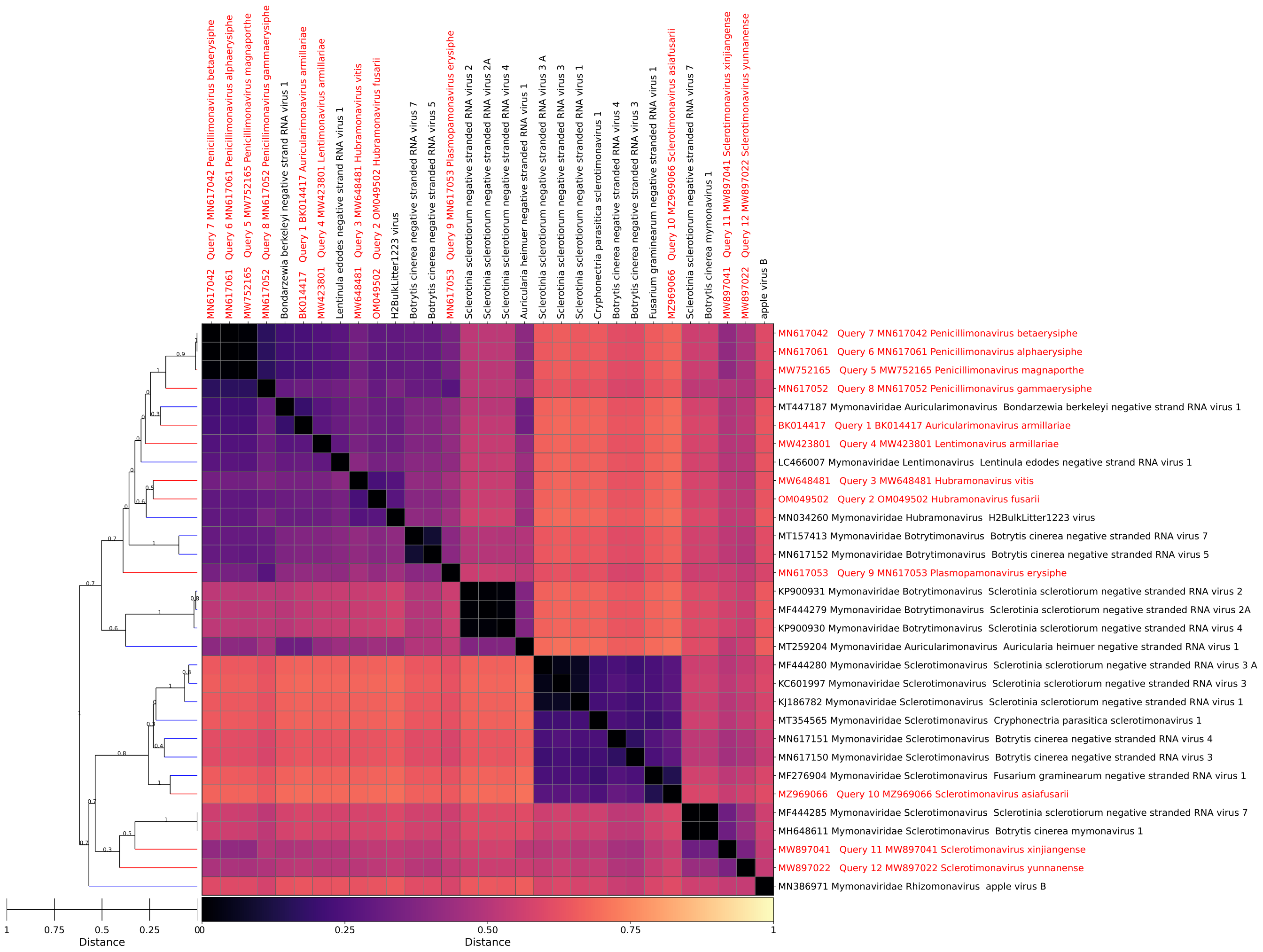


Figure 41: *Mymonaviridae*, GRAViTy-V2 heatmap

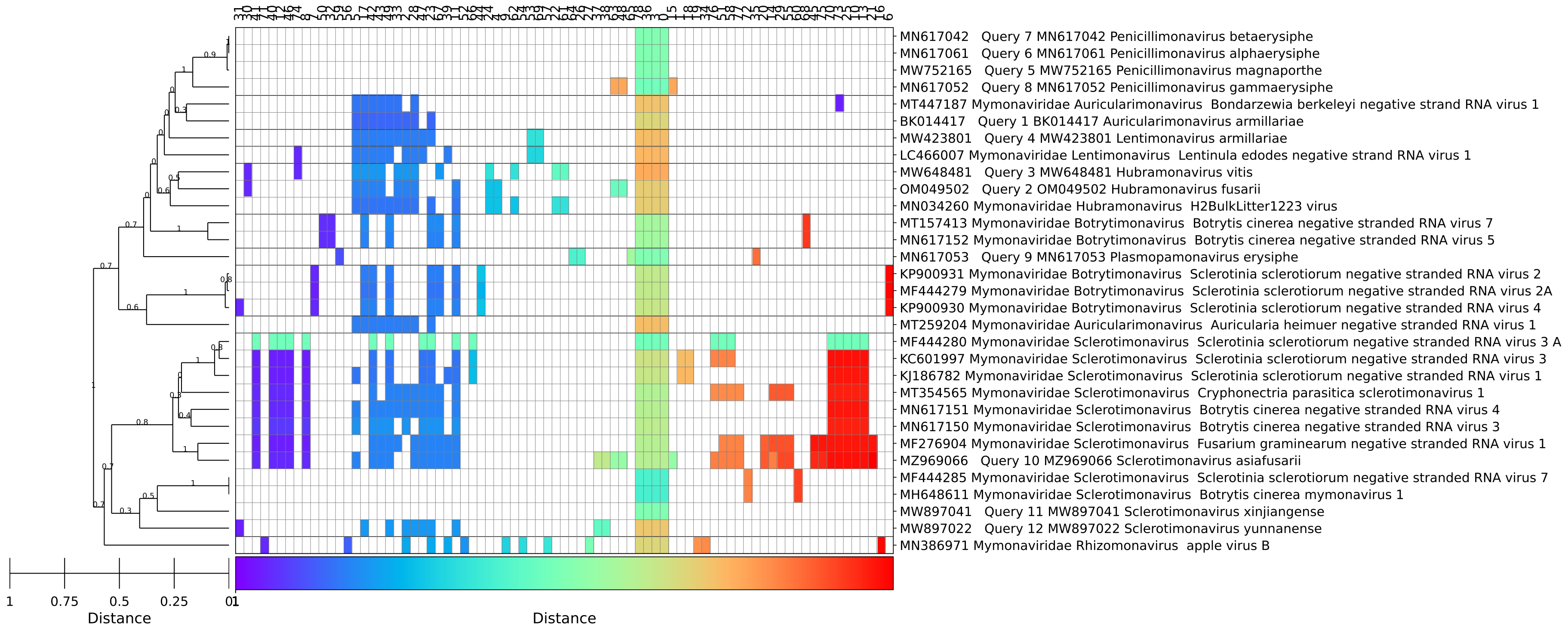


Figure 42: *Mymonaviridae*, GRAViTy-V2 barcode

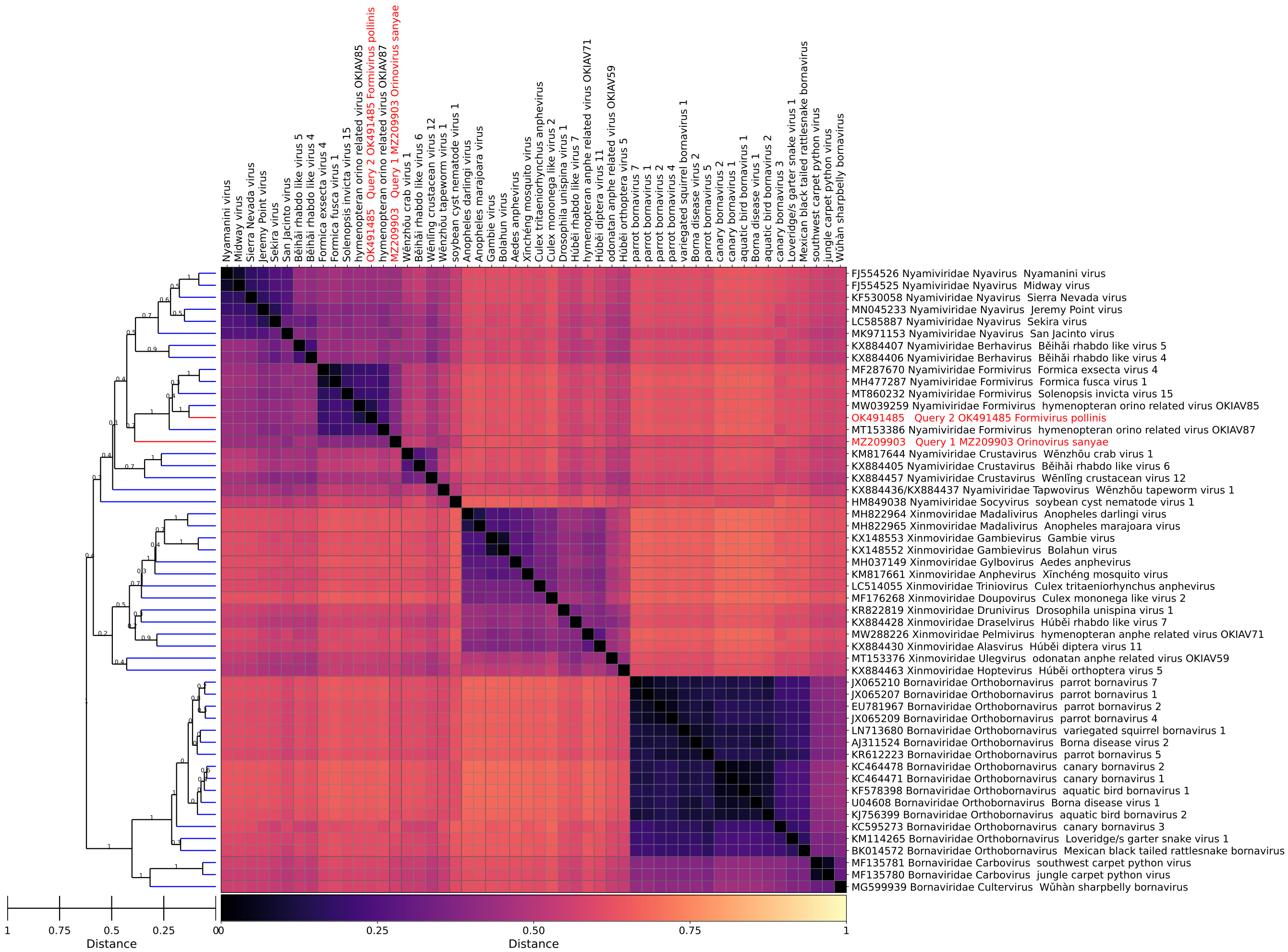


Figure 43: *Nyamiviridae*, GRAViTy-V2 heatmap

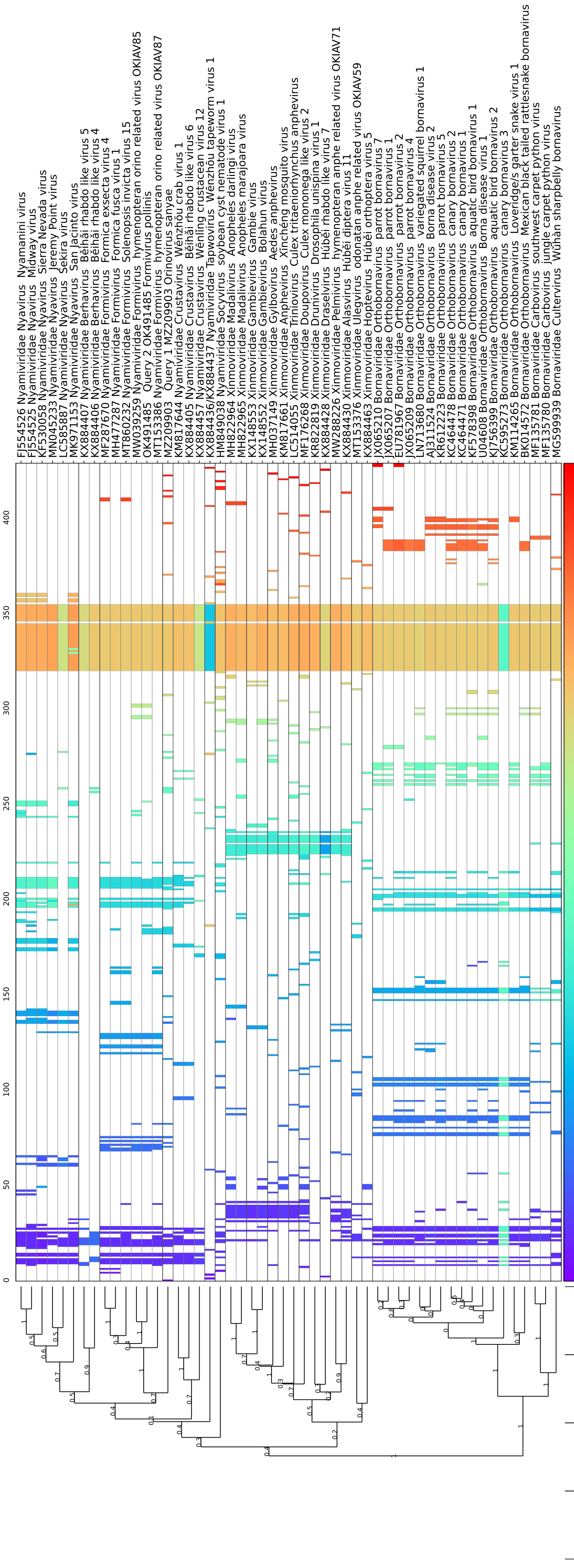


Figure 44: *Nyamiviridae*, GRAViTy-V2 barcode

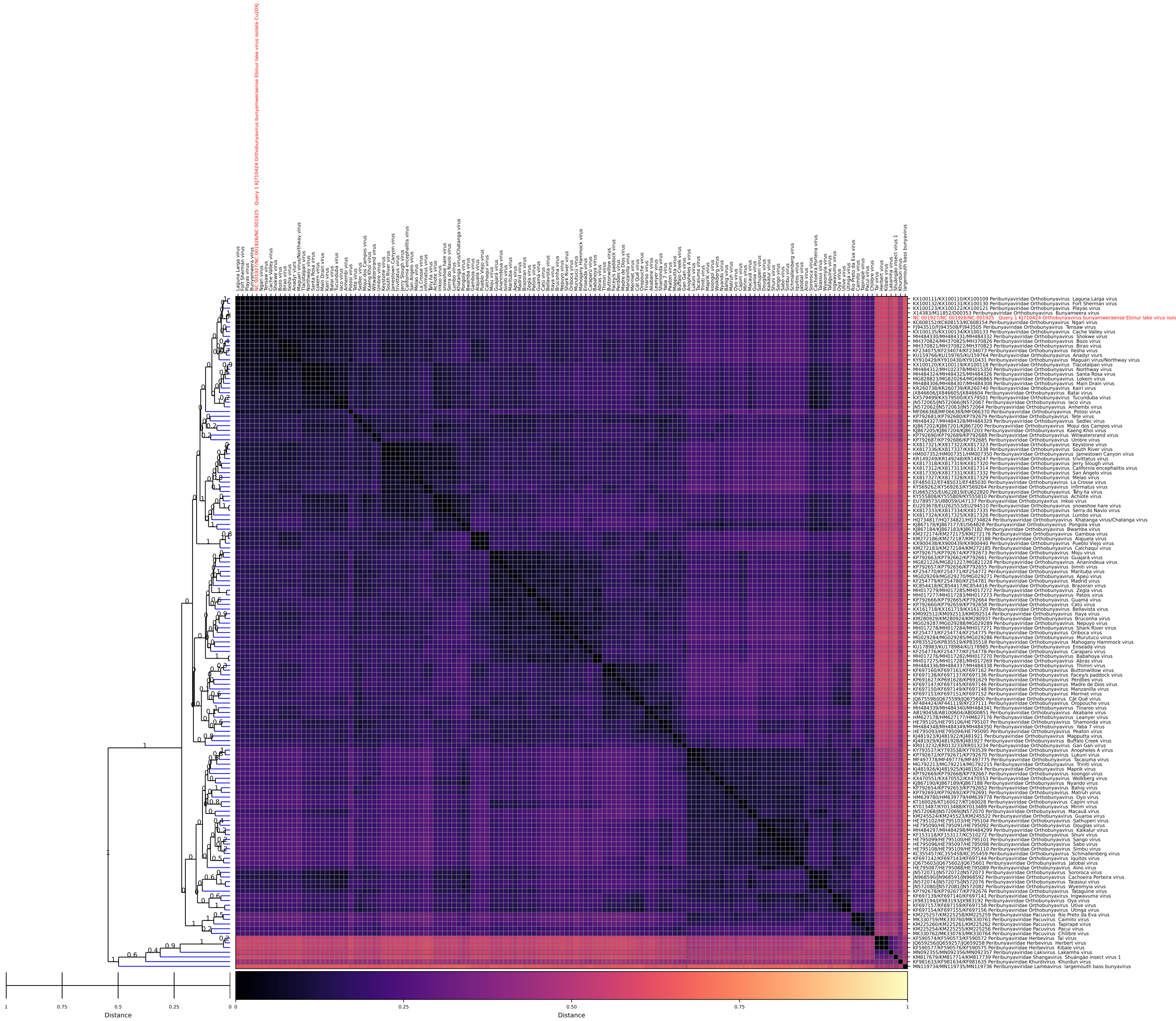

Figure 45: *Orthobunyavirus* (017M), GRAViTy-V2 heatmap

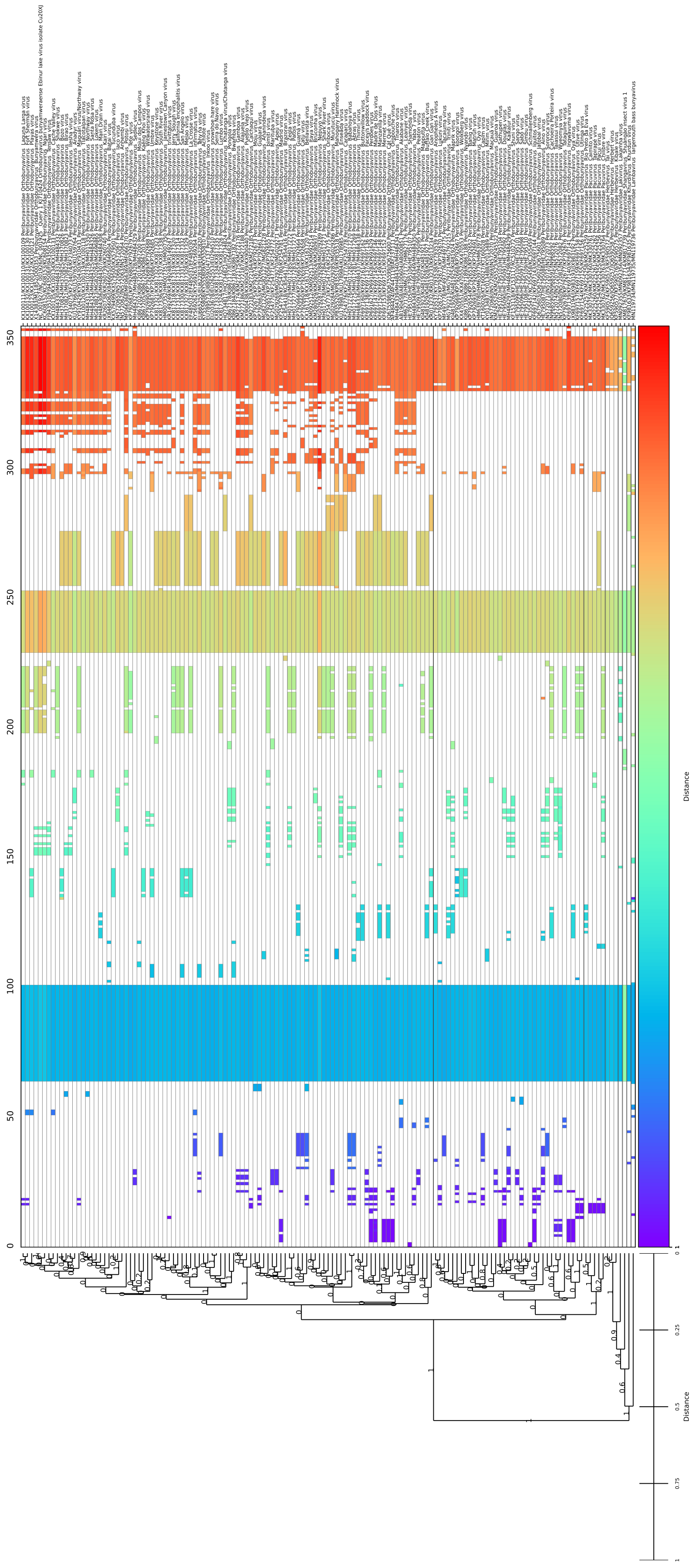


Figure 46: *Orthobunyavirus* (017M), GRAViTy-V2 barcode

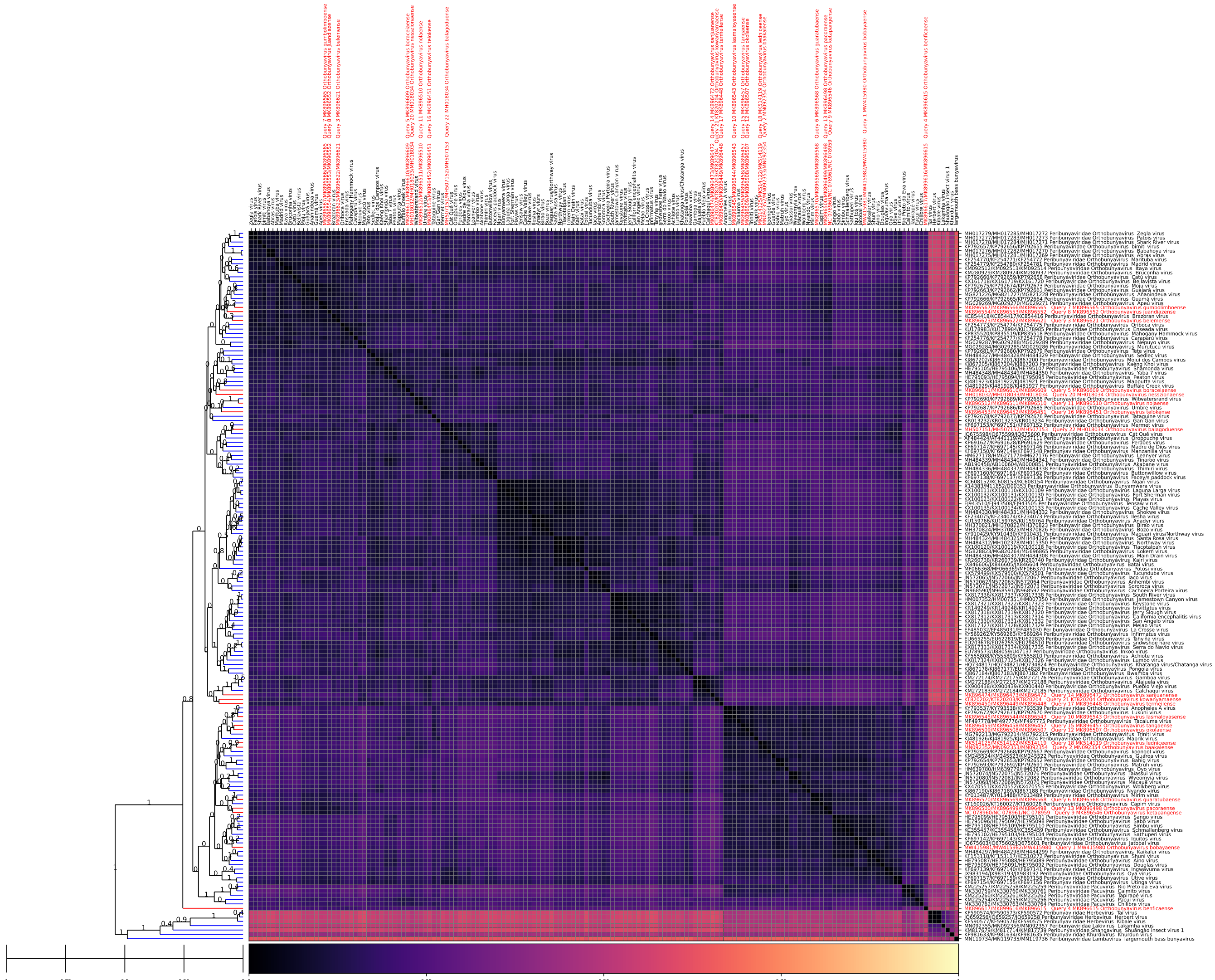


Figure 47: *Orthobunyavirus* (018M), GRAViTy-V2 heatmap

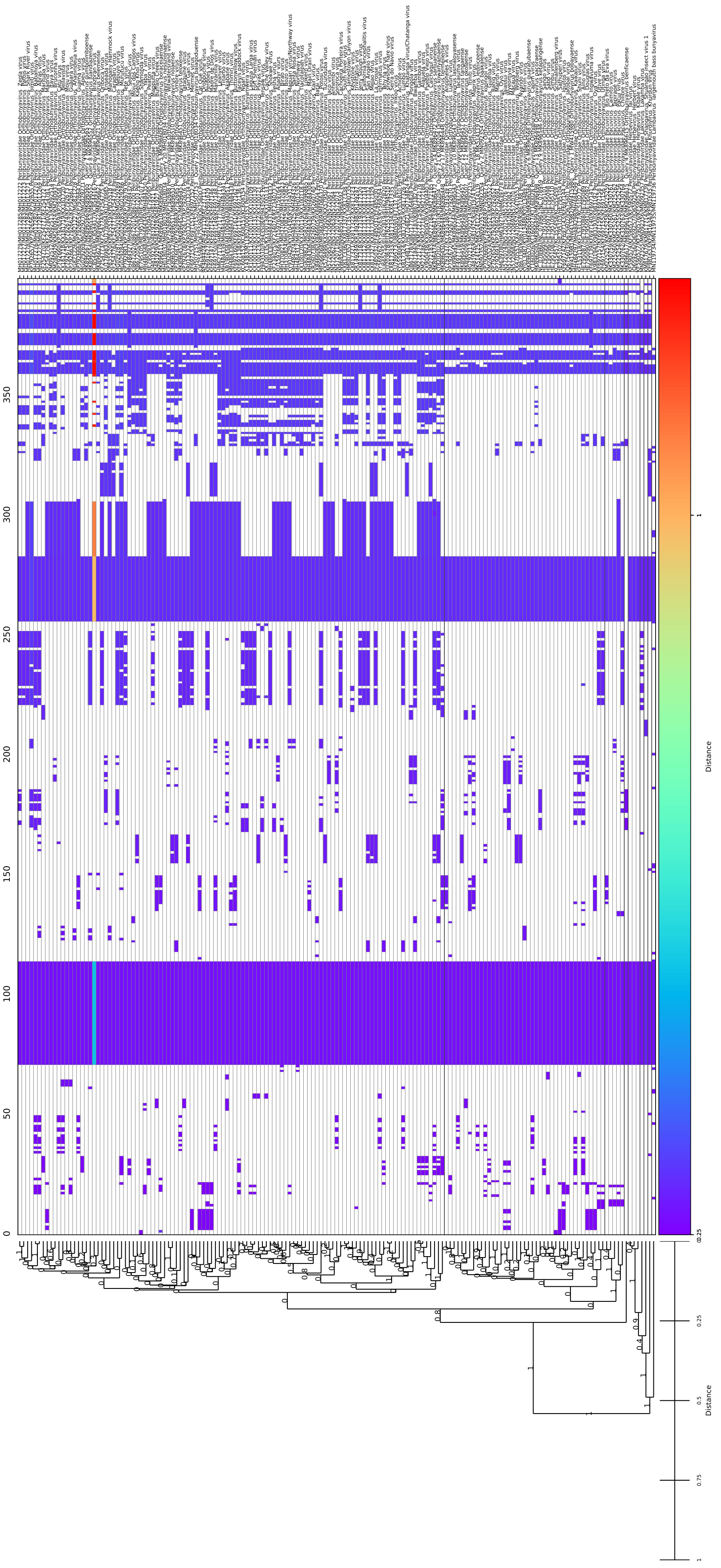

Figure 48: *Orthobunyavirus* (018M), GRAViTy-V2 barcode

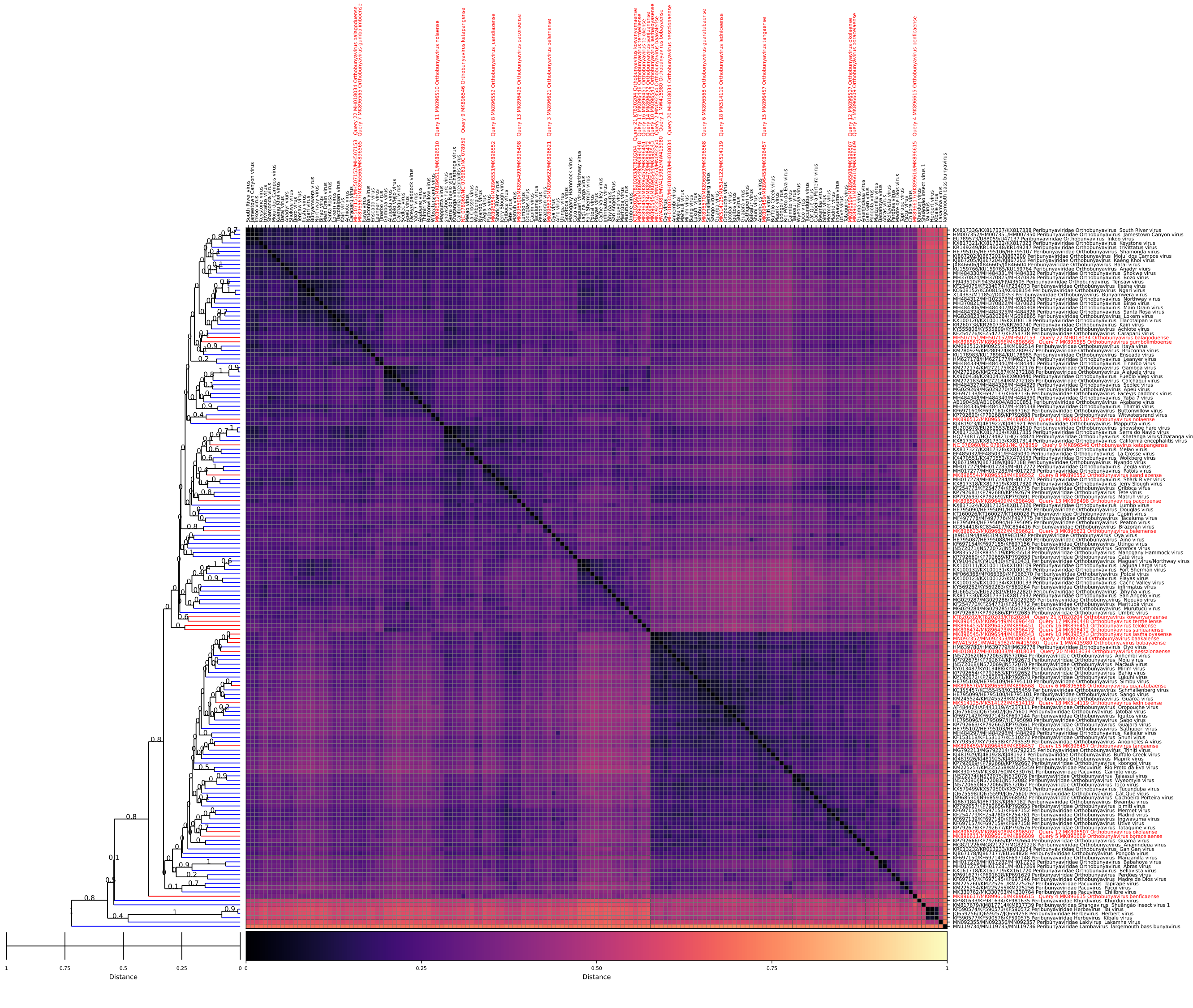


Figure 49: *Phasmaviridae*, GRAViTy-V2 heatmap

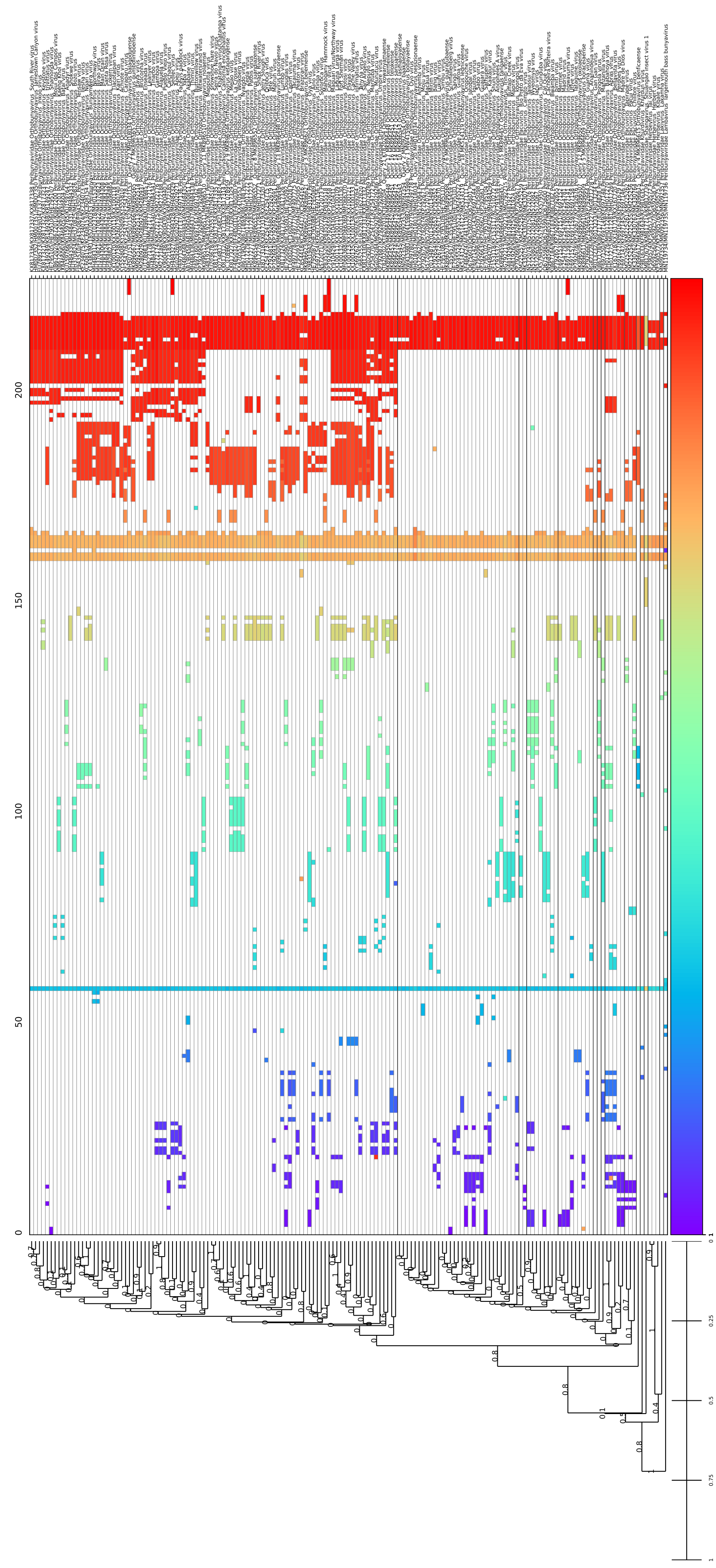


Figure 50: *Phasmaviridae*, GRAViTy-V2 barcode

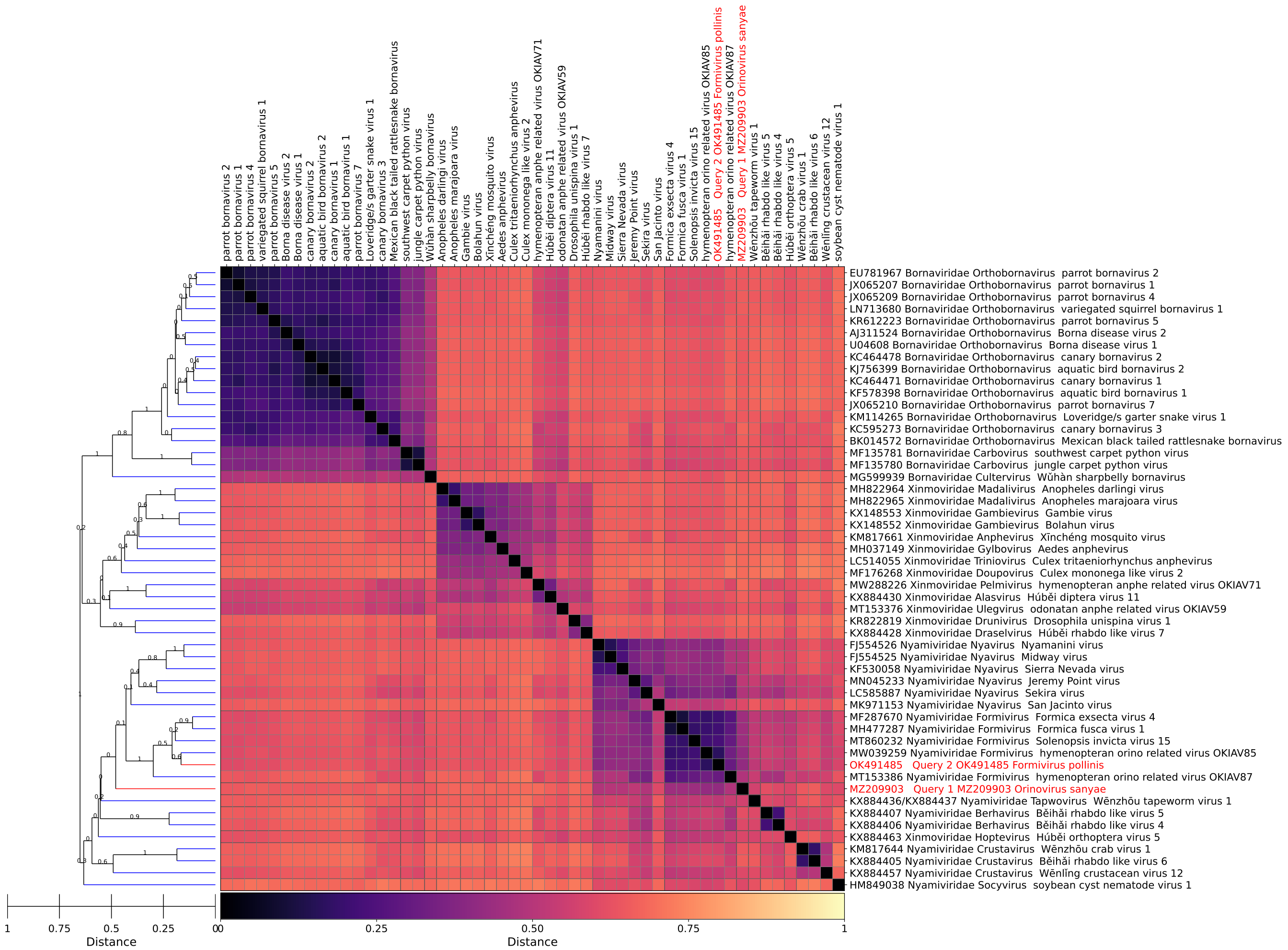


Figure 51: *Phenuiviridae*, GRAViTy-V2 heatmap

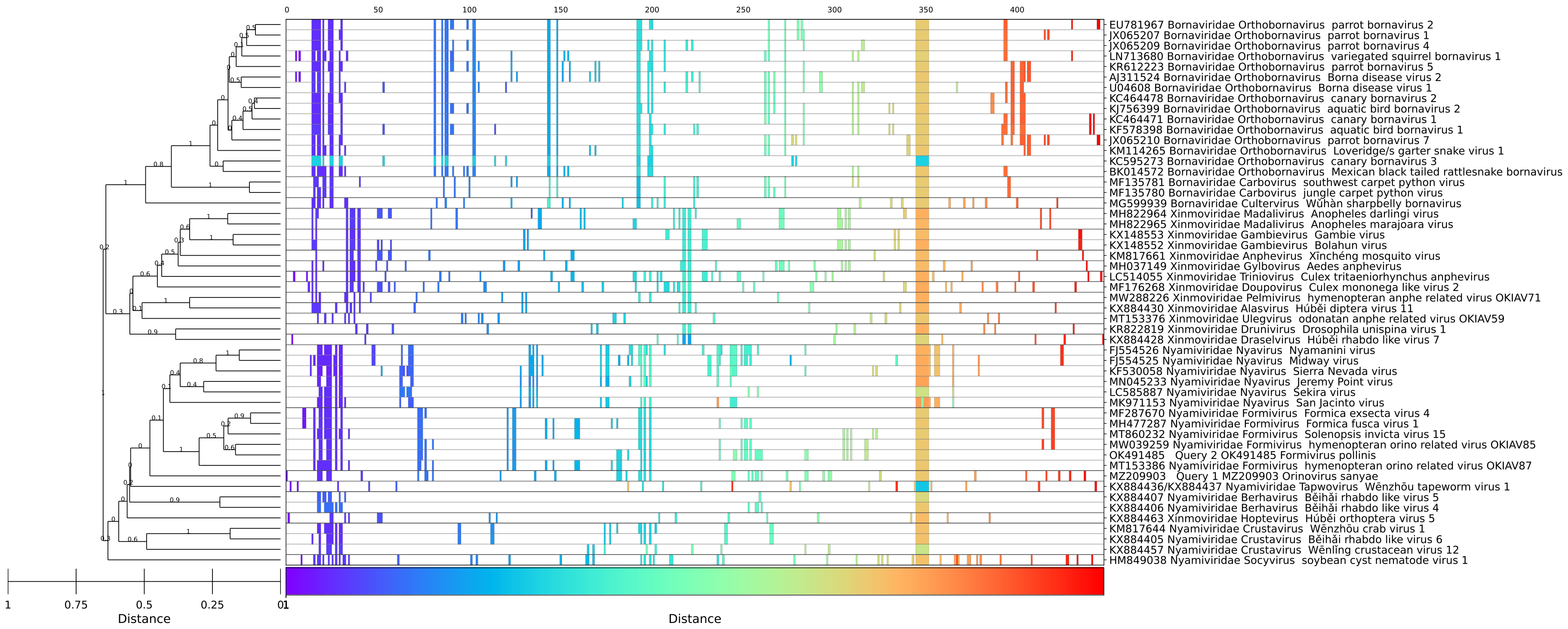


Figure 52: *Phenuiviridae*, GRAViTy-V2 barcode

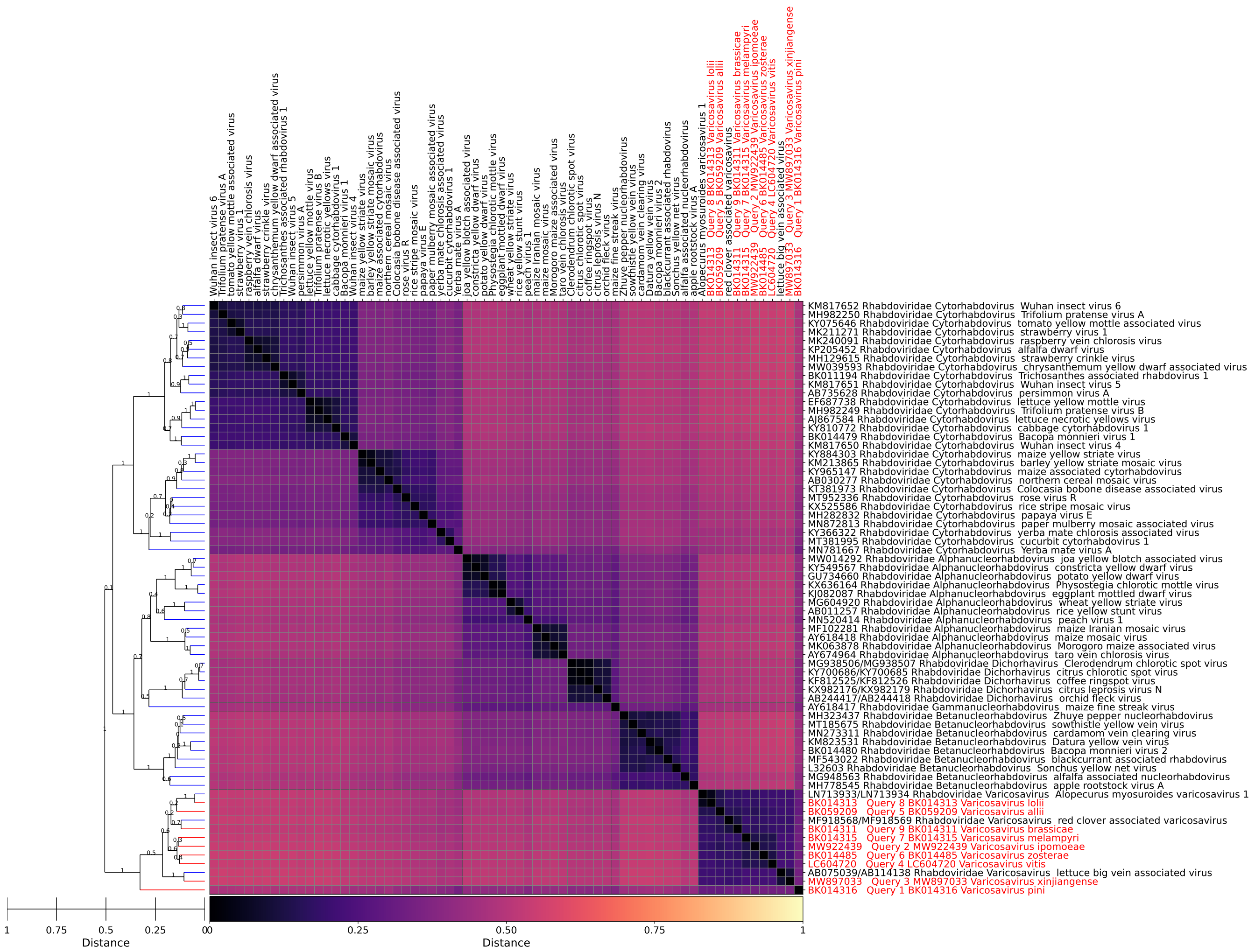


Figure 53: *Varicosavirus*, GRAViTy-V2 heatmap

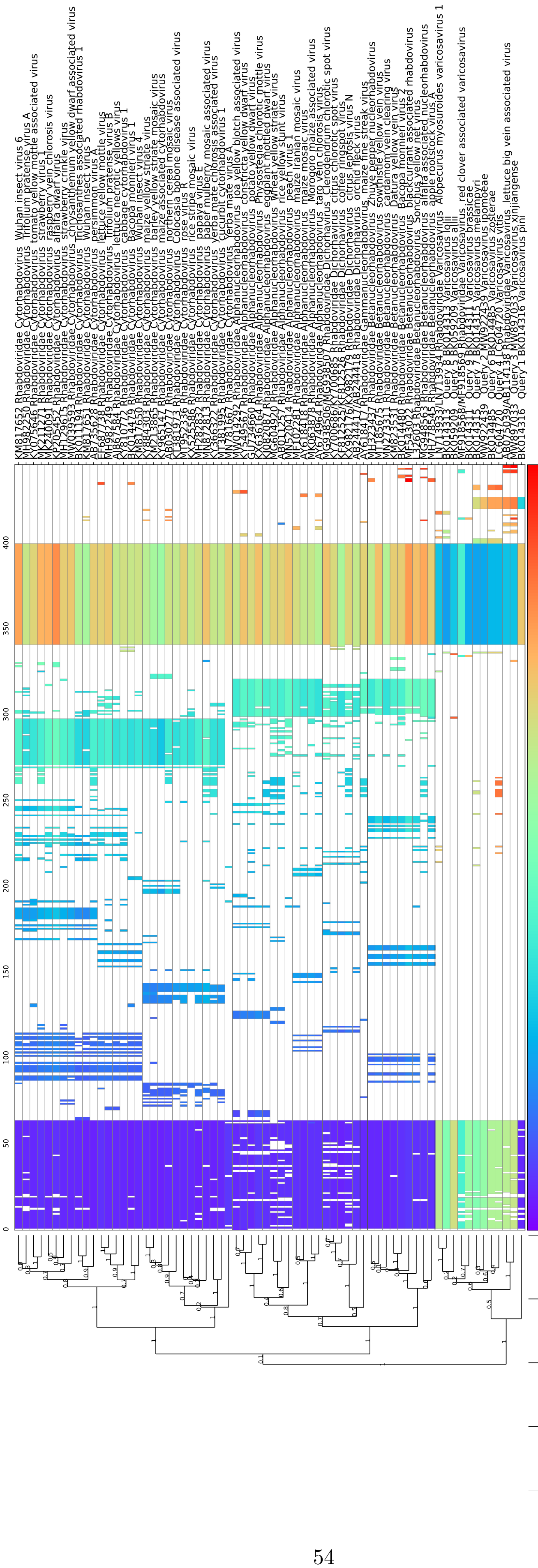


Figure 54: *Varicosavirus*, GRAViTy-V2 barcode

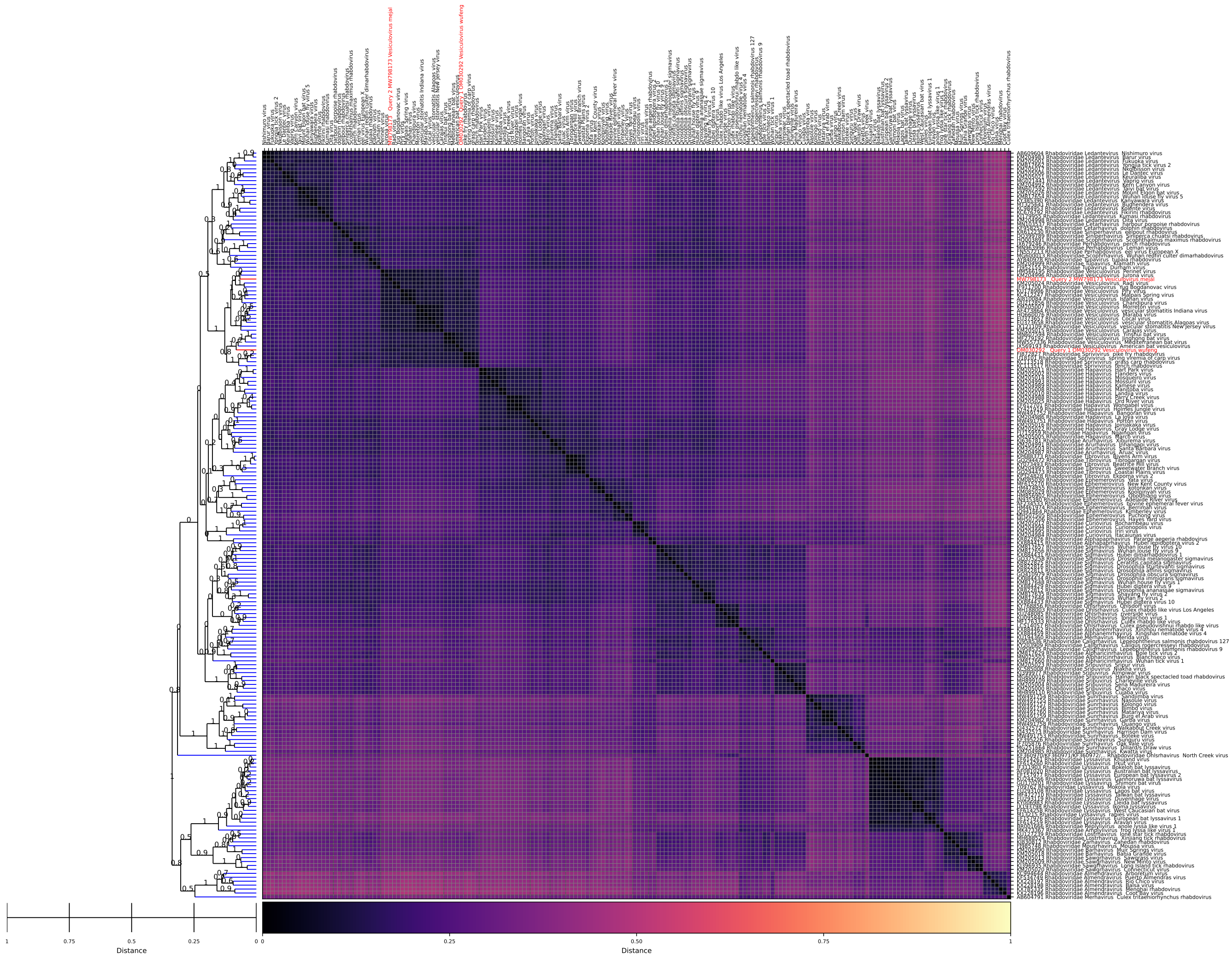

Figure 55: *Vesiculovirus*, GRAViTy-V2 heatmap

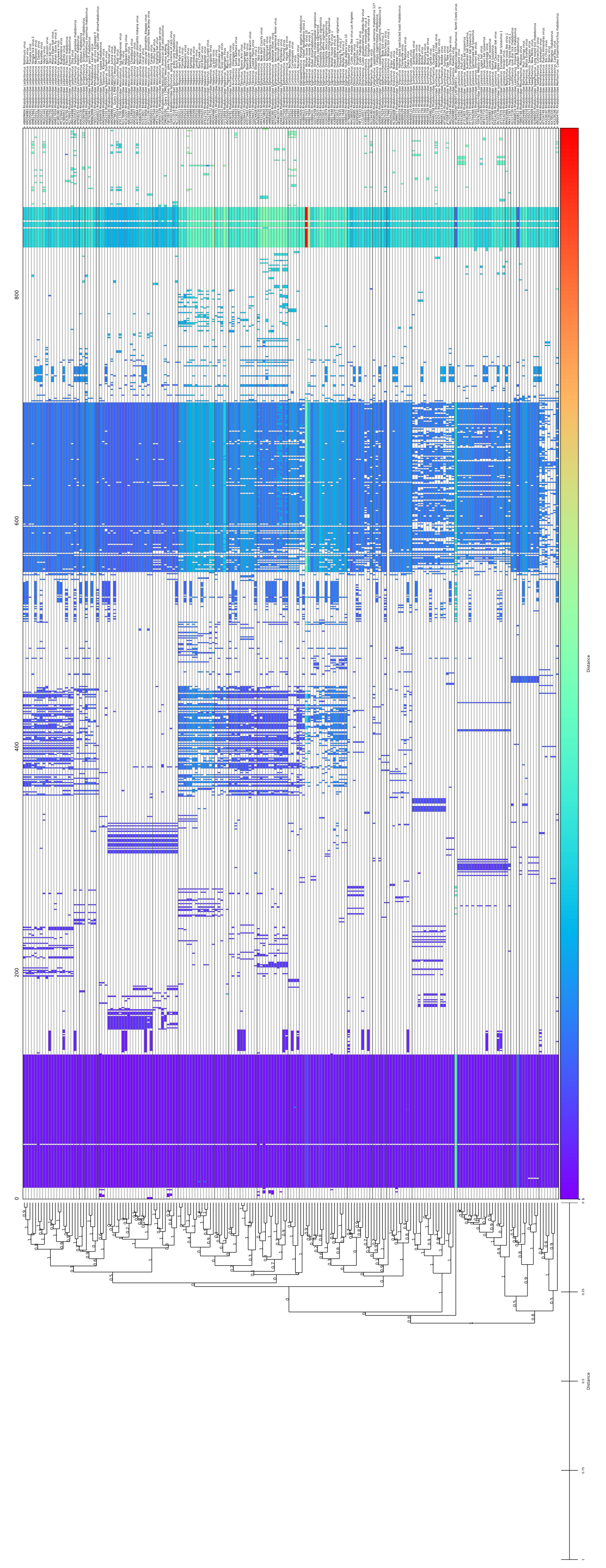

Figure 56: *Vesiculovirus*, GRAViTy-V2 barcode
