## Supplementary material for "GRAViTy-V2: a grounded viral taxonomy application": SI Document 1

Mayne, R., Aiwsakun, P., Turner, D., Adriaenssens, E. M. and Simmonds, P. (2024)

Supplementary information, Document 1

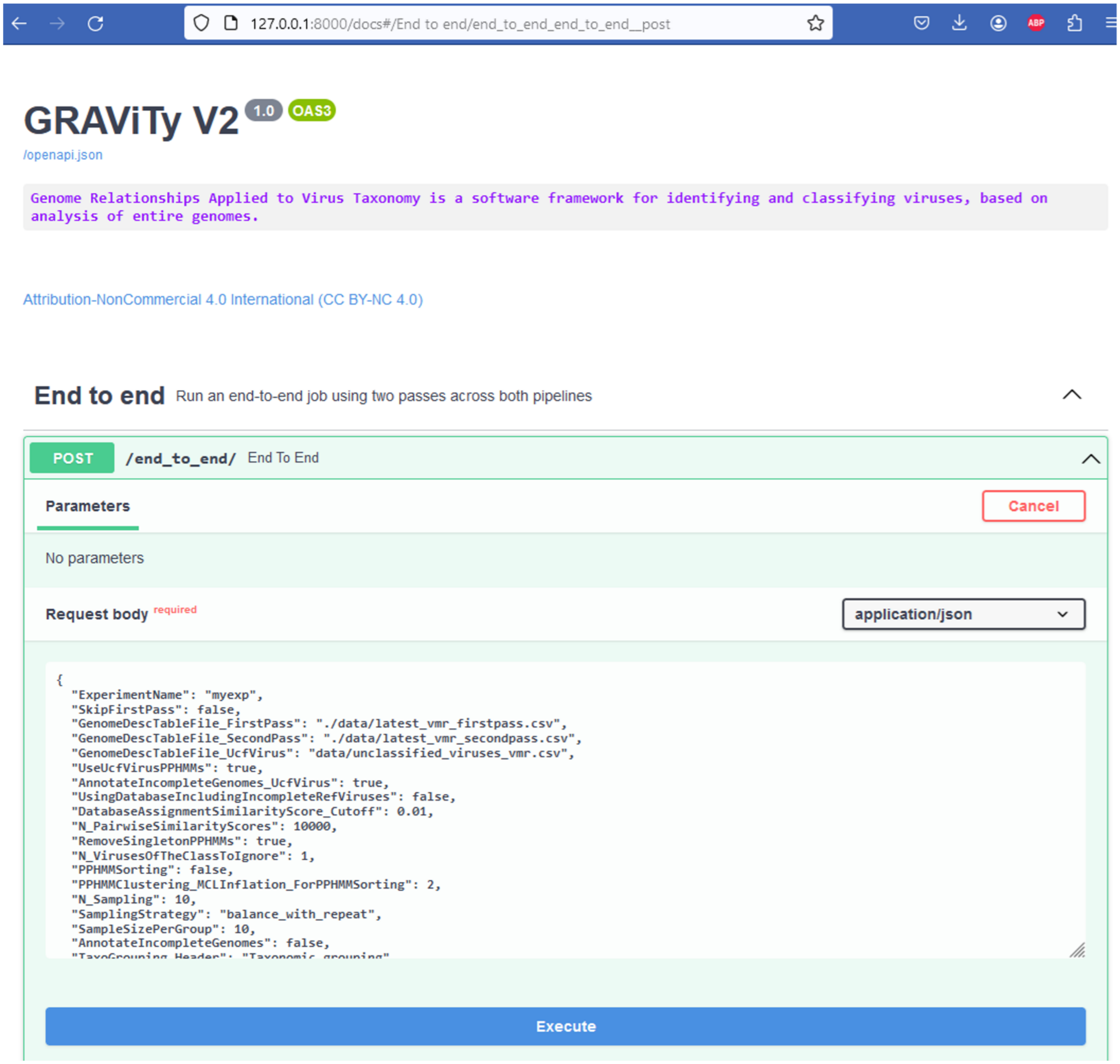


Figure 1: S1. **GRAViTy-V2's graphical user interface**, as accessed via a browser. View shows the main end-to-end endpoint (function) window expanded, revealing the dialogue box and "execute" button that may be used to refine and trigger experiments, respectively.

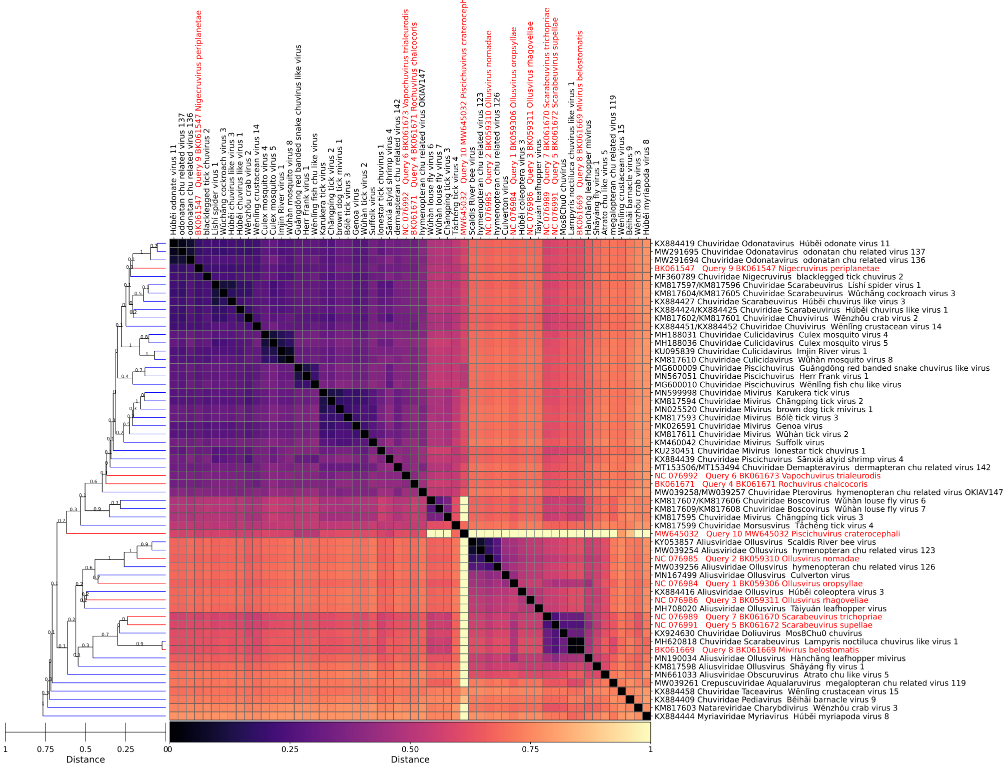


(a)

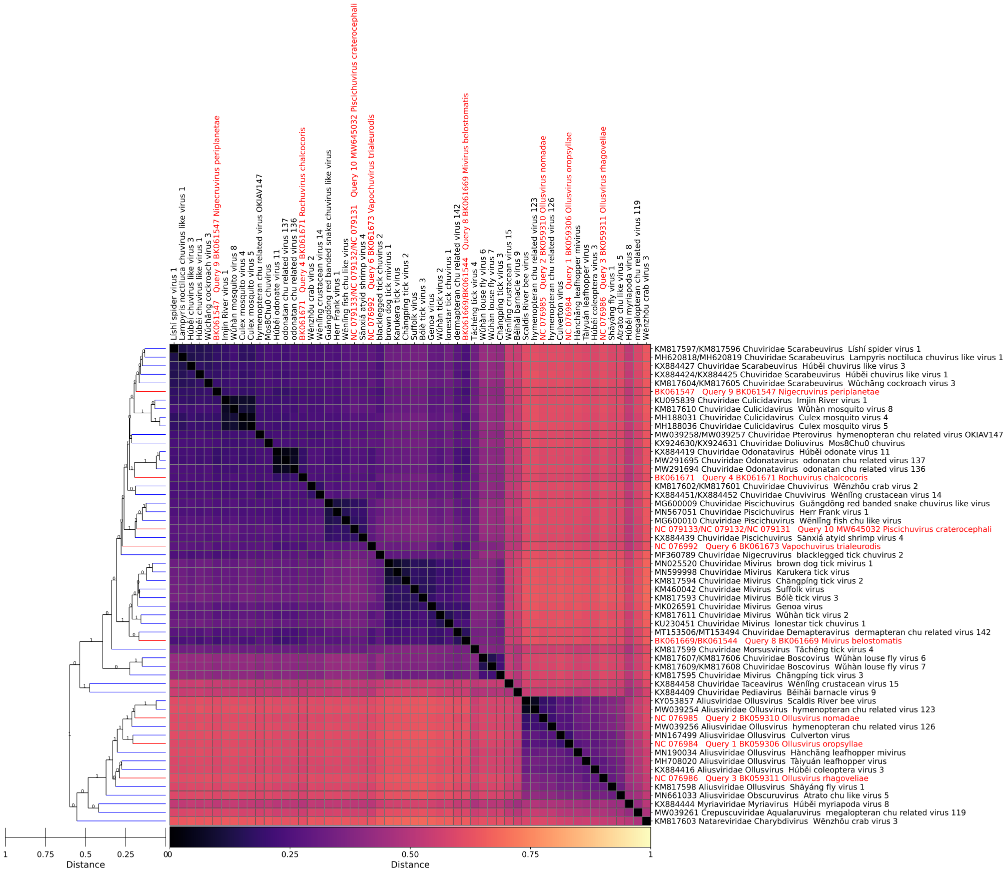


(b)

Figure 2: S2. **GRAViTy-V2 heatmaps, Jingchuvirales, showing effect of including coding incomplete sequences**. (a) Using sequences derived from TP, with 8 family violations and 2 genus violations due to including 6 incomplete sequences (BK061669, BK061670, BK061672, MW645032, MH620818 KX924630). (b) Corrected 4 sequences (BK061669, KX924630, MW645032, MH620818) and removed remaining 2 for which no replacements could be found (BK061670, BK061672), leaving a single genus violation (KM817595).

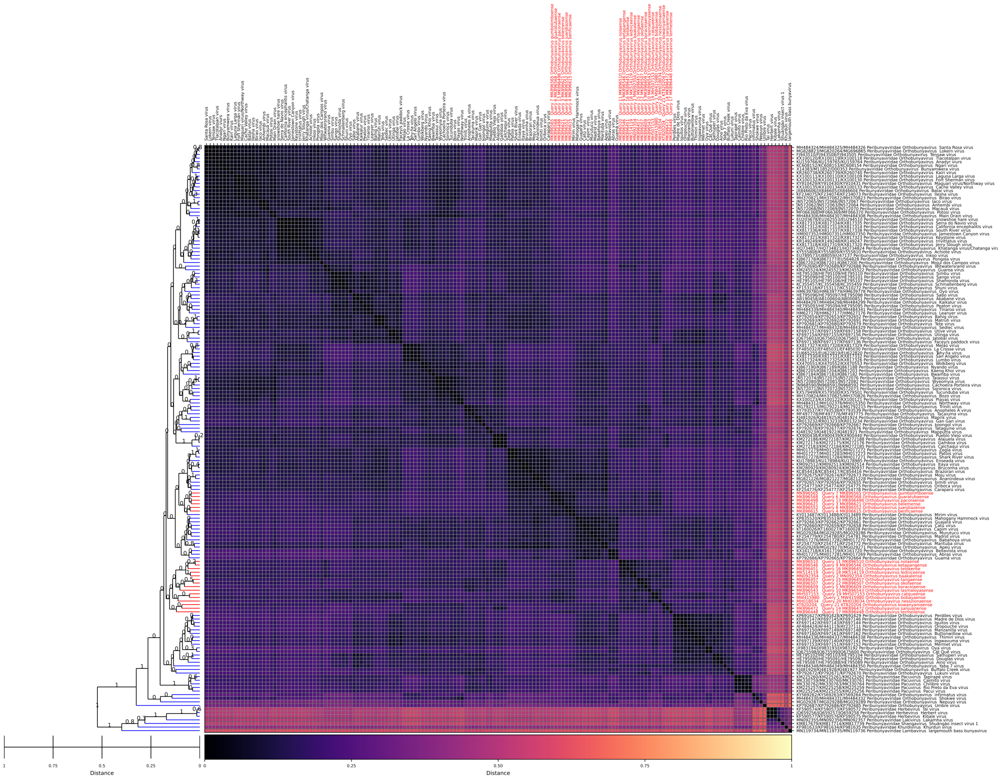

(a)

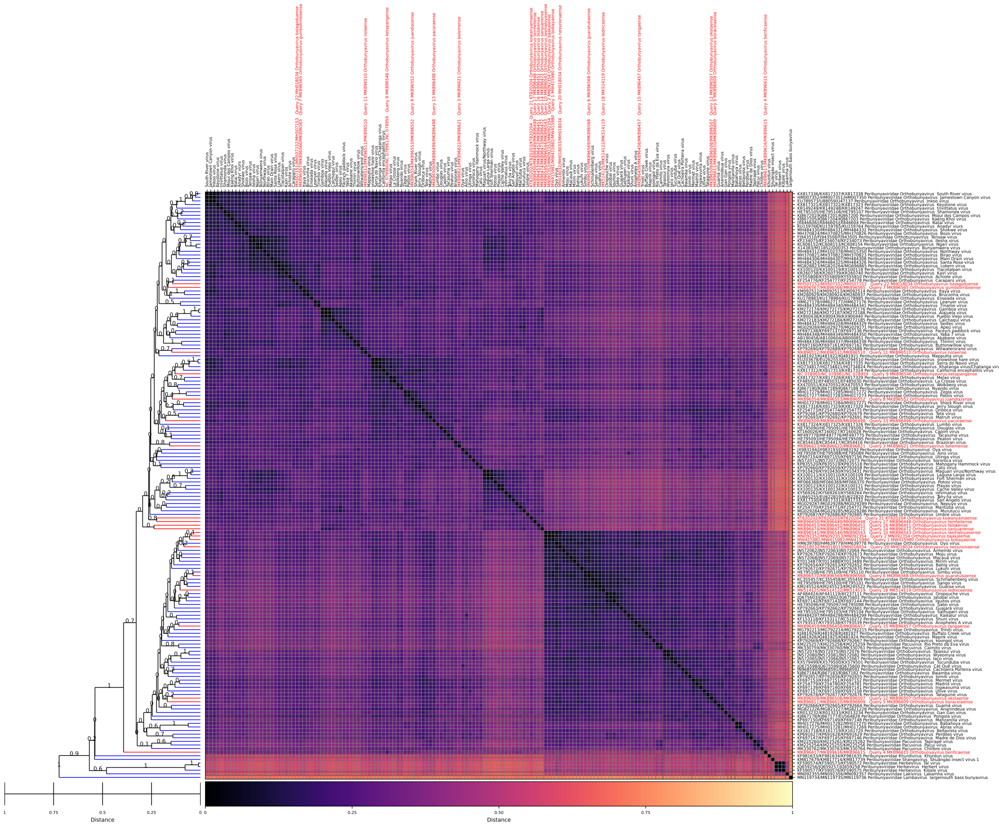

(b)

Figure 3: S3. **GRAViTy-V2 heatmaps, *Orthobunyavirus* (018M), showing effect of improperly assembled multipartite genomes**. (a) Input unclassified sequences (red) were provided pre-assembled rather than being assembled by GRAViTy-V2 from GenBank sequences. Assembly was inverse order to classified sequences, causing unclassified sequences to cluster together incorrectly. (b) As in a, with correct assembly of unclassified sequences.

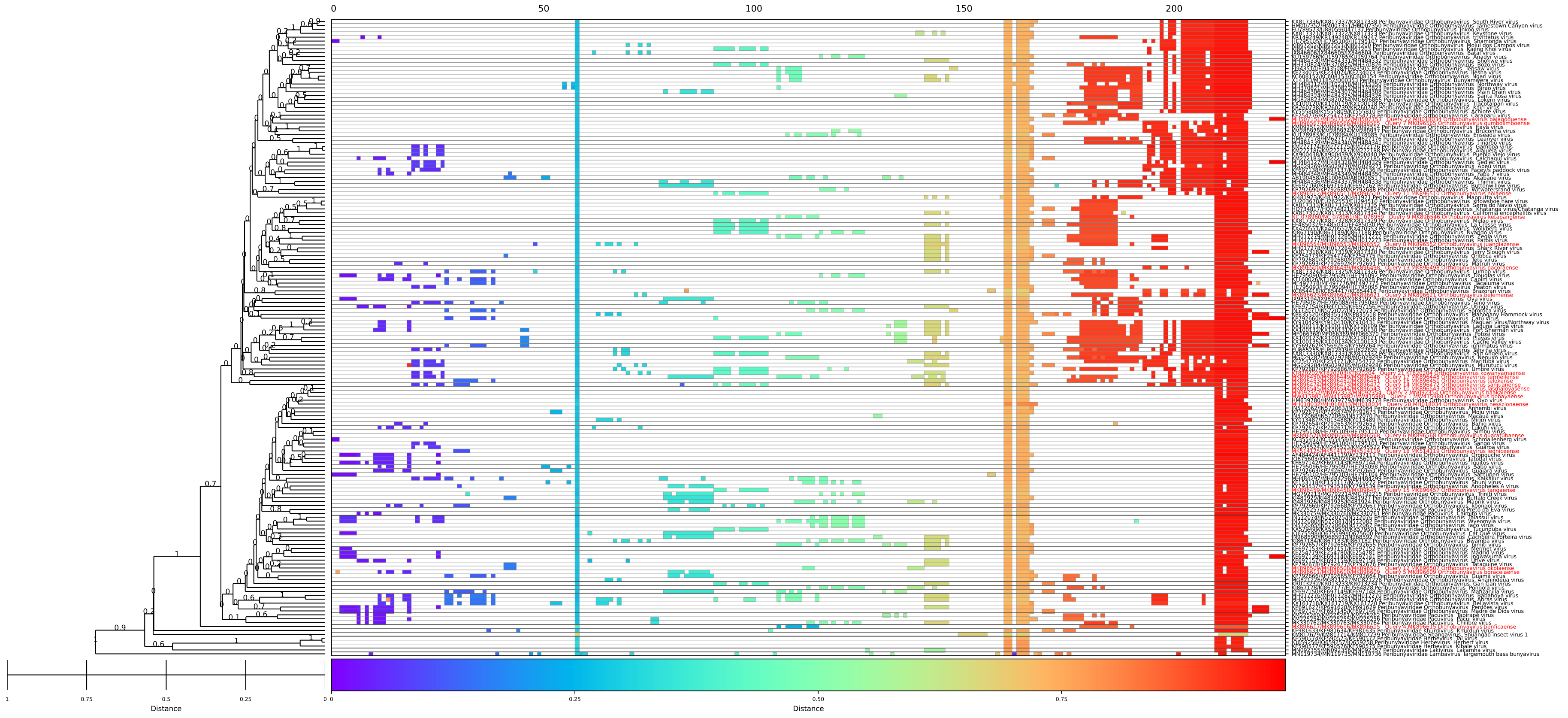


Figure 4: S4. **Barcode,** *Orthobunyavirus* (018M). Genus violation MK896615-7 (*Orthobunyavirus benficaense*) shows missing profile section in central (orange) profile block, corresponding to M segment which is non-coding. Corresponds to heatmap above (Fig. S3a).
